# High-dimensional profiling demonstrates complexity, tissue imprinting, and lineage-specific precursors within the mononuclear phagocyte compartment of the human intestine

**DOI:** 10.1101/2021.03.28.437379

**Authors:** Thomas M. Fenton, Line Wulff, Gareth-Rhys Jones, Julien Vandamme, Peter B. Jørgensen, Calum C. Bain, Julie Lee, Jose MG. Izarzugaza, Kirstine G. Belling, Gwo-Tzer Ho, Ole H. Nielsen, Lene B. Riis, Tune H. Pers, Henrik L. Jakobsen, Allan M. Mowat, Søren Brunak, William W. Agace

## Abstract

Mononuclear phagocytes (MNP), including macrophages and classical dendritic cells (cDC), are highly heterogeneous cells with distinct functions. Understanding MNP complexity in the intestinal lamina propria (LP), particularly in humans, has proved difficult due to the expression of overlapping phenotypic markers and the inability to isolate these cells without contamination from gut-associated lymphoid tissues (GALT). Here, we exploited our novel method for isolation of human GALT-free LP to carry out single-cell (sc)RNA-seq, CITE-seq and flow cytometry analysis of human ileal and colonic LP MNPs. As well as classical monocytes, non-classical monocytes, mature macrophage subsets, cDC1s, and cDC2s, we identified a CD1c^+^ cDC subset with features of both cDC2 and monocytes, which were transcriptionally similar to the recently described cDC3. While similar MNP subsets were present in both ileal and colonic LP, the proportions and transcriptional profiles of these populations differed between these sites and in diseased states, indicating local specialization and environmental imprinting. Using computational trajectory tools, we identified putative early committed pre-cDC subsets and developmental intermediates of mature cDC1, cDC2 and cDC3, as well as monocyte–to-macrophage trajectories. Collectively, our results provide novel insights into the heterogeneity and development of intestinal LP MNP and an important framework for studying the role of these populations in intestinal homeostasis and disease.

**One sentence summary:** Fenton and Wulff *et al*. use single-cell methods to explore the complexity of the mononuclear phagocyte compartment of the human intestinal lamina propria, identifying distinct dendritic cell and macrophage subsets, site-specific transcriptional signatures, and lineage-specific precursors.

## Introduction

The mononuclear phagocyte (MNP) family consists of conventional dendritic cells (cDC), classical monocytes, non-classical monocytes and macrophages, each of which play specific roles in immune responses, tissue homeostasis and inflammation^1–3^. Whereas cDC are the main cells involved in the induction and shaping of adaptive immune responses^4^, tissue-resident macrophages are primarily involved in maintaining local tissue homeostasis, defense against infection and tissue repair^5,6^. Recent studies have highlighted considerable heterogeneity amongst MNP^7–10^, and it is now evident that these cells develop unique functions depending on the tissue context and niches in which they reside^11,12^. Despite this, our understanding of MNP diversity within human tissues remains limited. A better understanding of MNP diversity is essential not only for understanding tissue specific immune processes, but also for the possibility of therapeutically targeting MNP subsets.

The intestine is continually exposed to food and microbial products that are essential for our health^13,14^. The intestinal immune system must respond appropriately to these products to maintain tissue homeostasis and at the same time retain the ability to mount effective immunity to intestinal pathogens. Furthermore, the intestine is not just a homogenous tube but consists of several anatomically and functionally distinct segments. For example, the small intestine, whose surface is characterized by finger-like projections termed villi, is the major site of food digestion and absorption. Conversely, the colonic surface consists of flattened crypts and it is home to the largest number and variety of microbes^15^. As a result, the concentration of dietary and microbial products and metabolites, many of which have direct impacts on local immune cell development and function, varies greatly along the length of the intestine. How such variations in intestinal anatomy, function, and luminal contents impact local MNP diversity in humans remains unclear.

Given its continual exposure to foreign material, it is unsurprising that the intestine contains the largest and most diverse immune compartments in the body. MNP are found in multiple niches throughout the intestine, including the intestinal lamina propria (LP), the muscularis mucosa and the gut-associated lymphoid tissues (GALT), including the multi-follicular Peyer’s Patches (PP) of the ileum and the mucosal- and submucosal-isolated lymphoid follicles (ILFs) that are distributed along the length of the intestine^16^. Much of our understanding of the roles of intestinal MNP diversity and function comes from studies in mice. These studies have not only highlighted the different roles that MNP subsets play in intestinal homeostasis, but also show that MNP composition and function is highly dependent on the intestinal niche in which they reside^1,3,12,17,18^. Consistent with this, recent single-cell transcriptomic analysis suggests considerable heterogeneity within the macrophage compartment of the human colonic mucosa and muscularis mucosa^19^. While these findings highlight the importance of assessing MNP diversity in different intestinal niches, this has not been possible in the human LP due to a lack of protocols to isolate LP tissue free from contaminating submucosa and GALT.

Here we used our recently developed techniques to isolate intestinal LP free from contaminating GALT and submucosa^20,21^ to assess the phenotypic, transcriptional and developmental diversity of MNP populations in the human ileum and colon LP. Our results provide novel insights into LP MNP diversity as well as site-specific adaptations within these populations, and provide an important framework for designing target approaches for modulating intestinal immune responses.

## Results

### MNP populations of the human ileum and colon LP are highly diverse

To assess MNP diversity within the intestinal LP, surgical samples of uninvolved ileum and colon from colorectal cancer patients (>10 cm from the tumour) were processed to remove contaminating GALT and submucosa (SM), as we described recently^20,21^. Following LP digestion, single-cell RNA sequencing (scRNA-seq) was performed on flow cytometry-sorted CD45^+^CD3^-^CD19^-^ HLADR^int/+^ cells from LP cell suspensions, using the 10x Chromium system (**Fig. 1A**). Sequences were obtained from six colonic LP and four paired ileal LP samples (**Table S1**). Distinct clusters of *CD3E*^+^ T cells, *CD79A*^+^ B cells, *VWF*^+^ endothelial cells, *MS4A2*^+^ mast cells, *COL3A1*^+^ stromal cells, and *NRXN1*^+^ glia were identified and excluded from further analysis (**Fig. S1A**). The MHCII genes were expressed by one ‘supercluster’ and two peripheral clusters (**Fig. S1B**), which were computationally isolated and re-clustered. These 28,758 MHCII^+^ cells comprised distinct clusters of *IL3RA*^+^ plasmacytoid DC (pDC), *CLEC9A*^+^ cDC1, and *FCGR3A^+^* non-classical monocytes (**Fig. 1B**), together with a supercluster that contained cells expressing either the cDC2-associated marker *CD1C,* the monocyte/macrophage (Mono/Mac)-associated marker *CD14*, or both *CD1C* and *CD14* (**Fig. 1C**). Flow cytometry analysis of colon LP CD45^+^ HLA-DR^+^ lineage^-^ cells confirmed the presence of CD1c and CD14 single positive cells, as well as cells expressing variable levels of both CD1c and CD14 (**Fig. 1D**).

**Fig. 1.**
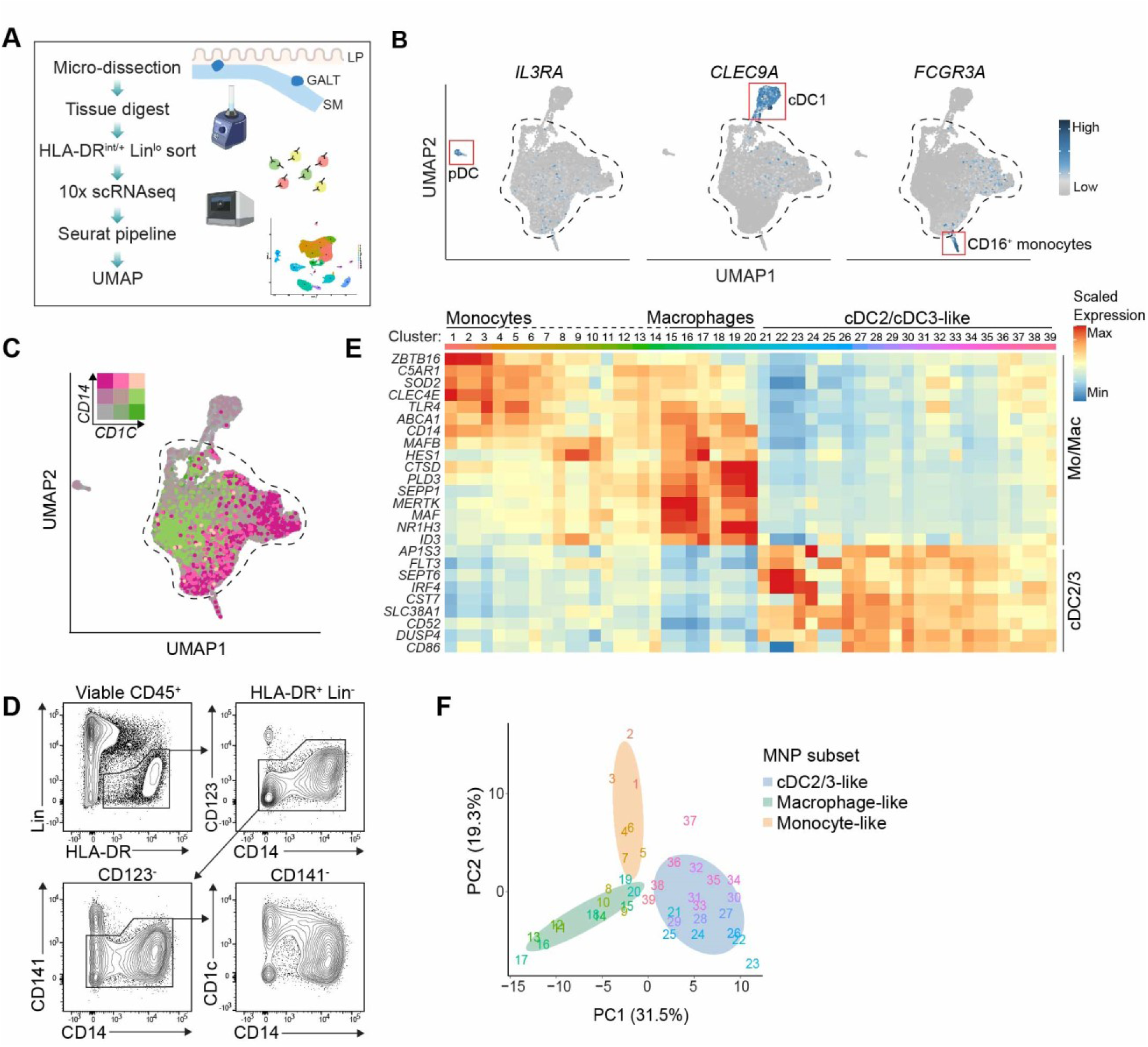
Using high-resolution clustering to disentangle MNP subsets of the human ileal and colonic LP. (**A**) Experimental pipeline for the generation of single-cell transcriptional data of intestinal LP MNP. (**B**) UMAP clustering of pooled ileal (n=4) and colonic LP (n=6) MNPs (28,758 cells), showing normalized gene expression of signature genes for pDC1 (*IL3RA*), cDC1 (*CLEC9A*) and non-classical monocytes (*FCGR3A*) signature gene. (**C**) UMAP of *CD1C* and *CD14* gene expression by intestinal LP MNP. Dashed line encompasses MNP which are not identified as pDC, cDC1, or non-classical monocytes. (**D**) Flow cytometry gating strategy showing CD1c and CD14 expression on colonic LP CD123^-^CD123^-^ MNP, representative of 10 non-IBD resection patients. (**E**) Curated pseudo-bulk heatmap (using averaged gene expression per cluster) of clusters within the dashed line of the UMAP in (**C**), showing expression of known monocyte, macrophage and cDC2/3 signature genes for clusters 1-39. ISG= interferon-stimulated genes, CCG= cell cycle genes. (**F**) Pseudo-bulk principal component analysis of clusters from **E** using signature gene lists from blood-derived cDC2, classical monocytes, and in-vitro generated monocyte-derived macrophages^106^.

To further characterize subsets within this MNP supercluster, we re-clustered these cells at high resolution (**Fig. 1E**). These clusters were present in all four ileal samples and six colonic LP samples, albeit in slightly different proportions (**Fig S1C**). To explore the identity of these clusters we used a supervised approach to assess individual clusters for their expression of known monocyte-, macrophage-, and cDC2-associated genes^7,22–25^. Based on this analysis, clusters were broadly separated into three main groups. The first clearly defined group (clusters 1-3) expressed the classical monocyte transcription factor *ZBTB16*^26^, while the second group (clusters 15-20) expressed high levels of macrophage-associated genes *SEPP1, MERTK* and *MAF,* as well as other TFs involved in tissue-resident macrophage development such as *ID3*^27^ (**Fig. 1E**). Clusters 4-14 expressed intermediate levels of these monocyte and macrophage signature genes. Thus, clusters 1-20 appeared to represent monocytes, macrophages and transitional intermediates between these cell types. Within the third group (clusters 21-39), clusters 21-35 expressed lower levels of monocyte and macrophage associated genes and high levels of the cDC2-associated genes *AP1S3*, *FLT3, SEPT6* and *IRF*^9,28^ (**Fig. 1E**), while clusters 36-39 expressed both cDC2-associated genes, including *FLT3* and *IRF4,* as well as monocyte and macrophage-associated genes, including *CD14* and *C5AR1* (CD88) (**Fig. 1E**). As a novel population of cDC3 has been reported to co-express cDC2 and monocyte markers^7,25,26,28–30^ (reviewed in^31^), we designated this third group as consisting of mixed cDC2 and cDC3-like cells.

To independently assess the accuracy of this grouping, we performed pseudo-bulk PCA analysis of the clusters using gene signatures taken from published cDC2, classical monocyte, and *in vitro* monocyte-derived macrophage data sets^32^. This analysis separated the clusters in a similar manner to our supervised approach, as PC1 drove separation of cDC2/DC3-like clusters from monocytes and macrophages and PC2 drove separation of monocyte from macrophage clusters (**Fig. 1F**). In summary, our analysis suggests that MHCII^+^ clusters within the human ileal and colonic LP consist of classical and non-classical monocytes, pDC, cDC1, macrophages, and cDC2/DC3-like cells.

### Identification of cDC1, cDC2 and cDC3-like subsets in the human intestine

To further explore cDC diversity within the intestinal LP, we computationally isolated and recombined the cDC1 and cDC2/DC3-like cells identified in Figure 1. Cells were clustered at high resolution to allow more accurate cluster designation, tSpace was used to perform trajectory-based clustering identification^33^ and the data was visualised by 2-dimensional representation of a 3-dimensional tSpace UMAP (Flat tUMAP). All sub-clusters were present in both ileal and colonic LP datasets (**Fig. S2A**). Seven clusters, located together at the top of the tUMAP, were enriched in cells expressing high levels of mitotic G2M/S genes (**Fig. S2B)** and relatively low levels of MHCII genes (**Fig. S2C**), both of which are characteristics of cDC precursors and will be considered later. To assess the identity of the remaining clusters, we first analysed expression of canonical cDC1 signature genes *CLEC9A, CADM1, XCR1, BATF3,* and *IRF8* and identified 7 clusters with a clear cDC1 signature score (**Fig. S2D**). We then ranked the remaining clusters based on their average module expression of cDC2- or cDC3-associated signature genes published by Bourdeley et al^30^ and generated with the AddModuleScore from Seurat (**Fig. S2E**). This allowed us to tentatively identify cDC2-like and cDC3-like clusters, as well as clusters that exhibited no particular bias in their cDC2/DC3 signature score, which we termed ambiguous clusters (**Fig. S2E**). We also identified a population of *LAMP3*^+^ cDC, which expressed high levels of the maturation markers *CCR7* and *CD40* (**Fig. S2F**), suggested to represent mature cDC with a migratory capacity towards lymph nodes^34,35^.

cDC1, cDC2-like and cDC3-like clusters grouped together into 3 super-clusters located in largely distinct areas of the tUMAP projection, with the ambiguous clusters positioned between the cDC2- and cDC3-like cells (**Fig. 2A**). Analysis of genes differentially expressed (DEG) between these subsets further supported their designation as cDC1, cDC2 and cDC3-like cells (**Fig. 2B** and **Table S2**). Specifically, the top DEG for the cDC1 cluster included *CLEC9A, CADM1* and *ID2,* the cDC2 cluster expressed high levels of *IRF4, PLAC8* and *CCL22*, the cDC3-like cluster expressed high levels of *C1QA, S100A9* and *CD163,* while the ambiguous cluster expressed genes associated with both cDC2 and cDC3-like cells (**Fig. 2B**). *CD1C, CLEC10A* and *FCER1A* were expressed at comparable levels by the ambiguous as well as the cDC2 and cDC3-like clusters in both ileal and colonic LP, as observed previously in blood cDC2 and cDC3^26,30^ (**Fig. 2B**). Gene Ontology (GO) terms differentially expressed between the cDC subsets included ‘Antigen processing and presentation of peptide antigen via MHC class I’ for cDC1, ‘positive regulation of T-helper cell differentiation’ for cDC2 and cDC3-like cells, and ‘cellular response to molecule of bacterial origin’ and ‘inflammatory response’ for cDC3-like cells (**Figure S2G**).

**Fig. 2.**
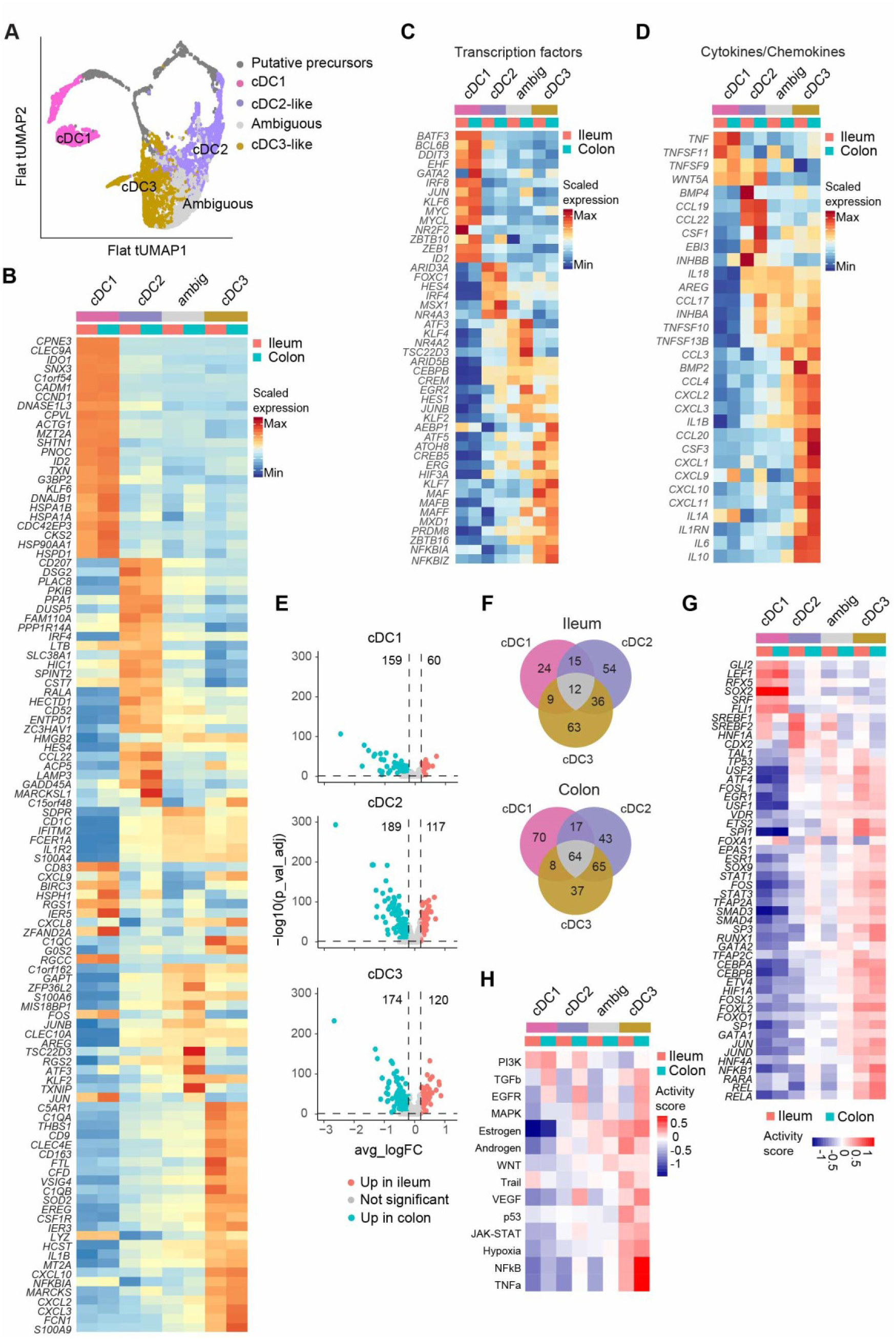
Transcriptional characterization of cDC subsets from ileal and colonic LP. (**A**) Flattened 3D tspace UMAP (tUMAP) plot of ileal and colonic LP cDC clusters grouped into indicated populations based on high-definition clustering and analysis in Fig. S2. (**B**) Pseudo-bulk heatmap of top 25 DEG expressed by mature cDC1, cDC2-like, cDC3-like and ambiguous cDC subsets of ileal (red) and colonic (blue) LP. (**C** and **D**) Manually curated pseudo-bulk heatmaps of differentially expressed (**C**) transcription factors and (**D**) cytokines and chemokines, between indicated cDC subsets. (**E**) Volcano plots of DEGs of indicated cDC subsets between pooled ileum and colon LP samples. Dashed lines indicate significance cut-offs. Adjusted P-values < 0.05 and |avg. logFC| > 0.2. (**F**) Venn diagrams depicting number of genes upregulated in indicated cDC subset in the ileum (upper) and colon (lower) LP, as well as genes commonly upregulated within these subsets. (**G**) DoRothEA based inferred transcription factor activity and (**H**) PROGENy based inferred signaling pathway activity in indicated cDC subsets and tissue.

To gain further insights into potential functional differences between cDC subsets, we manually curated a list of DEG between cDC subsets based on GO terms associated with transcription factors and cytokines/chemokines^36^ (**Fig. 2C and D)**. In addition to classical cDC1-associated TFs such as *IRF8*, *ID2* and *BATF3,* ileal and colonic cDC1 specifically expressed other TFs such as *ZEB1* (**Fig. 2C**), which was recently implicated in cDC1-mediated Th1 responses^37^. A number of TFs associated with development, including *ARID3A*, *FOXC1*, *HES4*, and *MSX1,* were enriched in cDC2-like cells, as well as *NR4A3,* which has been implicated in DC activation^38^. As expected, cDC3-like cells expressed the highest levels of macrophage-associated genes including *MAF, MAFB, MAF* and *ZBTB16*, as well as the inhibitory TFs *NFKBIA* and *NFKBIZ*, while cells within the ambiguous cluster expressed high levels of several TFs associated with activation, including *ATF3* and *JUNB* (**Fig. 2C**). The cDC also expressed various cytokines and chemokines in subset-specific patterns. Thus, both ileal and colonic LP cDC1 expressed high levels of the TNF family members *TNF* and *TNFSF11* (RANK-L), cDC2-like cells expressed high levels of *CCL19, CCL22* and *EBI3* and cDC3-like cells expressed a wide range of cytokines and chemokines, including *IL10*, *IL1B* and *IL6* and the interferon-inducible chemokines *CXCL9, CXCL10, and CXCL11* (**Fig 2D**); the latter, primarily expressed by a subset of cDC3-like cells (**Fig S2F**).

To assess potential transcriptional differences between ileal and colonic LP cDC, we performed DEG analysis between these sites for each of the cDC1, cDC2 and cDC3-like subsets. The transcriptional profile of each cDC subset differed between the ileum and colon LP (**Fig. 2E, Table S3**). While many DEG were cDC subset-specific, 64 genes were upregulated by all three cDC subsets in the colonic compared with the ileal LP and 12 genes were upregulated by all three cDC subsets in the ileal compared with the colonic LP (**Fig 2F**). To broadly assess differences between ileal and colon cDC subsets we performed GO analysis using enrichR (GO biological processes 2021). Pathways upregulated in all colonic cDC subsets included ‘regulation of cellular response to stress’ and ‘positive regulation of cytokine production’ (**Fig S2H**). Ileal cDC showed more subset-specific responses, with upregulation of genes involved in ‘protein targeting to ER’ (cDC1 and cDC2), ‘intestinal cholesterol absorptio’ (cDC2 and cDC3-like cells), and ‘regulation of complement activation’ (cDC1 and cDC3-like cells) (**Fig. S2H**).

To assess potential differences in TF and signalling pathway activity between ileal and colonic LP cDC subsets, we used the Discriminant Regulon Expression Analysis package (DoRothEA), which infers transcription factor activity from expression of downstream target genes^39^ (**Fig. 2G**), and the Pathway RespOnsive GENes package (PROGENy), which infers pathway activity in cells based on expression levels of pathway response genes^40^ (**Fig. 2H**). DoRothEA analysis suggested selective SOX2, FLI1, LEF1, and FOXA1 activity in cDC1 (**Fig 2G**), while the cDC3-like cells showed enhanced activity of a broad range of TF associated with cell activation including JUN, JUND, NFKB1, REL, RELA, STAT1 and STAT3 (**Fig. 2G**), consistent with their TF and cytokine/chemokine gene expression profiles (**Fig 2C** and **D**). DoRothEA analysis also suggested tissue-specific differences in cDC TF activity. For example, all ileal LP cDC subsets showed increased SREBF1 and SREBF2 activity, both of which are involved in sterol/cholesterol metabolism), as well as HNFA1 and CDX2 activity, both of which have been implicated in driving intestine-specific cell fate transcriptional programs^41,42^ (**Fig 2G**). Conversely, colonic cDC subsets showed highly specific activity of FOXA1 (**Fig 2G**), which has been implicated in intestinal epithelial cell fate decisions^43^. PROGENy analysis suggested that the PI3K pathway was particularly active in cDC1, while cDC3-like cells displayed a broad activation of the Estrogen, Androgen, WNT, TRAIL, VEGF, p53, JAK-STAT, hypoxia, NFκB, and TNFα pathways relative to the other cDC subsets (**Fig 2H**), consistent with gene expression and DoRothEA analysis (**Fig. 2C, D** and **G**). Again, differences were found between ileum and colon LP cDC, with, for example, TGFβ, EGFR and MAPK signalling appearing more active in colonic cDC compared with ileal cDC and TRAIL signalling appearing more active in ileal compared with colonic LP cDC1 and DC3-like cells (**Fig 2H**). Collectively, these results highlight the distinct transcriptional activities of human intestinal cDC subsets and the importance of the environment in regulating the transcriptional profile of these cells.

### Intestinal cDC subset composition differs along the length of the intestine and changes during inflammation

CITE-seq analysis demonstrated that cDC2-like, cDC3-like and ambiguous cDC populations could be distinguished from monocytes and macrophages based on their high expression of CD1c and low expression of CD14 (**Fig. 3A**). We next used LEGENDScreen^TM^ to identify surface markers that could subdivide intestinal CD1c^+^CD14^-^ MNP and thus potentially distinguish intestinal cDC2 from cDC3-like cells by flow cytometry. We observed that CD11a and CD207 separated both colonic and ileal LP CD1c^+^CD14^-^ MNP into 4 populations (**Fig. 3B**, for pre-gating see **Fig. S3A**). To determine the usefulness of these markers in enriching for cDC2 and cDC3-like cells, we stained colonic LP MNP with CITE-seq antibodies to CD11a and CD207, again identifying cells within the cDC super cluster that were CD207^-^CD11a^-^, CD207^+^CD11a^-^, CD207^+^CD11a^+^ or CD207^-^ CD11a^+^ (**Fig. 3C**). While the ambiguous cDC population distributed evenly between all 4 quadrants (**Fig. 3D**), cDC2-like cells were enriched in the CD207^+^ CD11a^-^ (Q1) gate, while the cDC3-like cells were enriched in the CD207^-^CD11a^+^ (Q4) gate (**Fig. 3C** and **D**). CITE-seq analysis of paired ileal and colonic LP samples from one patient showed similar enrichment of cDC2-like cells amongst Q1 cells and cDC3-like cells amongst Q4 cells in the ileum (**Fig. S3B** and **C**).

**Fig. 3.**
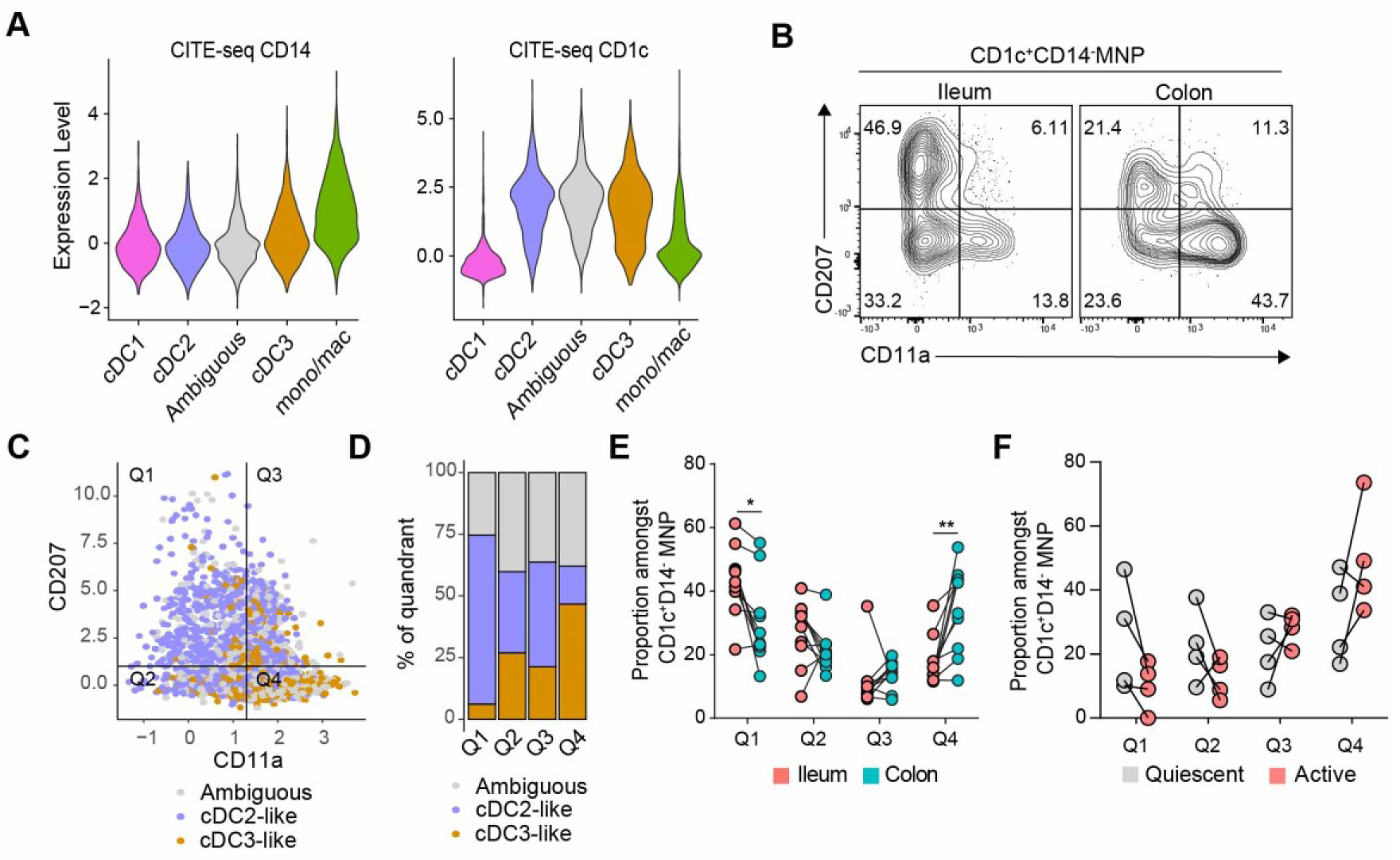
cDC2- and cDC3-like cells are found in different proportions in the ileal and colonic LP. (**A**) Violin plots of CD14 and CD1c surface expression by colonic MNP subsets after denoising and scaling to background (DSB)-normalization of CITE-seq. Results are pooled from three colonic and one ileal LP samples. (**B**) CD207 and CD11a surface expression on CD1c^+^CD14^-^ MNP from indicated tissues using flow cytometry. Results are representative of 10 ileal and colonic LP samples. (**C** and **D**) Pooled colonic LP samples from three CRC resection patients showing (**C**) CD207 and CD11a expression on colonic cDC2-like, cDC3-like and ambiguous cDC clusters using DSB-normalized CITE-seq, and (**D**) Proportion of each cDC cluster within each of the 4 quadrants (Q1-4) depicted in (**C**). (**E** and **F**) Proportion of CD207^+^CD11a^-^ (Q1), CD207^-^CD11a^-^ (Q2), CD207^+^CD11a^+^ (Q3) and CD207^-^CD11a^+^ (Q4) cells amongst CD1c^+^CD14^-^ MNP. Samples from **(E)** paired ileal and colonic LP from CRC resection patients (n=10), and (**F**) paired colonic biopsies taken from areas of quiescent or active inflammation from CD patients (n=4) as assessed by flow cytometry. Three to five biopsies were taken per site, and the inflammatory activity of each site was scored as quiescent or active by the endoscopist at time of removal. Statistical significance was determined using 2-way ANOVA with Sidak’s multiple comparisons, *p<0.05, **p<0.01.

To assess whether ileal and colon LP contained different proportions of these populations, flow cytometry analysis was performed on CD1c^+^CD14^-^ MNP from paired ileal and colonic resection samples (**Fig. 3E**). Ileal LP CD1c^+^CD14^-^ MNP were significantly enriched in CD207^+^CD11a^-^ (Q1) cDC2-like cells compared with the colonic LP, while colonic LP CD11c^+^CD14^-^MNP were enriched in CD207^-^CD11a^+^ (Q4) cDC3-like cells (**Fig. 3E**). To investigate whether the proportions of these populations may change during intestinal inflammation, we analysed biopsies from treatment-naïve patients undergoing endoscopic screening for inflammatory bowel disease (IBD) diagnosis who were subsequently diagnosed with Crohn’s disease (CD). This analysis suggested a trend towards reduced proportions of CD207^+^CD11a^-^ (Q1) cDC2-like cells and enhanced proportions of CD207^-^CD11a^+^ (Q4) cDC3- like cells in areas of active inflammation, although the differences were not significant (**Fig. 3F**).

### The human intestinal LP contains putative cDC1, cDC2 and cDC3 precursors

Recent studies have identified putative committed precursors of cDC1 (pre-cDC1), cDC2 (pre- cDC2), and more recently, cDC3 (pre-cDC3), as well as uncommitted pre-cDC precursors, in human bone marrow, blood and tonsils^25,26,29,30,44,45^. To explore whether cDC precursors might also be present in human intestine, we focused on the HLA-DR^low^ cDC (**Fig. S2C**), which, using high-resolution tSpace based clustering, consisted of 8 clusters (**Fig. 4A**). These cells were highly proliferative compared with the mature cDC (**Fig. 4B**) and expressed low levels of *ITGAX* (encoding CD11c) (**Fig. 4C**); features consistent with previous studies of pre-cDCs in mice^46,47^ and in humans^29,45^. Given that these proliferating clusters formed three distinct branches that aligned with cDC1, cDC2 and cDC3-like cells, we hypothesized that each branch potentially represented cDC subset-specific precursors.

**Fig. 4.**
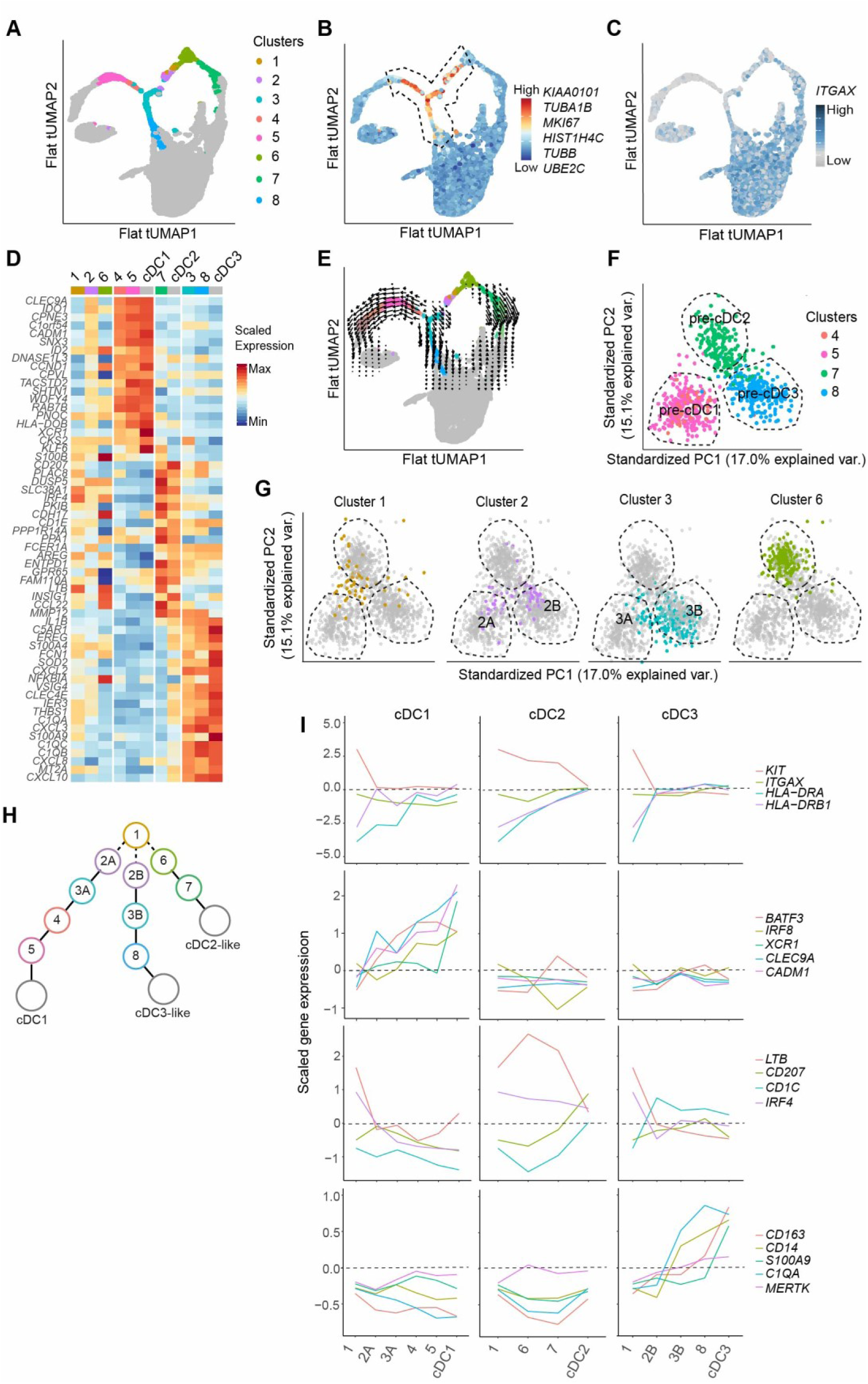
Identification of cDC committed precursors in the human intestine. (**A-C**) tUMAP as in Fig. 2A of ileum and colonic LP cDC clusters. (**A**) HLA-DR^low^ cDC clusters (cluster 1-8) overlaid onto cDC tUMAP, (**B**) proliferation score of indicated cell-cycle-associated genes and (**C**) *ITGAX* expression levels overlaid onto cDC tUMAP. (**D**) Heat map of top 20 DEG (calculated using p.adj. < 0.05) between cDC1, cDC2 and cDC3-like cells, showing expression levels in HLA-DR^low^ cDC clusters. (**E**) RNA velocities (arrows) of HLA-DR^low^ cDC clusters 3-5 and 7-8 calculated with Velocyto package embedded on (A). (**F**) PCA plot of cell from clusters identified by shared DEG (from **D**) as either pre-cDC1 (clusters 4 and 5), pre-cDC2 (cluster 7) or pre-cDC3 (cluster 8) and (**G**) location of clusters not identifiable in (D) (clusters 1-3 and 6) overlaid on the PCA plot in (**F**). (**H**) Model of precursor cluster trajectories towards mature cDC subsets based on tSPACE, velocity and transcriptional analysis. (**I**) Expression of indicated genes across proposed cDC1-, cDC2-, and cDC3- trajectories.

To assess this possibility, we generated signatures composed of the top 50 DEGs which distinguished the mature cDC subsets from each other and examined how these were expressed by the various clusters of HLA^low^ cDC. This analysis showed that cluster 4 and 5 shared a gene expression profile with cDC1, while cluster 7 expressed cDC2-like DEGs and cluster 3 and 8 had a similar gene expression pattern to DC3-like cells (**Fig. 4D**), suggesting the possibility that these clusters represented distinct cDC lineage specific precursors. RNA velocity analysis of mRNA splicing patterns^48^ further supported this idea, with cluster 4 appearing to be at the beginning of a trajectory with directionality into cluster 5, and thereafter into the mature cDC1 clusters (**Fig. 4E**). Similarly, cluster 7 showed a trajectory into the mature cDC2 clusters, while cluster 3 showed a trajectory towards cluster 8 and then into mature cDC3 (**Fig 4E**); similar patterns were observed in the ileum and colon LP (**Fig. S4A**). Collectively, these gene expression and splicing patterns suggest that clusters 4 and 5 represent pre-cDC1, while cluster 7 represents pre-cDC2 and clusters 3 and 8 represent pre-cDC3.

Three adjacent clusters (clusters 1, 2, and 6) did not express DEG specific to the mature cDC subsets (**Fig. 4D**) and we hypothesized that they may be earlier, less-committed precursors. To determine whether clusters 1, 2, and 6 showed evidence of commitment to any of the cDC lineages, we used the top 50 DEGs from each of the committed precursor clusters 5 (putative pre-cDC1), 7 (putative pre-cDC2), and 8 (putative pre-cDC3) as input for a PCA of all the HLA^low^ clusters. Of these total 150 DEGs between putative precursor clusters, 79 were also DEGs in the mature cDC populations. Committed precursor clusters 4, 5, 7 and 8 split into 3 distinct areas in PC1-2 (**Fig. 4F**). Using this approach, cluster 6 aligned clearly with pre-cDC2, while most of cluster 3 aligned, as expected, with pre-cDC3 (3B) and there were a few cells in the pre-cDC1 area (3A); one subset of cluster 2 (2A) aligned with pre-cDC1 and another subset of cluster 2 (2B) aligned with pre-cDC3 (2B) (**Fig. 4G** and **Fig. S4B)**). In contrast, cluster 1 did not overlap clearly with any of the pre-cDC groups (**Fig. 4G**).

To investigate the identity of the cells in cluster 1, we compared their gene expression profile with that of a recently published human bone marrow hematopoietic single-cell dataset^49^. Cluster 1 showed greatest correlation with hematopoietic stem cells (HSC), multipotent progenitors (MPP), lympho-myeloid precursors and early promyelocytes, together with some overlap with mature BM cDC. However they showed no overlap with late promyelocytes, myelocytes and classical monocytes (**Fig. S4C**). Thus cluster 1 appears to represent early lympho-myeloid progenitors with a potential bias towards the cDC lineage.

To further assess the relationship between mature cDC and their putative precursors we aligned clusters along the three putative cDC1, cDC2 and cDC3 developmental trajectories (**Fig. 4H**), and examined the expression of DC precursor and cDC subset associated genes across these trajectories (**Fig. 4I)**. Compared with other clusters, cluster 1 expressed the highest levels of *KIT*, similar levels of *ITGAX* and the lowest levels of MHCII gene (**Fig. 4I**), consistent with the suggestion that cells within this cluster represent early progenitors^25,26,29,44^. In agreement with the proposed trajectories, pre-cDC1 clusters progressively increased their expression of cDC1 related genes *BATF3*, *IRF8, CLEC9A* and *CADM1* as they transitioned through clusters 2A, 3A, 4 and 5 to mature cDC1. *XCR1* expression increased during the final transition from cluster 5 to mature cDC1 (**Fig. 4I**), consistent with recent studies in mice suggesting this is a late pre-cDC1 marker^50^. Expression of these genes remained low along the putative cDC2 and cDC3 trajectories. Conversely, expression of the cDC2 associated genes *IRF4* and *LTB* remained high across the cDC2 trajectory but was down-regulated along the cDC1 and cDC3 trajectories (**Fig. 4I**). *CD207* expression selectively increased along the cDC2 trajectory while *CD1C* expression increased along both cDC2 and cDC3 trajectories and decreased along the cDC1 trajectory (**Fig. 4I**). Finally, the putative pre-cDC3 clusters displayed a progressive increase in expression of the cDC3 associated genes *CD163, CD14, S100A9, C1QA* and *MERTK* as they transitioned through clusters 2B, 3B and 8 to mature cDC3-like cells (**Fig 4I**).

### Diversity of intestinal LP monocytes and macrophages

Studies of tissue macrophages in both humans and mice have highlighted significant niche-specific phenotypic, functional and ontogenic diversity^11,12,27,51,52^. Whether the human ileum and colon LP contains transcriptionally similar or distinct populations of intestinal macrophages remains unclear. Further, while most murine intestinal LP macrophages derive from monocytes along a ‘waterfall’ of phenotypic intermediates^53–56^, whether a similar monocyte ‘waterfall’ exists in the human intestine LP and the characteristics of such intermediates remains to be determined. To address these questions, ileal and colonic LP clusters that were identified as monocytes and macrophages (**Fig. 1E and F**) were analysed in isolation. tSpace principal components were used for trajectory-based clustering and UMAP embedding, resulting in the identification of 11 clusters (M1-M11) (**Fig. 5A**). DEG analysis of these trajectory-based clusters demonstrated that cluster M1 expressed high levels of the monocyte-associated genes, *S100A9*, *FCN1* and *VCAN*^25,57^; clusters M2 and M3 expressed intermediate levels of *S100A9*, *VCAN*, and *ITGAX* and low levels of *C1QC;* clusters M4-6 lacked expression of *S100A9* and *FCN1* and expressed intermediate levels of *ITGAX* and *C1QC;* and clusters M7 and M8 expressed high levels of *CD209* and *C1QC*, consistent with mature macrophages^57,58^ (**Fig. 5B and C**). The minor clusters M9 and M10 shared some features with mature macrophages, including high expression of *MHCII* and *C1Q* genes, but expressed low levels of *CD209* and *CD163* (**Fig. 5B and C**). Finally, the smallest cluster, M11, expressed high levels of cell-cycle associated genes including *MKI67* and *KIAA0101* (**Fig. 5B**), indicating that these represented a population of proliferating cells. Given the transcriptional identification of cluster M1 as monocytes, we used M1 as the starting point for pseudo-time analysis. Consistent with the transcriptional data (**Fig. 5B and C**), pseudo-time analysis suggested that clusters M2-M3 represented early, and clusters M4-M6, late transitional states, moving towards clusters M7-M10 (**Fig. 5D**). Using PROGENy analysis, the mature macrophage clusters M7-M9 showed evidence of responding to TGFβ (**Fig. S5A**), consistent with previous work in mice^58^. They also showed activation of the p53 pathway involved in multiple aspects of cell function, including cell-cycle arrest (**Fig. S5A**). Thus, we defined M2-M3 clusters and M4-M6 clusters as early and late intermediates, respectively, and clusters M7-M10 as mature macrophages (**Fig. 5B**). Analysis of single-cell data from paired ileal and colonic LP samples from four CRC patients showed that all clusters were present in both sites (**Fig. 5E**). There was also a trend towards increased proportions of intermediate clusters in the colon compared with the ileum LP, and increased proportions of more mature M7 and M8 macrophages in the ileal compared to the colonic LP (**Fig. 5E**).

**Fig. 5.**
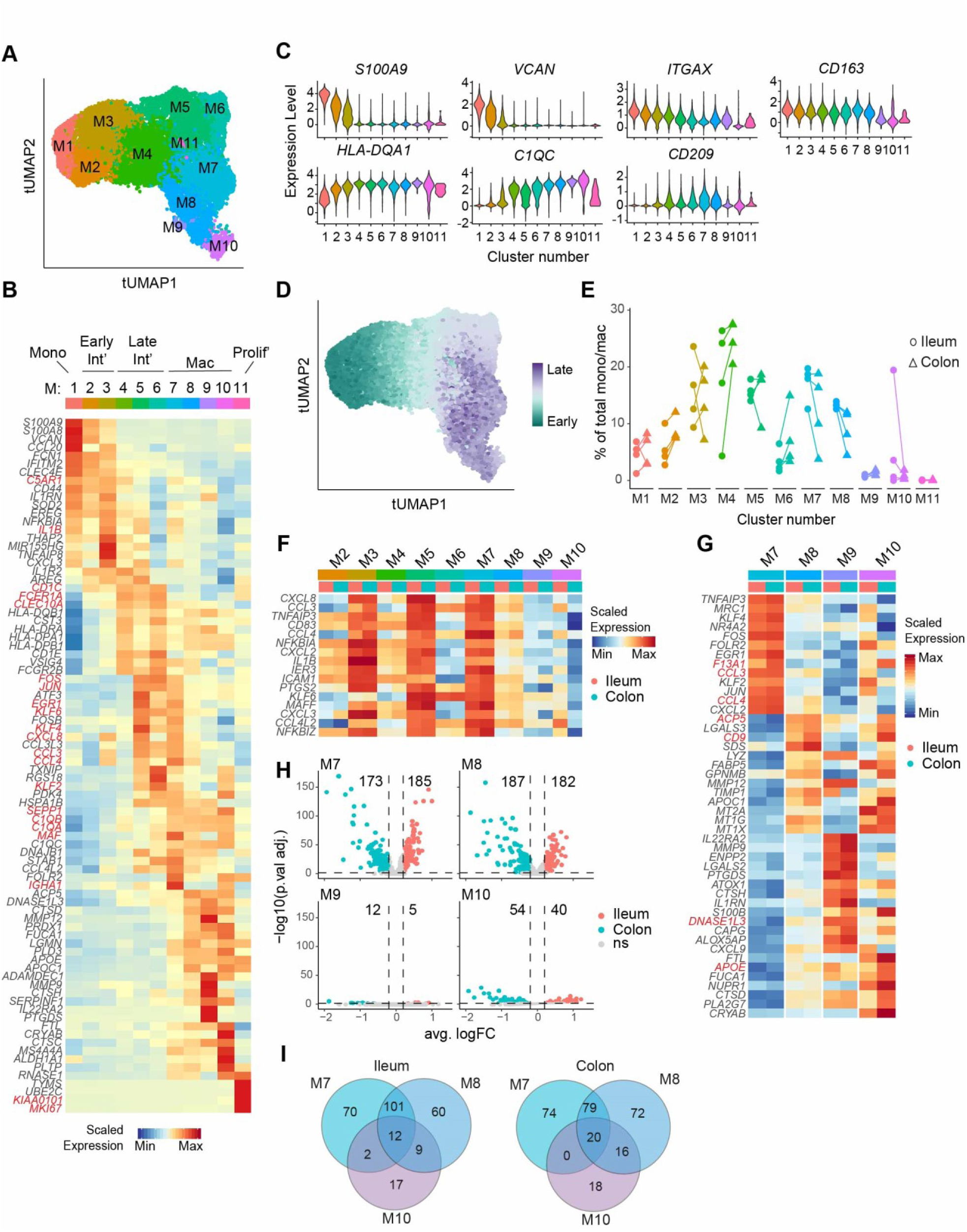
Characterization of intestinal LP macrophage populations. (**A**) tSpace UMAP (tUMAP) and Louvain clustering of ileal and colonic LP MNP identified as belonging to the monocyte-macrophage lineage. (**B**) Pseudo-bulk heatmap of scaled gene expression of top 10 DEG (ordered by avg. logFC) between M1-M11 clusters. (**C**) Violin plots of normalized gene expression of indicated maturation-associated genes in M1-M11 clusters. (**D**) Pseudotime of cells calculated by averaging all tSpace trajectories starting from M1. (**E**) Proportions of indicated clusters in paired ileal and colonic LP samples. (**F)** Pseudo-bulk heatmap of scaled gene expression of selected proinflammatory genes for clusters M1-M10 in the ileal and colonic LP. (**G**) Pseudo-bulk heatmap of scaled gene expression of top 13 DEG (ordered by avg. logFC) between mature macrophage clusters M7-M10 in ileal and colonic LP. (**H**) Volcano plots demonstrating DEGs between ileal and colonic LP in mature macrophage clusters M7-M10. Dashed lines indicating significance cut-offs. Adjusted P-values < 0.05 and |avg. logFC| > 0.2. (**I**) Venn diagrams showing overlap of ileum- and colon-specific DEGs for mature macrophage subsets M7, M8, and M10.

Analysis of clusters in both the ileal and colonic LP demonstrated that the early intermediate cluster M3, the late intermediate cluster M5 and the mature macrophage cluster M7 shared features of activation, including expression of the proinflammatory genes *IL1B, CXCL8*, *CCL3*, *CCL4,* and *NFKBIA* (**Fig. 5F**). Consistent with this, PROGENy analysis indicated enhanced TNFα and NFκB signalling (**Fig. S5A**), while DoRothEA analysis suggested enhanced canonical NFκB activation (NFKB1, RELA, REL, SP1 signalling), in particular in cluster M3, but extending to clusters M5 and M7, compared with other clusters (**Fig. S5B**). These clusters also showed evidence of enhanced EGFR and MAPK signalling (**Fig. S5A**), with M5 and M7 additionally showing activity of ELK1, a stimulus response transcription factor^59^, and activating transcription factor (ATF) 2 and ATF4^60^ (**Fig. S5B**). In contrast, clusters M2, M4 and M6 shared few common DEG distinct from clusters M3, M5 and M7, but these clusters did show evidence of enhanced WNT and JAK-STAT signalling (**Fig. S5A**), and activity of the transcriptional repressor RE1-Silencing Transcription factor (REST), and the MHCII promoter-associated regulatory factor X5 (RFX5) (**Fig. S5B**). Collectively, these results suggest that monocytes undergo a functional dichotomy in the intestine as they mature into tissue resident macrophages.

To gain further insights into the identity of the mature macrophage clusters M7-M10 we isolated these subsets and performed DEG analysis between them (**Fig. 5G, Table S4)**. The major mature macrophage cluster M7 expressed high levels of proinflammatory genes (**Fig. 5F and G, Table S4**), and genes associated with inhibition of NF-κB and PRR signaling including *NR4A2*^61^, *EGR1*^62^, and *KLF2/KLF4*^63^ (**Fig. 5G**). Consistent with their inflammatory profile, GO analysis demonstrated that M7 macrophages were enriched in several inflammatory pathways compared with other mature macrophage clusters, including ‘cellular response to molecule of bacterial origin’, ‘cellular response to cytokine stimulus’ and ‘regulation of inflammatory response’ **(Fig. S5C**). Amongst the mature macrophage clusters, M7 also expressed the highest levels of *LYVE1, SIGLEC1* (CD169)*, FOLR2,* and *MAF* (**Fig. S5D**), markers previously associated with perivascular macrophages^64,65^. While these genes have been associated with self-renewing embryonically derived MHCII^lo^ macrophages in other tissues^66^, the M7 cluster expressed high levels of *MHCII*, but not high levels of *ADAMDEC1* or *C2* (**Fig. 5C**) that have previously been associated with intestinal self-renewing macrophages^67^. Furthermore, while high levels of LYVE1 have been associated with intestinal submucosal macrophages^19,68^, the submucosa was removed during our LP isolation protocol, and cluster M7 did not express the submucosal macrophage associated genes *COLEC12*^19^ or *MARCO*^68^ (**Fig. S5D**). Thus, cluster M7 likely represents the recently identified *FOLR2* expressing macrophages present in the middle and base of intestinal crypts^68^.

The second major macrophage cluster M8, and the minor macrophage cluster M10, both expressed high levels of the metallothionein genes *MT1G*, *MT1X*, and *MT2A* (**Fig. 5G**), which have been implicated in metal sequestration in GM-CSF-dependent anti-microbial responses^69^. These clusters also expressed high levels of *LIPA, PLD3, LGALS3* and *CD68* (**Fig. 5G** and **Fig. S5D**), all of which have been associated with lipid metabolism and phagocytosis, and are signatures of lipid-associated macrophages^70,71^. Cluster M10 additionally expressed high levels of *APOE*, *PLA2G7*, and *FUCA1* (**Fig. 5G**), involved in lipid transport/metabolism, as well *FTL* involved in iron transport (**Fig. 5G**), both of which represent important functions of adipose tissue macrophages (ATMs)^72,73^. Consistent with this, cluster M10 showed evidence of TRAIL pathway activity (**Fig. S5A**), that has been associated with ATMs^74^, as well as activity of NR5A1, a transcription factor involved in steroidogenesis and lipid metabolism^75^ (**Fig. S5B**). Both clusters M8 and M10 were enriched in GO terms associated with responses to metal ions and ‘chylomicron remnant clearance’. Thus, macrophage clusters M8 and M10 showed transcriptional signatures indicative of lipid signaling, and cluster M10 in particular shared a transcriptional profile previously associated with ATMs.

Cells within the minor macrophage cluster M9 expressed high levels of the anti-inflammatory genes *ATOX1* and *IL1RN,* as well *IL22RA2,* the gene encoding IL22 binding protein (**Fig. 5G)**. They also expressed high levels of the metalloproteases *MMP9* and *MMP12* and the pro-angiogenic genes *ENPP2* and *PTGDS*, **(Fig. 5G**), all of which have been implicated in vascular homeostasis, and they showed the highest vascular endothelial growth factor (VEGF) pathway activity (**Fig S5A**), collectively indicating potential interactions with endothelium. M9 also expressed the highest levels of *CD4, C2*, and *ADAMDEC1* (**Fig. S5D**), genes associated with self-renewing macrophages in the mouse intestine^67^ and of *IL4I1* (**Fig. S5D**), recently identified as a marker of LP macrophages located at the top of colonic crypts and small intestinal villi^68^.

To determine whether the transcriptional profile of macrophage clusters M7-M10 differed between intestinal sites, we performed DEG analysis between the ileum and colon for each of these subsets. The major macrophage clusters M7 and M8 showed many DEG between the ileum and colon, while M10 and, even more so, M9, showed fewer differences (**Fig. 5H, Table S5**). GO analysis demonstrated that the M7 cluster within the ileum was enriched in genes associated with ‘antigen processing and presentation of exogenous peptide antigen’, ‘positive regulation of immune response’, ‘regulation of tumor necrosis factor production’, ‘positive regulation of reactive oxygen species metabolic process’, and ‘interferon-gamma-mediated signaling pathway’ compared with the colonic M7 cluster, and these pathways were similarly enriched in the ileal M8 compared to colonic M8 cluster **(Fig. S5E**). In contrast, the M7 cluster within the colon was enriched for genes associated with ‘receptor-mediated endocytosis’, ‘protein targeting to the ER’, ‘response to unfolded protein’, ‘regulation of cellular response to stress’, and ‘Fc-gamma receptor signaling pathway involved in phagocytosis’ and these pathways were also enriched in the colonic M8 compared with ileal M8 cluster **(Fig. S5E**). Collectively, these data suggest that the transcriptional profiles and functions of mature LP macrophages are influenced by the intestinal site in which they reside in a highly subset-specific manner.

### Flow cytometry-based identification of intestinal monocytes, intermediates and mature macrophage subsets

To identify surface antigens that may help identify the stages of intestinal monocyte development by flow cytometry, LEGENDScreen^TM^ was used to screen for surface marker expression on colonic CD14^+^CD1c^lo^ MNP (**Fig. S6A** and **B**). CD11c, CD11a, CD206, and CD55 showed heterogenous expression levels on CD14^+^CD1c^lo^ cells (**Fig. S6B**) and we thus used antibodies recognising these surface markers, together with CD14 and CD1c, in CITE-seq analysis of MNP from three colonic LP (**Fig. 6A**) and one ileal LP (**Fig. S6C**). CD55 was expressed at high levels by the monocyte cluster M1, at intermediate levels by the early intermediate clusters M2 and M3, but not by more mature clusters (**Fig. 6A**). CD11a was highly expressed by M1, M2 and M3 clusters, and at intermediate levels by the late intermediate clusters M4 and M5, but not by other clusters (**Fig. 6A**). CD206 was expressed by late intermediate clusters and by mature clusters M7 and M8, but not by the small M9 and M10 clusters. CD11c was expressed at high levels by cluster M1, all intermediate clusters and cluster M9, while mature M7 and M8 clusters showed heterogenous levels of expression and M10 little expression (**Fig. 6A**). CD14 was expressed at highest levels by cluster M1 and the early intermediate clusters M2 and M3 (**Fig. 6A**), while CD1c was poorly expressed by monocytes and macrophages, but a proportion of each of the intermediate clusters (M2-M6) expressed some CD1c (**Fig. 6A**). This analysis further suggested that the mature macrophage clusters could be broadly distinguished from monocytes and intermediates by their lack of expression of CD55 and CD11a and subsequently distinguished from one another based on their expression of CD14, CD206 and CD11c, with cluster M7 being CD14^+^CD206^+^, cluster M8 CD14^lo^CD206^int^, cluster M9 CD14^lo^CD206^-^CD11c^hi^ and cluster M10 CD14^lo^CD206^-^CD11c^-^ (**Fig. 6A**). The smallest M11 cluster contained too few cells for accurate CITE-seq analysis.

**Fig. 6.**
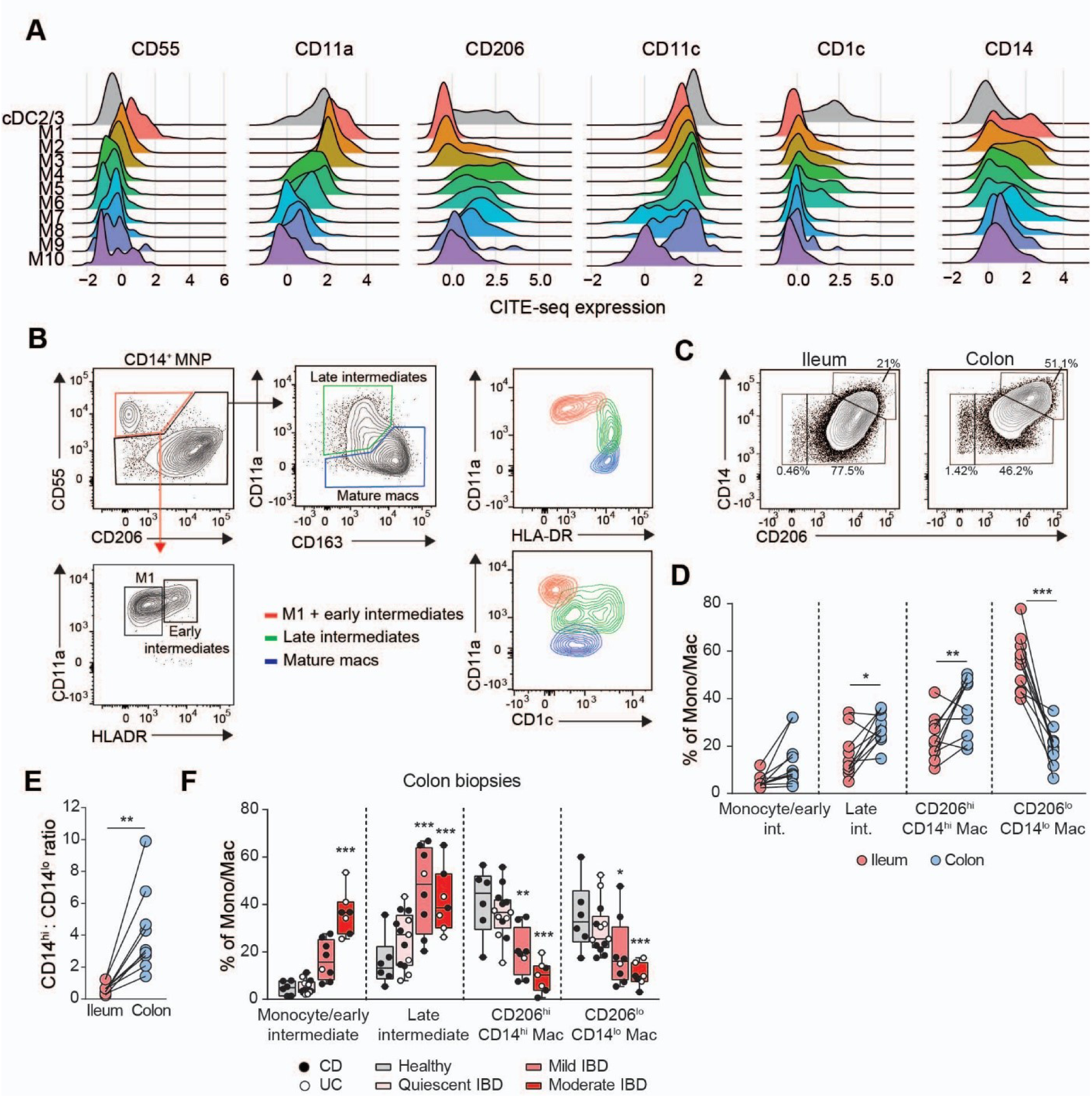
Flow cytometry analysis of mono/mac subsets. (**A**) DSB-normalized CITE-seq expression of indicated surface markers on pooled colonic LP macrophage clusters after exclusion of the minor proliferating M11 cluster, as well as total cDC2/3 clusters as control. Data are integrated from 3 independent colon samples. (**B-E**) Flow cytometry analysis of LP CD14^+^ mono/mac subsets obtained from digested CRC patient resection samples. (**B**) Colonic LP CD14^+^ MNP, showing gating strategy to identify putative mono/mac subsets, and surface expression of CD11a, HLA-DR and CD1c on each identified subset. CD14^+^ MNP were pre-gated as viable CD3^-^ CD19^-^ CD38^-^ CD123^-^ HLADR^+^ CD14^+^ singlets and data are representative of 10 patients. (**C**) Surface expression of CD14 vs. CD206 on ileal and colonic LP CD14^+^CD55^-^CD11a^int/low^ mature macrophages, showing 10 patients concatenated together. (**D**) Proportion of each mono/mac subset in paired ileal and colonic LP. Each symbol represents a paired ileal/colonic sample. Statistical significance was determined using 2-way ANOVA with Sidak’s multiple comparisons, *p<0.05, **p<0.01, ***p<0.001 (**E**) Ratio of CD14^hi^CD206^hi^ to CD14^lo^CD206^lo^ mature macrophage subsets within the ileal and colonic LP. Each symbol represents a paired ileal or colonic sample. Statistical significance was determined using Wilcoxon matched-pairs signed rank test, **p<0.01. (**F**) Proportion of monocyte/macrophage subsets based on flow cytometry analysis of digested colonic biopsies. Extent of IBD inflammation was scored at the time of biopsy by the clinician as quiescent, mild, or moderate. Each symbol represents a CD (filled circle) or UC (open circle) sample. Statistical significance was determined using 2-way ANOVA with Dunnett’s multiple comparisons comparing each IBD set to healthy controls, *p<0.05, **p<0.01, ***p<0.001

Based on these results and the transcriptional analysis depicted in **Fig. 5C**, we designed an antibody panel for putative identification of M1 and early intermediates, late intermediates, and mature macrophages by flow cytometry (**Fig. 6B**). Gating on mature macrophages, we identified three populations based on differential expression of CD14 and CD206 (**Fig. 6C**). These included CD14^hi^CD206^hi^ cells, which based on our CITE-seq analysis are likely enriched in M7 macrophages, while the CD14^lo^CD206^int^ cells are likely enriched in M8 macrophages, and the minor population of CD14^lo^CD206^lo^ cells is likely enriched in M9-M10 macrophages (**Fig. 6C**). To assess whether the proportions of these populations differed between the ileum and colon LP, we performed flow cytometry analysis on CD1c^-^CD14^+^MNP from 10 matched ileal and colonic resection samples (**Fig. 6D**). The proportion of intermediate cells among total mono/mac appeared higher in the colon compared with the ileal LP (**Fig. 6D**) and the colonic LP contained a higher proportion of CD14^hi^ CD206^hi^ macrophages and lower proportion of CD14^lo^CD206^lo^ macrophages compared within the ileal LP (**Fig. 6D and E**). To assess whether the proportions of these populations changed during inflammation, we performed similar analysis of colonic biopsies from healthy, CD and ulcerative colitis patients (**Fig. 6F**). This revealed a clear correlation between the presence of inflammation and increased proportions of early and late intermediates and decreased proportions of both mature CD14^hi^CD206^hi^ and CD14^lo^CD206^lo^ macrophage populations (**Fig. 6F**). Thus, multiple stages of monocyte-macrophage differentiation can be identified in the human intestine by flow cytometry, and these differ between the between the ileum and colon as well as in the setting of IBD.

## Discussion

MNPs play critical roles in tolerance, immunity and inflammation, but MNP are functionally heterogeneous and their subsets acquire distinct functions depending on the niche in which they reside. Characterizing MNP diversity in distinct human tissues is thus essential for our understanding of their various roles in tissue homeostasis and disease. Here, we used scRNA-seq, CITE-seq and flow cytometry analysis to characterize MNP diversity within the human intestinal mucosa, the largest barrier surface of the body. We provide evidence that the human ileum and colon contain numerous mature, transcriptionally distinct MNP subsets, as well as putative lineage-specific precursors. We further show that the proportion and transcriptional profile of MNP subsets changes along the length of the intestine and in the setting of intestinal inflammation. Our results provide an important roadmap of the human intestinal MNP compartment and a framework for future studies aimed at modulating this compartment therapeutically.

Consistent with previous findings^76,77^, pDC and non-classical monocytes represented only a minor fraction of cells within our intestinal LP MNP scRNA-seq datasets. As non-classical monocytes patrol the vasculature^78,79^, it may be that these represent contamination from the bloodstream, although there is also evidence that murine non-classical monocytes may migrate into tissues in response to tissue damage^78^, potentially including into the gut wall^80^. Consistent with previous immunohistochemical and flow cytometric analysis^81^, cDC1 were clearly identified as a distinct population in our LP MNP scRNA-seq datasets, but high-resolution clustering and a mixture of supervised and unsupervised approaches were required to distinguish the remaining MNP subsets from one another. This led to the identification of monocytes, monocyte-macrophage intermediates, mature macrophages and cDC2 subsets. In addition, there was a population of MNP that expressed both cDC2 and monocyte-associated genes, that were transcriptionally similar to the recently described population of cDC3 identified in several human tissues^7,25,26,30^. Compared with cDC2 and cDC1, these cDC3-like cells expressed a broad range of TFs associated with cell activation, together with a unique range of cytokines and chemokines, suggesting they play distinct roles in intestinal immune homeostasis.

Interestingly, our trajectory analysis suggested a degree of transcriptional convergence between cDC2 and cDC3-like cells within the intestinal LP, and we were unable to distinguish some of these cells at the transcriptional level. Consistent with this, we were also unsuccessful in identifying surface markers that could distinguish cDC2 from cDC3-like cells definitively. However, our CITE-seq analysis demonstrated that CD207^+^CD11a^-^CD1c^+^ cells were highly enriched in cDC2, while CD207^-^CD11a^+^CD1c^+^ cells were enriched in cDC3-like cells. Using these markers, we found that cDC3-like cells were present in higher proportions in the colon compared with the ileal LP, while cDC2 showed the opposite pattern. Furthermore, and consistent with previous studies in mice^82–84^, related cDC subsets displayed distinct transcriptional profiles and evidence of different signalling pathways when originating from the ileal or colonic LP. Thus, local environmental signals along the length of the human intestine may fine-tune cDC function.

Lineage-restricted cDC precursors have been identified in human blood, bone marrow and lymphoid tissues^25,29,44,45,85,86^, but it has been unclear whether such precursors exist in human non-lymphoid peripheral tissues, such as the intestine. Here, our combined bioinformatic analyses provide evidence that both the human ileal and colonic LP contain cDC precursors that appear committed to either the cDC1, cDC2 or cDC3 lineage. Thus, we found that each mature cDC subset was directly connected in trajectory space to a distinct population of proliferating *HLA*^low^*ITGAX*^low^ cells. Secondly, these distinct proliferating populations displayed a unidirectional velocity-based developmental trajectory into either mature cDC1, cDC2 or cDC3-like cells. Finally, these putative lineage-restricted precursors displayed a progressive acquisition or loss of cDC lineage-associated marker genes and TFs as they transitioned towards each mature cDC subset. As expected, the number of these putative lineage-restricted cDC precursors was low and our ability to capture such cells was only made possible by our sorting strategy and use of surgical resections as opposed to biopsies. Our evidence of lineage-restricted cDC precursors in the human intestinal LP is consistent with recent studies indicating the presence of cDC1 and cDC2 restricted cDC precursors in the murine small intestine^46^, and that human cDC3 derive from distinct precursors to those of cDC1 and cDC2^26,30^.

In addition to putative lineage-restricted cDC precursors, we also identified a minor population of proliferating *ITGAX* expressing *HLA*^low^ cells that did not show transcriptional bias towards any particular cDC lineage. The transcriptional profile of these cells instead correlated best with early bone marrow precursors, indicating that these cells may lie upstream of lineage-committed cDC precursors. While such findings are consistent with the observation that haematopoietic stem cells and/or downstream myeloid precursors are present in the human intestine^87^, the lineage potential and role these cells play in maintaining the intestinal MNP compartment awaits further study.

Prior scRNA-seq analyses of human intestinal macrophages have focused primarily on the colon and demonstrated macrophage diversity within both the mucosa and underlying mucosa muscularis^19,68^. Here, we confirm and extend these findings by demonstrating that, although the ileal and colonic LP contain similar mature macrophage populations, these are present in different proportions in the two tissues. Furthermore, similar to recent findings in mice^88^, we found that the transcriptional profile of the two major mature LP macrophage subsets, M7 and M8, differed between the ileal and colonic LP. Specifically, both populations in the ileum were enriched in pathways associated with immune activation and antigen presentation, while those in the colon were enriched in genes associated with response to unfolded protein and stress. Thus, similar to our observations with cDC subsets, the function of analogous macrophage subsets appears to be fine-tuned depending on their location along the length of the intestine.

Our trajectory-based bioinformatics analysis suggested that both major macrophage subsets, M7 and M8, derived from infiltrating monocytes, via transitional intermediates, a process reminiscent of the ‘monocyte waterfall’ described in mouse intestine^53,54,58,89^. Interestingly, a transcriptional dichotomy was observed within these transitional intermediates, whereby some clusters of early and late intermediates expressed high levels of pro-inflammatory cytokine and chemokine genes, as well as inferred activity of NFκB/TNFα/EGFR signalling pathways and NFκB/ATF/AP-1 transcription factors; these were characteristics also of mature M7 macrophages. In contrast, other early and late intermediate clusters showed inferred activity of JAK-STAT and WNT pathways, and of the transcriptional repressor REST, which were also characteristics of mature M8 macrophages. Collectively, these results suggest that monocytes transition along two transcriptionally distinct trajectories after their entry into the intestinal LP, resulting in the generation of the two major mature M7 and M8 macrophage subsets. Given the distinct location of mature macrophage subsets within the LP^68^, we speculate that these trajectories are driven by distinct signals transitioning cells receive within their local environmental niche.

We established a flow cytometry panel capable of identifying intestinal seeding monocytes, early and late intermediates as well as mature LP macrophage populations and found that the colon LP was enriched in late intermediate cells and CD206^hi^CD14^hi^ macrophages (M7 cluster), while the ileum LP was enriched in CD206^lo^CD14^lo^ macrophages (M8 cluster). Thus, in addition to exhibiting site-specific transcriptional profiles, the proportions of mature macrophages and monocyte intermediates differs between the ileum and colon LP. While the mechanisms driving these site-specific differences remain unclear, we speculate that the presence of high proportions of intermediate monocytes in the colon LP may reflect higher turnover of this compartment under steady-state conditions. Our flow cytometry analysis also confirmed and extended prior studies^76,90,91^, by showing that the proportions of both early and late intermediate cells increased in active IBD colon, while the proportions of both major mature macrophage populations decreased. Furthermore, these alterations correlated with disease severity. Whether intermediate cells accumulating in IBD are transcriptionally similar to those present in the healthy intestine, or acquire a distinct transcriptional profile as a result of local inflammatory cues, as recently suggested in mice^56^, requires further study.

In summary, our single-cell data highlight marked heterogeneity in the MNP compartment of the human intestinal LP, varying along the length of the human intestine and in the setting of disease. Additionally, by identifying novel transcriptomic and phenotypic markers, our work provides a road map for the study of MNP subsets, and their contribution to intestinal immune responses in health and disease.

## Materials and Methods

### Study Design

The main objectives of this study were to determine (a) MNP heterogeneity within the human intestinal lamina propria, (b) whether precursors to mature MNP subsets were present in the intestine and, (c) whether the proportions and transcriptional profiles of mature MNP subsets differed between human ileum and colon. Our hypothesis was that multi-modal single-cell methods (scRNA-seq, CITE-seq, flow cytometry) combined with bioinformatic analysis would allow for the unambiguous identification of MNP subsets within the intestine. Patient material included (a) surgical material from colorectal cancer patients (>10 cm from tumour site) and (b) biopsies from treatment-naïve patients undergoing endoscopy for suspected IBD. Patients below 18 and above 85 years of age were excluded from the study. Surgical samples were only used when it was possible to readily dissect mucosa from submucosa. Each experiment was replicated in at least three patients unless otherwise specified. No sampling replication was performed within an individual patient due to limited tissue availability.

### Methods

#### HUMAN SUBJECTS

Resection samples were obtained from patients undergoing surgery for colorectal cancer after informed consent with ethical approval from the Videnskabsetiske Komité for Region Hovedstaden, Denmark (H-3-2013-118). Biopsy samples were obtained from adult patients attending routine colonoscopy for initial IBD disease surveillance (both CD and UC) or for ongoing disease assessment (see **Table S6** for anonymized patient information) at the Western General Hospital, Edinburgh, UK, after informed consent under existing approvals (REC:19/ES/0087). All patients were part of the Lothian IBD registry^92^ and a diagnosis of IBD was made using the Lennard-Jones criteria^93^. Endoscopic assessment of disease severity at the biopsy site was made at the time of endoscopy and biopsy sites were classified as quiescent, mild, moderate or severe. Three to five biopsies were taken per site and pooled for analysis.

#### METHOD DETAILS

##### Tissue processing

Surgical samples were processed as described previously^21^. Briefly, resection samples were taken at least 10 cm distant from the tumor site. Muscularis externa was removed using curved surgical scissors and the remaining tissue was incubated in RPMI-5 (RPMI/5% FCS/1% penicillin and streptomycin) containing 4 mM DTT for 2 x 10 min at 37°C on a shaking incubator (370 rpm) to remove mucus. Macroscopically visible submucosa (SM) was trimmed away using scissors and mucosa separated from SM under a stereo microscope using forceps. Epithelial cells were removed by incubating the mucosa in Ca2^+^ and Mg2^+^ - free HBSS containing 1% penicillin and streptomycin and 5 mM EDTA at 37°C for 10 min in a shaking incubator, and this procedure repeated four times. Isolated lymphoid follicles were dissected from the mucosa using a scalpel under a stereo microscope with a transmitted light source, and remaining GALT-free LP was cut into 2-4 mm^2^ pieces in preparation for digestion. LP was incubated in RPMI-5 containing DNase (30 µg/ml) and collagenase D (5 mg/ml or Liberase TM (2.5 mg/ml) for 45 min at 37°C under gentle shaking (370 rpm). The resulting LP cell suspension was passed through a 100 µm filter and washed twice in fresh RPMI for downstream analysis. Biopsy samples were processed using the same protocol.

##### Flow cytometry, MNP enrichment and cell sorting

Cell suspensions were stained with indicated antibodies in Brilliant stain buffer (BD Biosciences) containing 4% normal mouse serum, according to standard procedures, with dead cells identified by 7-AAD staining and excluded from analysis. Samples were analyzed on an LSR Fortessa 2 (BD Biosciences) using Flowjo software (Treestar). The Legendscreen assay (Biolegend) was performed as per the manufacturer’s instructions. For scRNA-seq experiments, LP cell suspensions were enriched for HLA-DR^+^ cells using anti-HLA-DR microbeads (Miltenyi Biotec) and LS MACS columns according to manufacturer’s instructions. Resulting cells were stained with the indicated antibodies (**Table S7)** and 7-AAD was added before sorting to exclude dead cells. Cells were sorted on a FACSMelody (BD) into MACS buffer and re-suspended in PBS containing BSA (0.4%) for subsequent 10x analysis. For some samples, cells were first stained with barcode-labelled antibodies prior to sorting for CITE-seq analysis (**Table S7**).

##### 10x Chromium and sequencing

Sorted single cells were subjected to droplet-based massively parallel scRNA-seq using the *Chromium Single Cell 3′ Reagent Kit v3.1 with Feature Barcoding technology for Cell Surface Protein* (10x Genomics) following the manufacturer’s instructions. In short, the 10x Chromium Controller generated nanoliter-scale Gel Bead-In Emulsions (GEMs) droplets, where each cell was labeled with a specific barcode and each transcript labeled with a unique molecular identifier (UMI). After reverse transcription, GEMs were broken down and the barcoded cDNA purified with Dynabeads MyOne Silane beads (Thermofisher). cDNA was amplified by PCR using the supplied 10x genomics primers and protocols (12 amplification cycles). Products were size separated with SPRIselect beads (Beckman Coulter) to obtain larger purified transcribed mRNA products for gene expression libraries as well as smaller cell surface protein products for CITE-seq library generation. To prepare gene expression libraries for sequencing, purified large size cDNA products were processed and ligated to an index for sample pooling before the sequencing process following the manufacturer’s instructions. For CITE-seq analysis, 5 µl of the purified smaller cDNA product was used as template for antibody-derived tag (ADT) sequencing and ADT sequencing libraries were constructed and indexed following the manufacturer’s instructions. Quality and quantity of the final libraries were measured using the Agilent 2100 Bioanalyzer equipped with High Sensitivity DNA chip (Agilent). Illumina sequencing was carried out at the Genomics Core Unit, Center of Excellence for Fluorescent Bioanalytics (University of Regensburg, Germany) or at the SNP&SEQ Technology Platform, Sweden. Libraries were sequenced using HiSeq, NextSeq and NovaSeq systems (300 cycles), aiming for a minimum of 30,000 read pairs/cell for sc-RNA and 3000 read pairs/cell for ADTs.

##### Bioinformatic analysis

###### Data processing

Sequencing data was pre-processed and aligned with CellRanger (version 2.2.0 for the first patient samples and version 3.1.0 for all other samples)^94,95^. Sequencing data from the samples stained with TotalSeq antibodies was processed with CITE-seq count^96^. Each sample was read into a Seurat (version 3.1.5)^97^ object in R (versions 3.5.1/4.0.1)^98^ and processed by removing cells with exceptionally low or high UMI, gene counts (< 500-1000 and > 3000-6000 genes/cell) and mitochondrial gene content (>10%) according to current best practice^99^ and likely representing debris and doublets. To normalize CITE-seq data by denoising and scaling protein levels against background (DSB-normalization), the debris removed from each sample was used as empty droplet information (free floating CITE-seq antibody), while isotype controls were used to normalize for non-specific binding^100^. The normalized protein data were incorporated with the RNA data for the individual samples by adding to it to the corresponding Seurat objects.

After log-normalization of RNA levels for individual samples, cell cycle gene modules were calculated using the Seurat CellCycleScoring function and a gene module representing cell cycle genes from the tool ccRemover by summing the raw counts of these genes/cell divided by the total number of reads per cell^101^. Then the top 3000 most variable genes were identified per sample with the selection method vst.

After initial data processing, all the samples were integrated with Seurat anchor integration and gene expression was scaled while regressing out the effect of cell cycle, UMI counts, and mitochondrial gene content based on their scoring on the individual samples. The merged data were dimensionality reduced with PCA and the 15 first PCs were chosen for downstream analysis. A shared nearest neighbor graph was constructed and used to cluster the data with Louvain clustering. The PCs were also used to dimensionally reduce the data further with UMAP for visualization purposes.

###### Differential gene expression and gene ontology

Differential gene expression was calculated with Seurat FindMarkers for comparisons between specific groups or FindAllMarkers for DEGs for all clusters, both using the standard non-parametric Wilcoxon rank sum test and base=exp(1).

Gene ontology analysis was performed based DEG lists of above and run on EnrichR’s web interface. The output tables based on GO Biological Processes 2021 were downloaded and plotted in R using the package ggradar.

###### Pseudo-bulk analysis and comparison with populations in public databases

Pseudo-bulk analysis for heatmaps and PCA was performed with the Seurat AverageExpression function. Module scores were calculated with Seurat’s AddModuleScore function on indicated selected gene sets or gene sets from literature. DoRothEA was run using confidence levels A+B and referring to the human DoRothEA transcription factor interaction database^39^. Progeny was run using organism = human and top = 500 genes^40^.

The processed Seurat object from Triana *et al*^49^. was downloaded and subsetted on the clusters labelled “HSCs & MPPs”, “Lymphomyeloid prog”, “Early promyelocytes”, “Conventional dendritic cell 1”, “Conventional dendritic cell 2”, “Late promyelocytes”, “Myelocytes”, “Classical Monocytes”. The gene expression data was averaged, and Pearson correlations calculated based on variable genes from our data also present in the bone marrow data set.

###### Trajectory inference

Trajectory inference was performed with tSPACE on PC spaces of indicated populations^33^. The outputs were dimensionality reduced with UMAP^102,103^ from tPCs 1-15 (for both cDCs and macrophages), to 2 and 3 dimensional tUMAPs with distance metric set to Pearson. 3D tUMAP for the cDC trajectory was angled and embedded into 2D using the Dufy package^104^. Clustering was performed with Louvain clustering for Seurat on the tPCs as input. Pseudotime (arbitrary time scale unit) was calculated by taking all trajectories from the tSPACE output from M1 and averaging these per cell. Splicing patterns were first determined on individual samples with the advanced run setting for velocyto^48^ with a repeated annotation file^105^. Genes were filtered by 0.2 for spliced data and 0.05 for unspliced data. RNA velocity estimates were then calculated for T=1 and only included genes with splicing information also present in the variable genes and only on cells of interest (e.g. precursors). The information was embedded on top of 2D tUMAP using n=400, scale=sqrt, grid.n=50 and arrow.scale=2.

### QUANTIFICATION AND STATISTICAL ANALYSIS

Statistical analysis of flow cytometry data was performed using Prism software (GraphPad). Statistical analysis of sequencing data was performed in R. Statistical tests used for experimental data are outlined in the figure legends.

### DATA AND CODE AVAILABILITY

scRNA-seq count data is available through CZ CELLxGENE: https://cellxgene.cziscience.com/e/bcdec5fa-a7fa-4806-92bc-0cd02f40242f.cxg/. Code is available: https://github.com/LineWulff/FentonWulff_LP_MNP

## Acknowledgments

We thank all patients and staff at Herlev hospital, in particular the Gastroenterology Team (Department of Pathology) for help in providing tissue samples. Sequencing was performed at the National Genomics Infrastructure (NGI) and Science for Life Laboratory SNP&SEQ Technology Platform in Uppsala (supported by the Swedish Research Council and the Knut and Alice Wallenberg Foundation). We thank Dr J Rizk (Copenhagen University) for valuable discussions regarding signaling pathways. This work was supported by grants awarded to W.W.A from the Danish Research Council (Sapere Aude III 1331-00136B), the Swedish Medical Research Council (2017-02072), the Swedish Cancerfonden (18 0598), the Gut Cell Atlas, an initiative funded by the Leona M. and Harry B. Helmsley Charitable Trust, US and a grant from the Lundbeck foundation (R155-2014-4184), Denmark, to W.W.A and S.B

## Author contributions

The study was designed by T.M.F., L.W., A.M.M., and W.W.A. Tissue samples were provided by H.L.J., L.B.R., T.H.P., O.H.N., G-R.J., and G-T.H. Experiments were performed by T.M.F., L.W., G-R.J., P.B.J., J.L., and J.V. Bioinformatic analysis was performed by L.W., K.G.B., and T.M.F. and supervised by S.B., K.G.B., and W.W.A. The manuscript was written by T.M.F., L.W., and W.W.A and reviewed and edited by A.M.M. and C.C.B. after input from all authors.

## Data and materials availability

Accession numbers:

## List of Supplementary Materials

Supplementary materials includes 6 supplementary figures and 7 supplementary tables

**Table S1.**
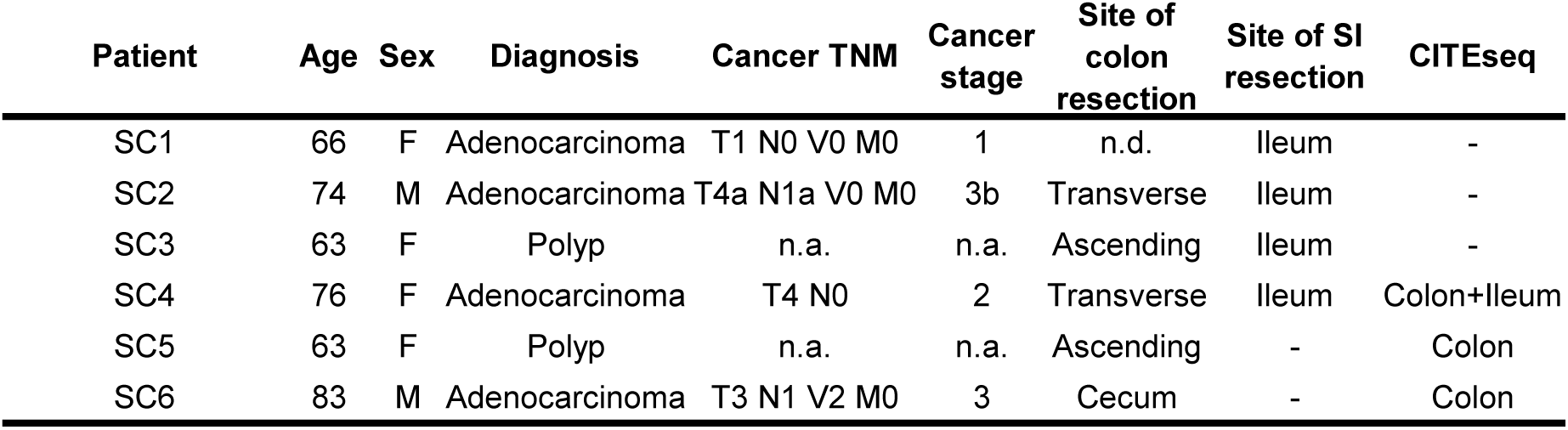
Characteristics of patient samples used for scRNAseq.

| Patient | Age | Sex | Diagnosis | Cancer TNM | Cancer stage | Site of colon resection | Site of SI resection | CITEseq |
| --- | --- | --- | --- | --- | --- | --- | --- | --- |
| SC1 | 66 | F | Adenocarcinoma | T1 N0 V0 M0 | 1 | n.d. | Ileum | - |
| SC2 | 74 | M | Adenocarcinoma | T4a N1a V0 M0 | 3b | Transverse | Ileum | - |
| SC3 | 63 | F | Polyp | n.a. | n.a. | Ascending | Ileum | - |
| SC4 | 76 | F | Adenocarcinoma | T4 N0 | 2 | Transverse | Ileum | Colon+Ileum |
| SC5 | 63 | F | Polyp | n.a. | n.a. | Ascending | - | Colon |
| SC6 | 83 | M | Adenocarcinoma | T3 N1 V2 M0 | 3 | Cecum | - | Colon |

**Table S2.**
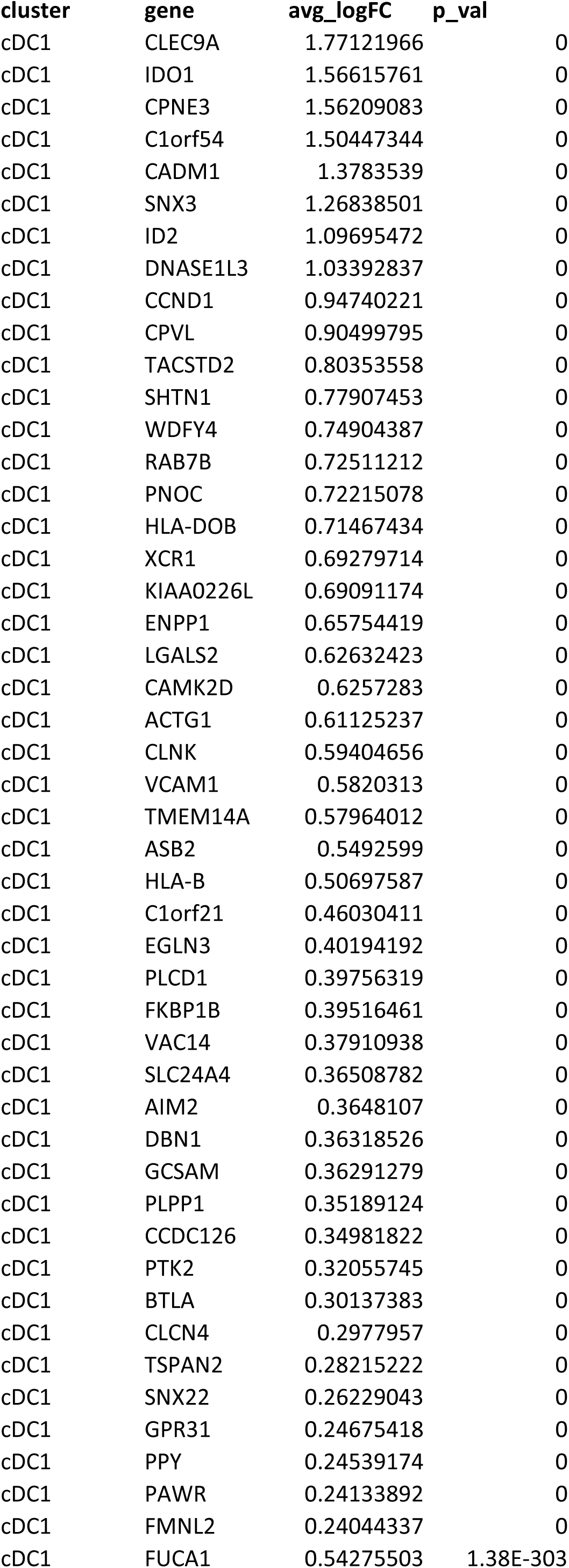
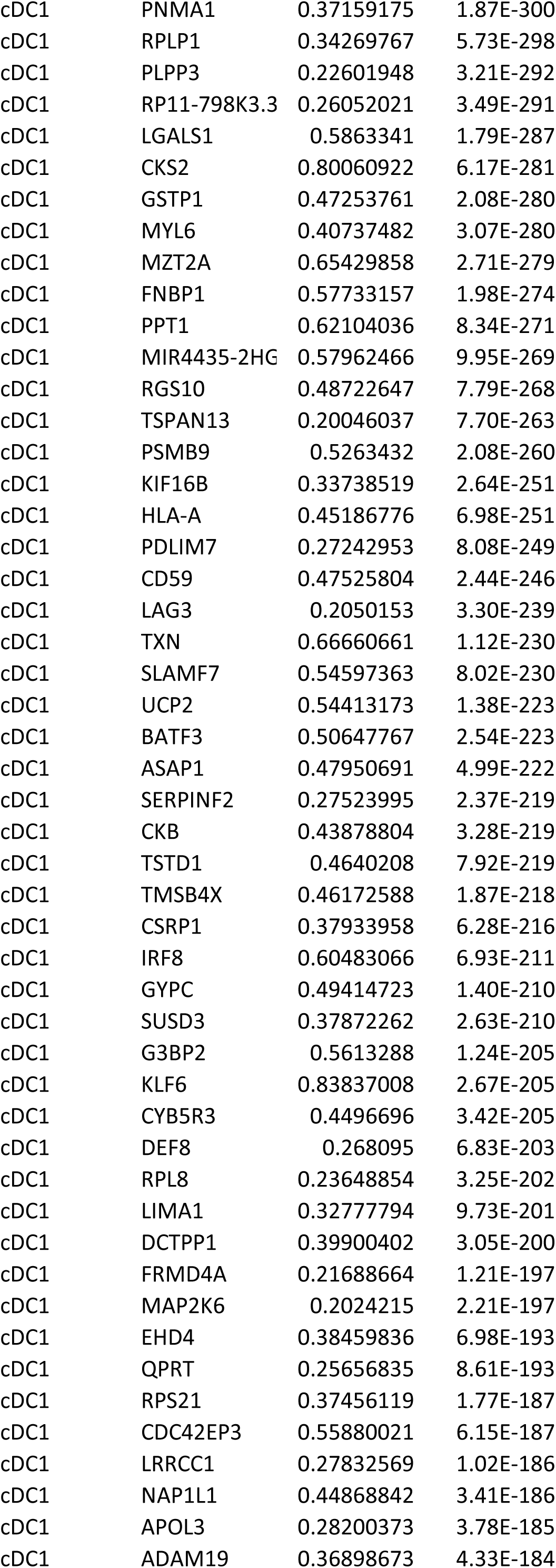
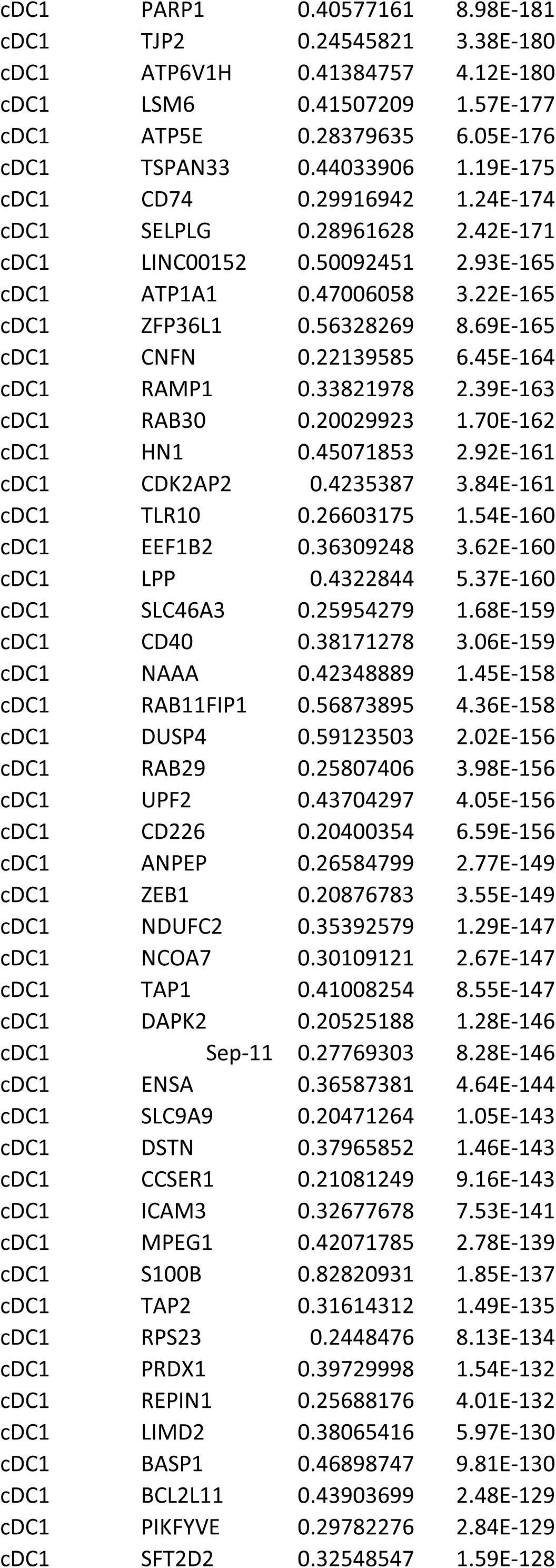
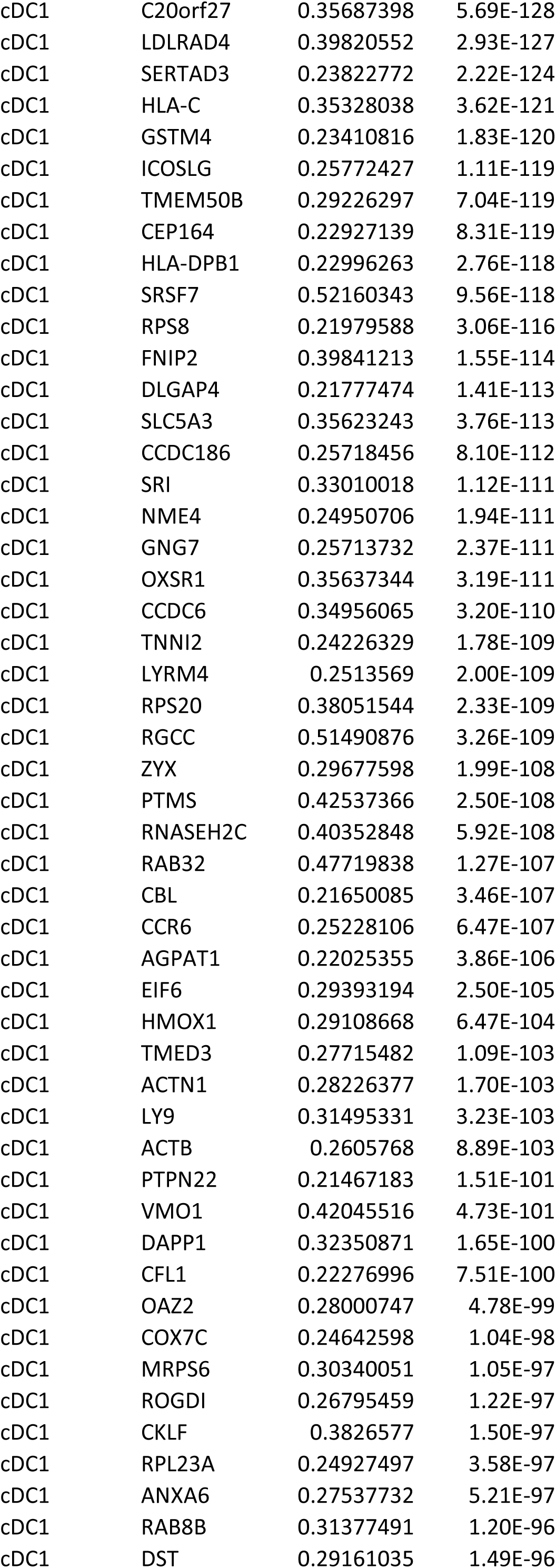
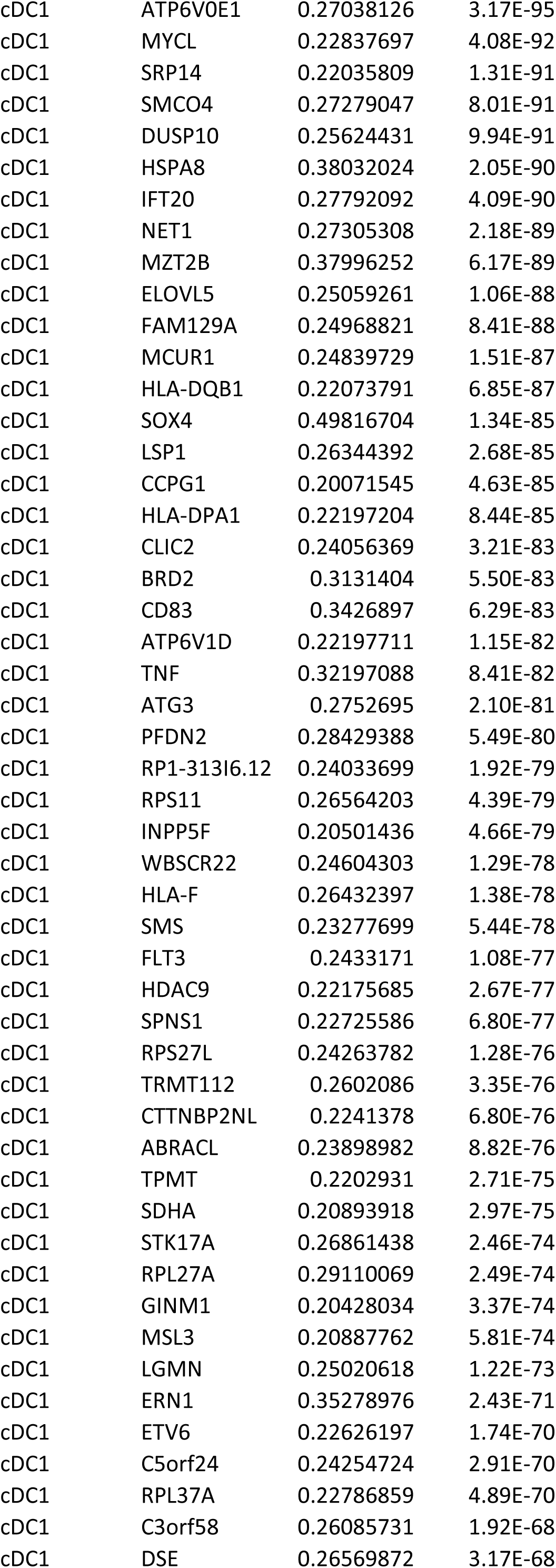
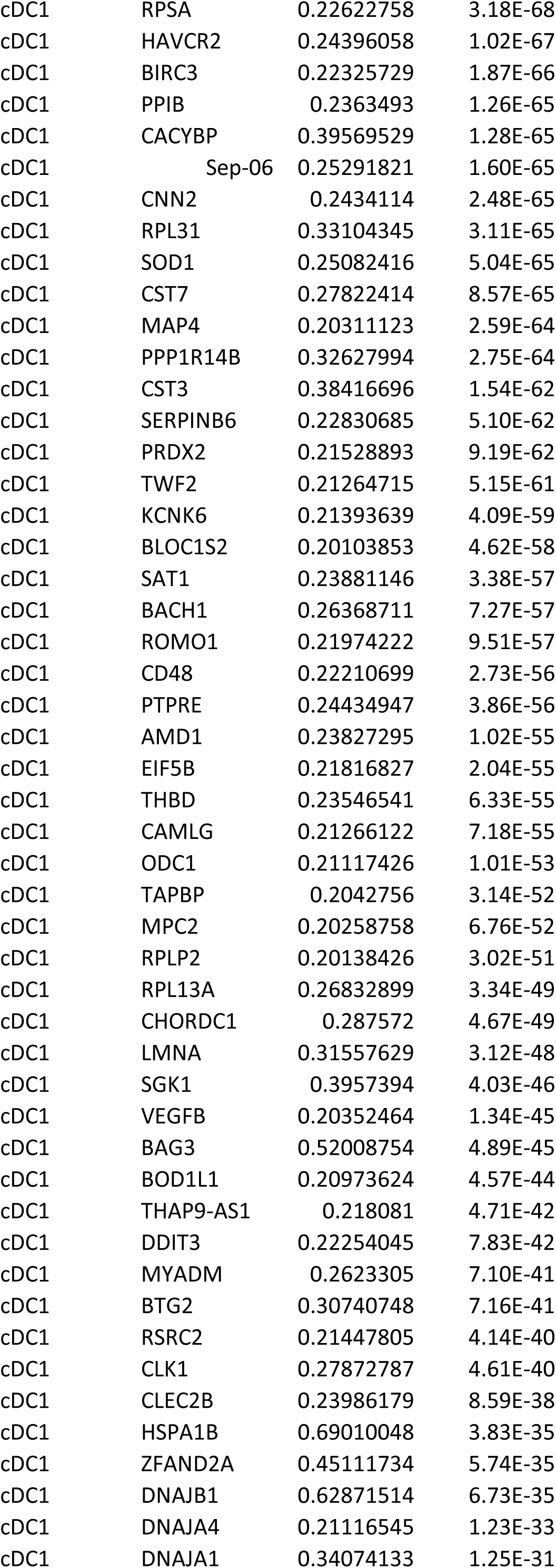
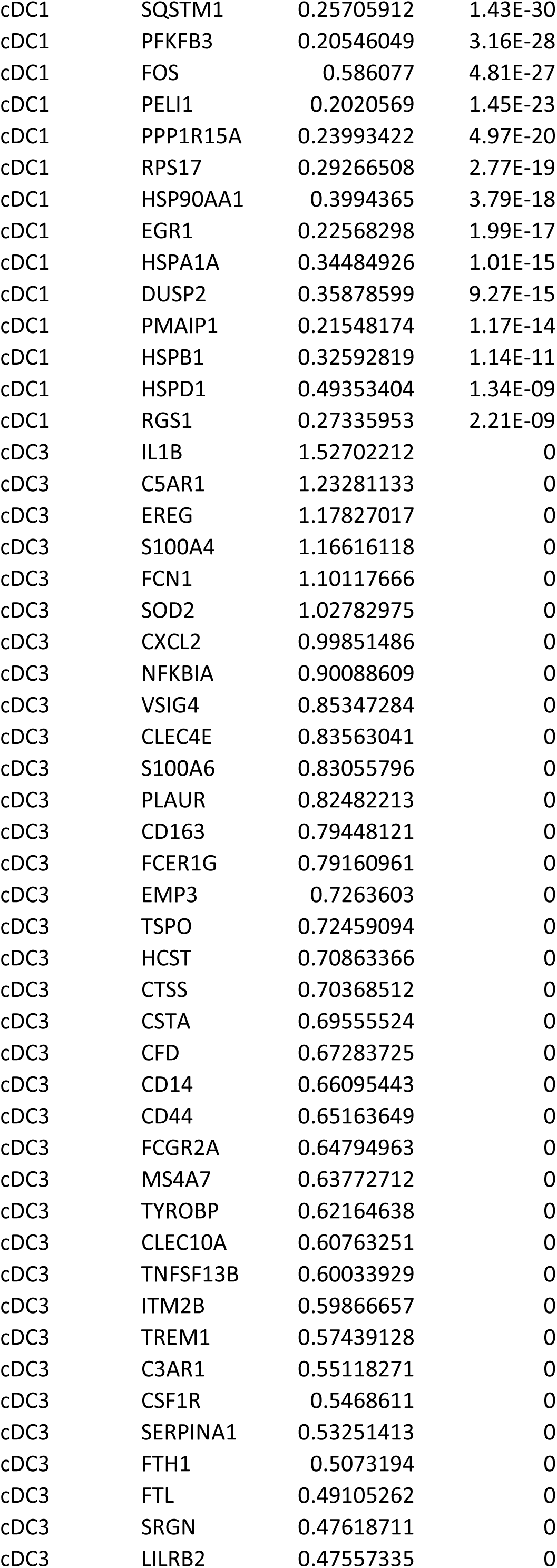
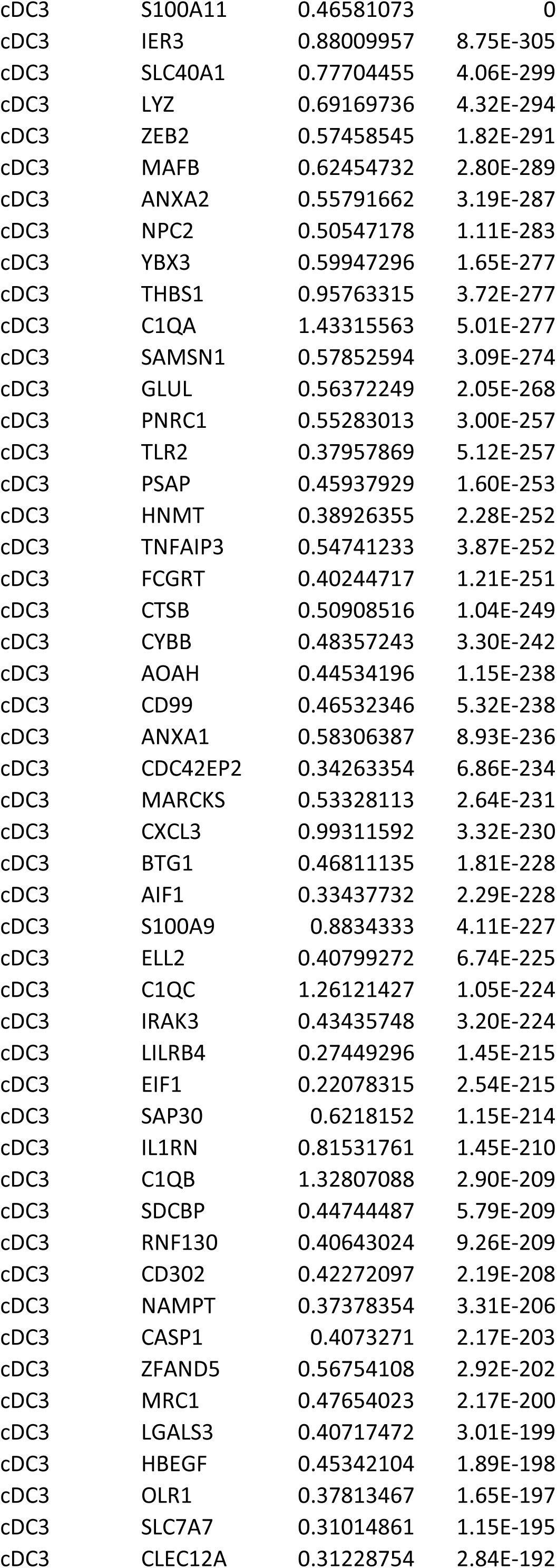
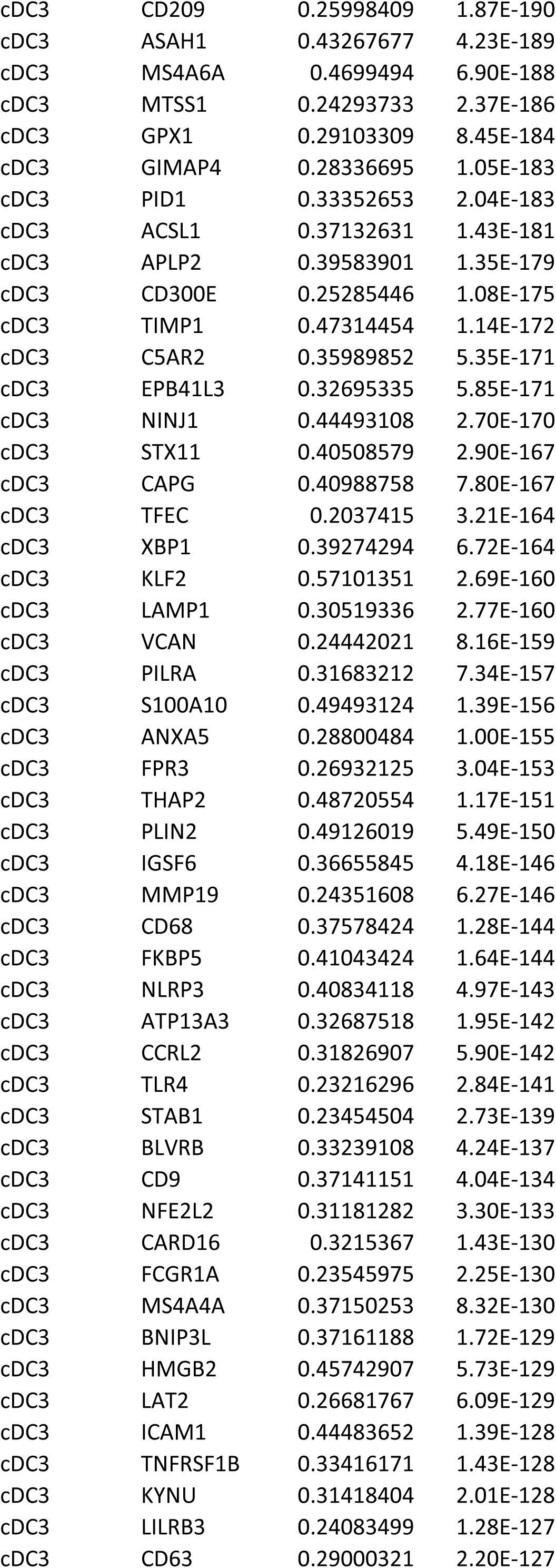
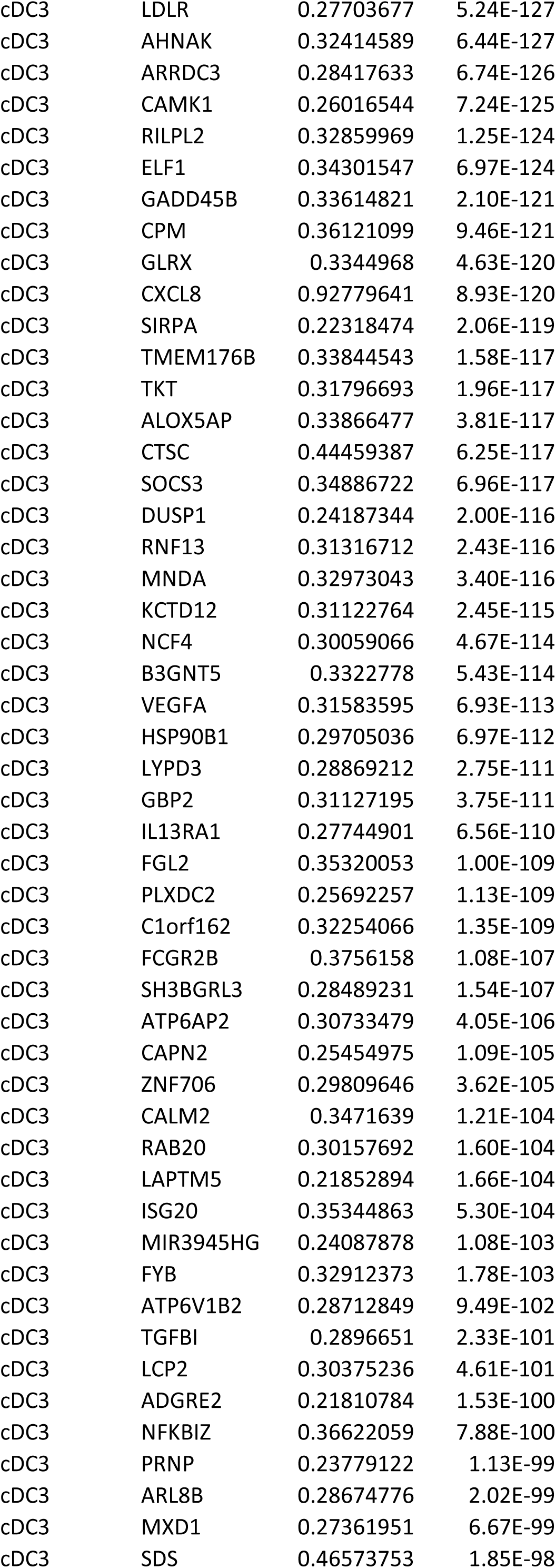
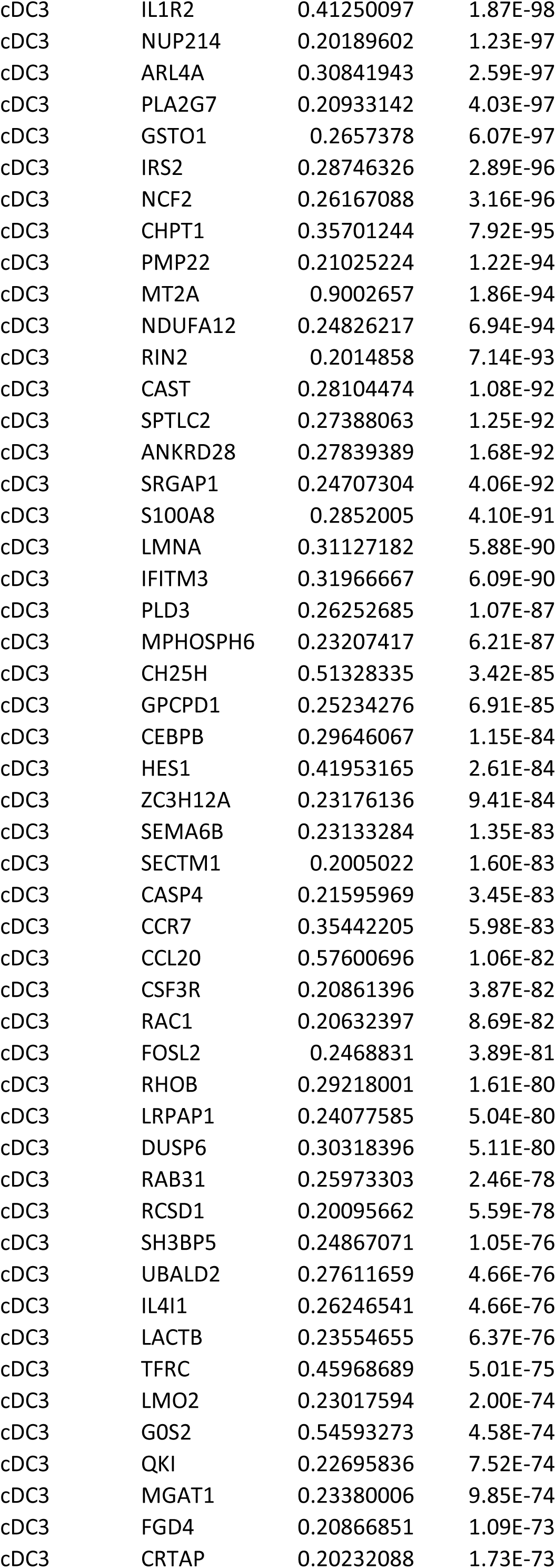
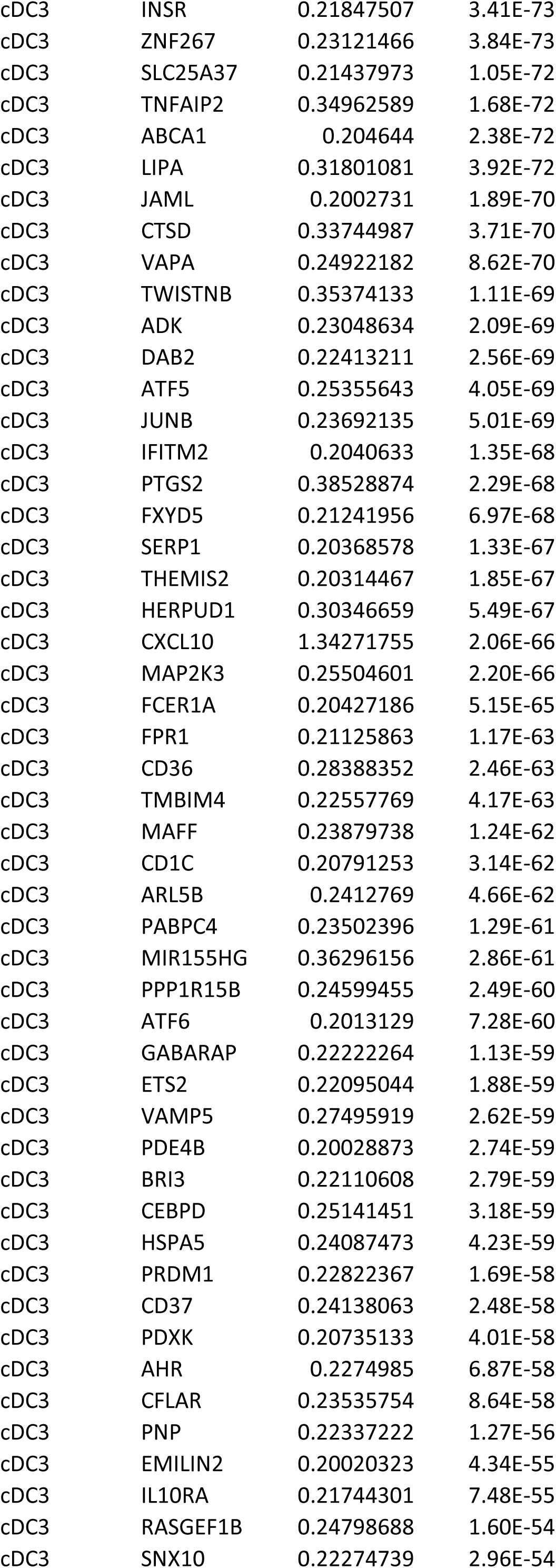
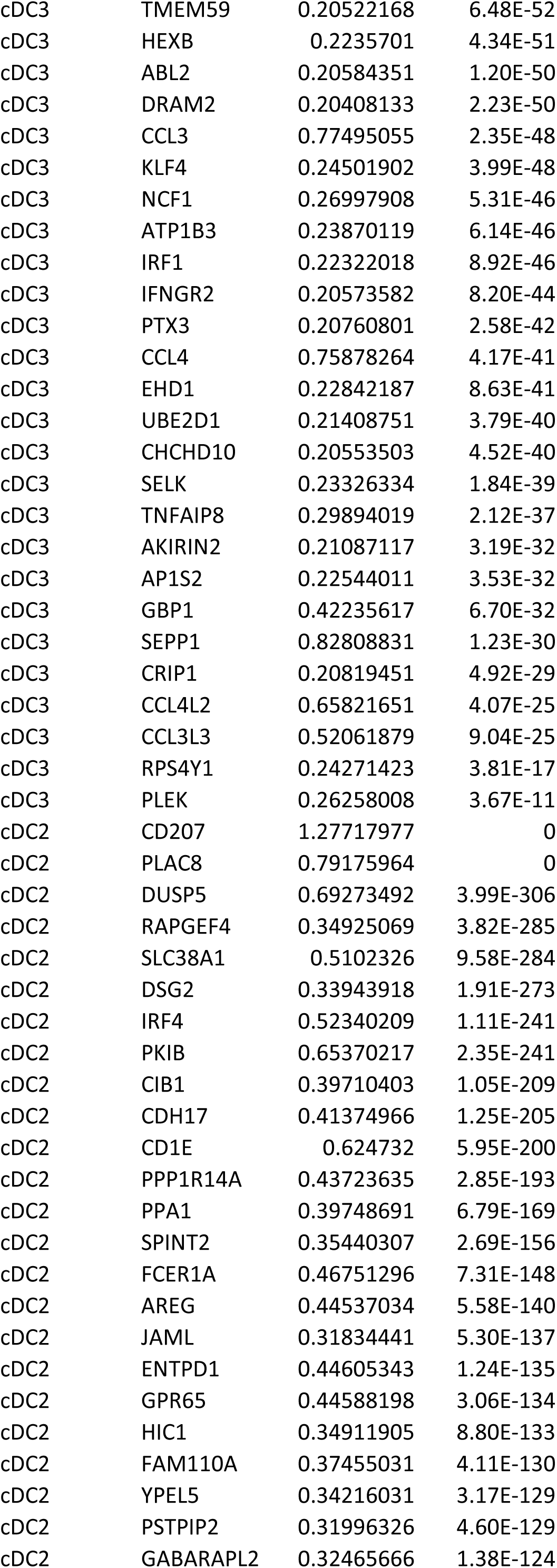
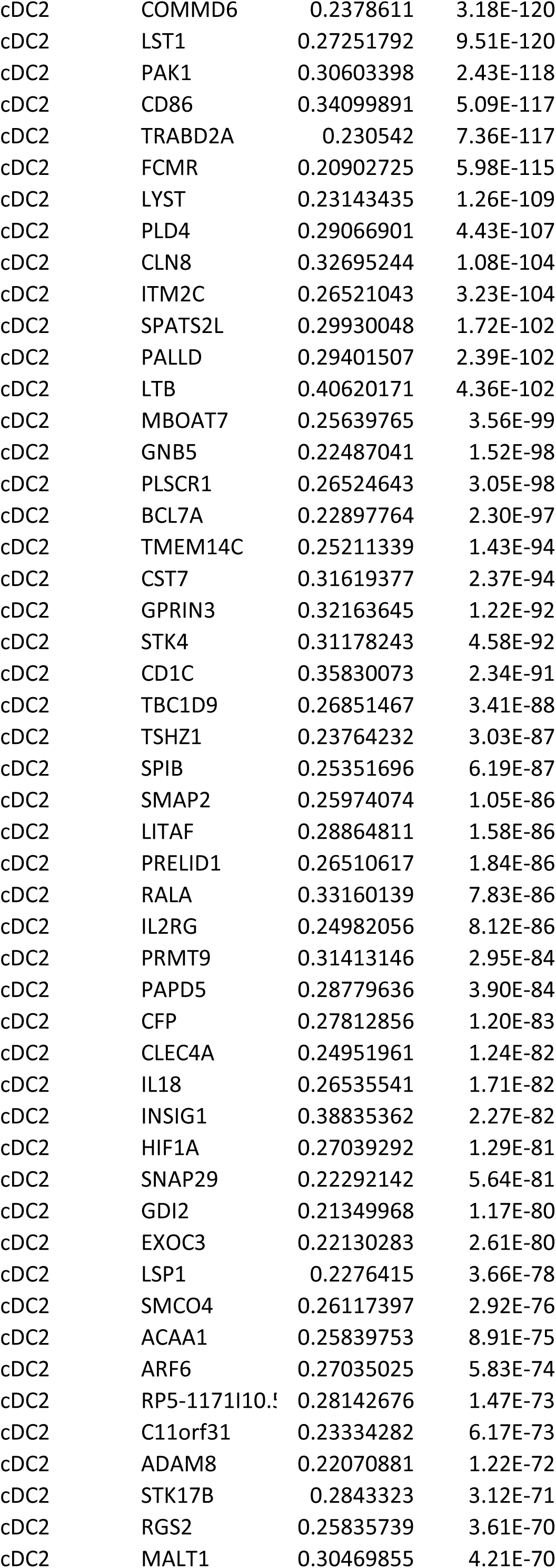
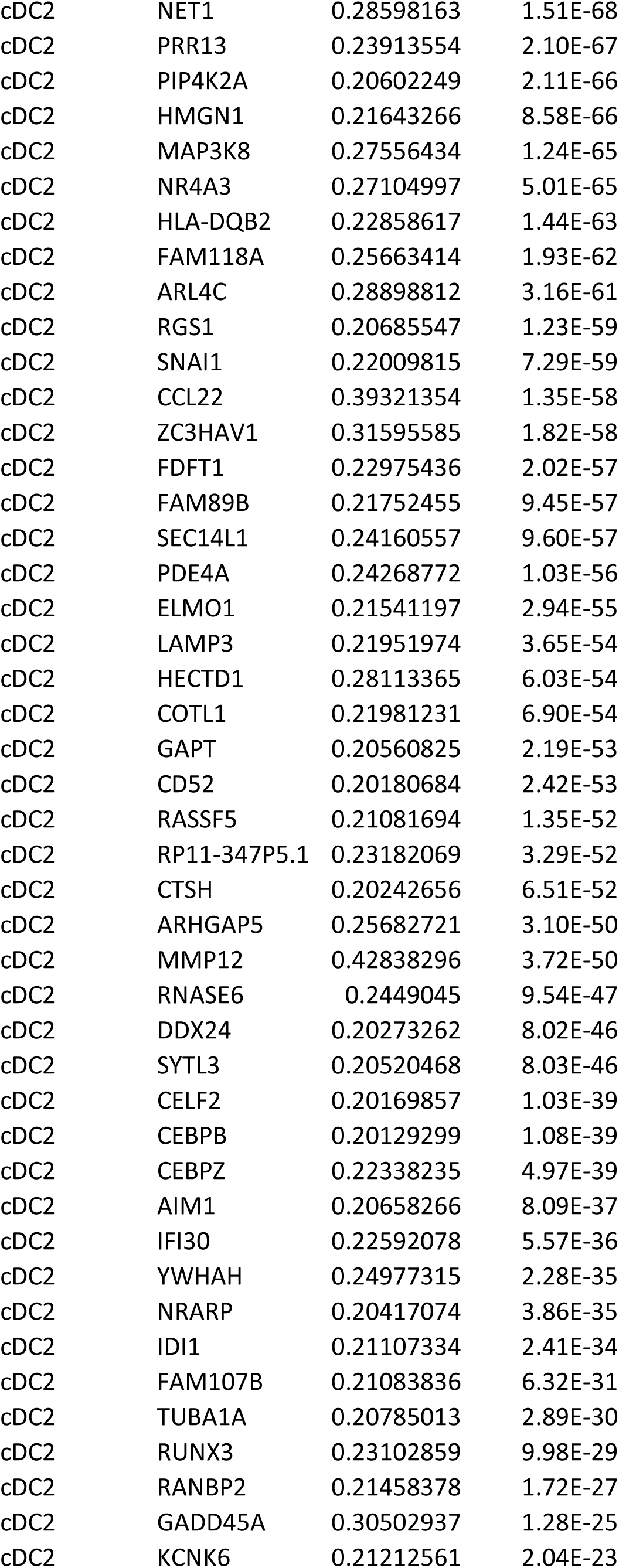
Complete list of DEG comparing clusters defined as cDC1, cDC2, and cDC3 for combined ileal and colonic LP.

**Table S3.**
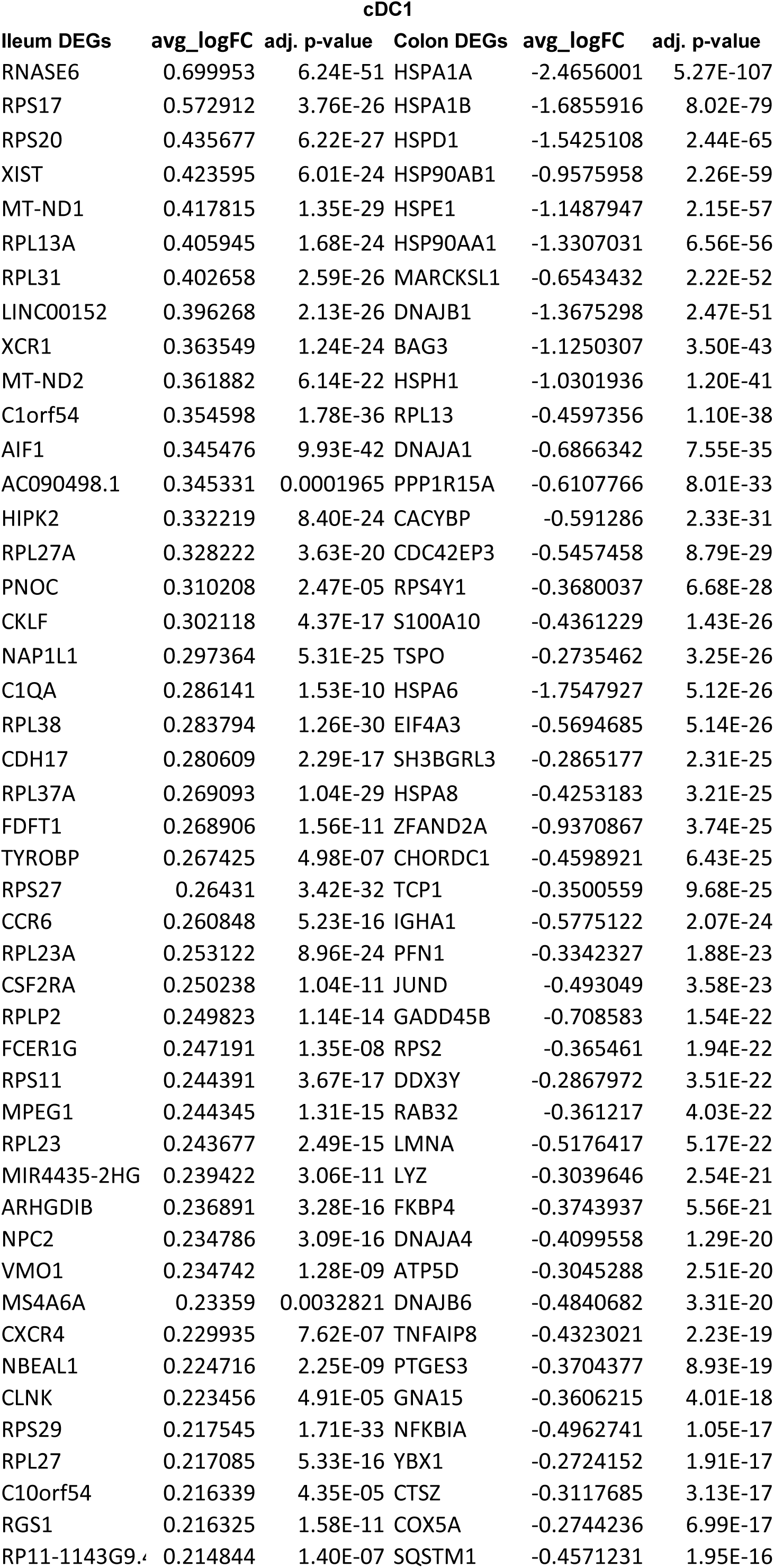
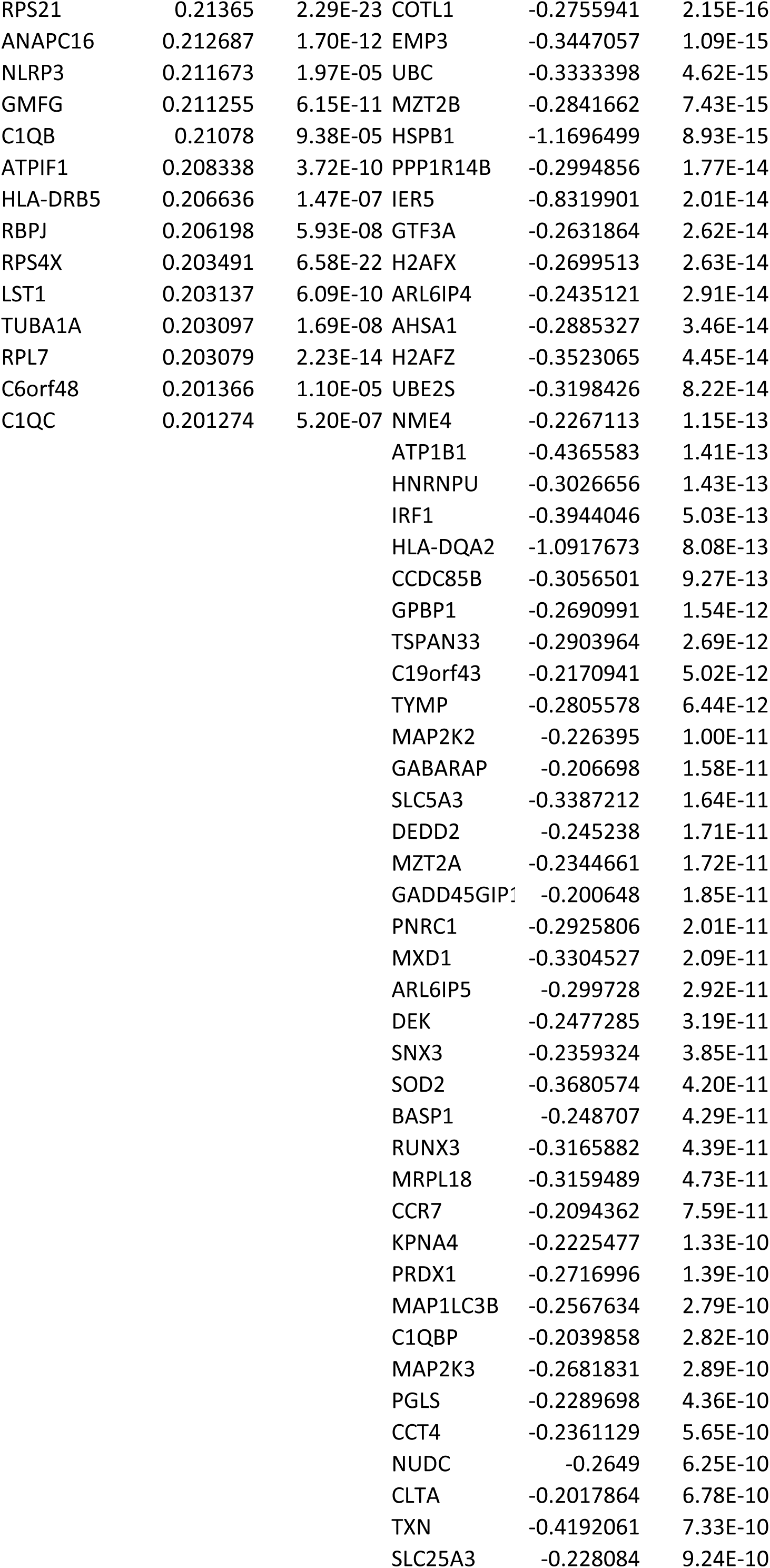
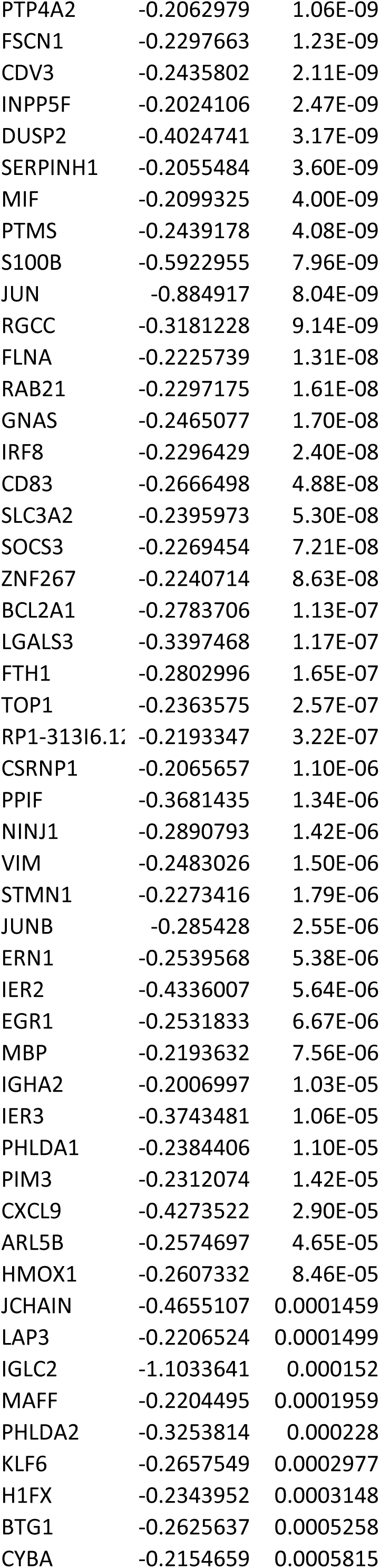
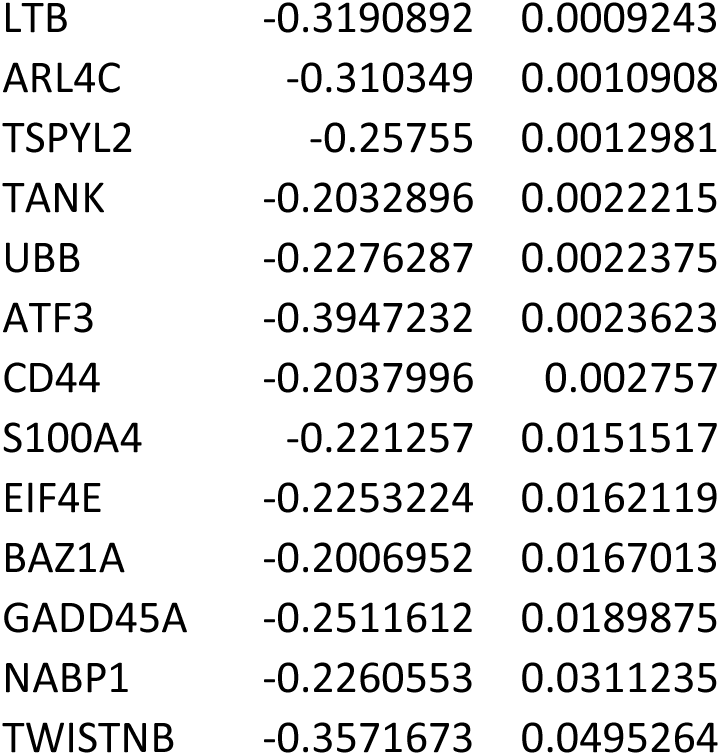
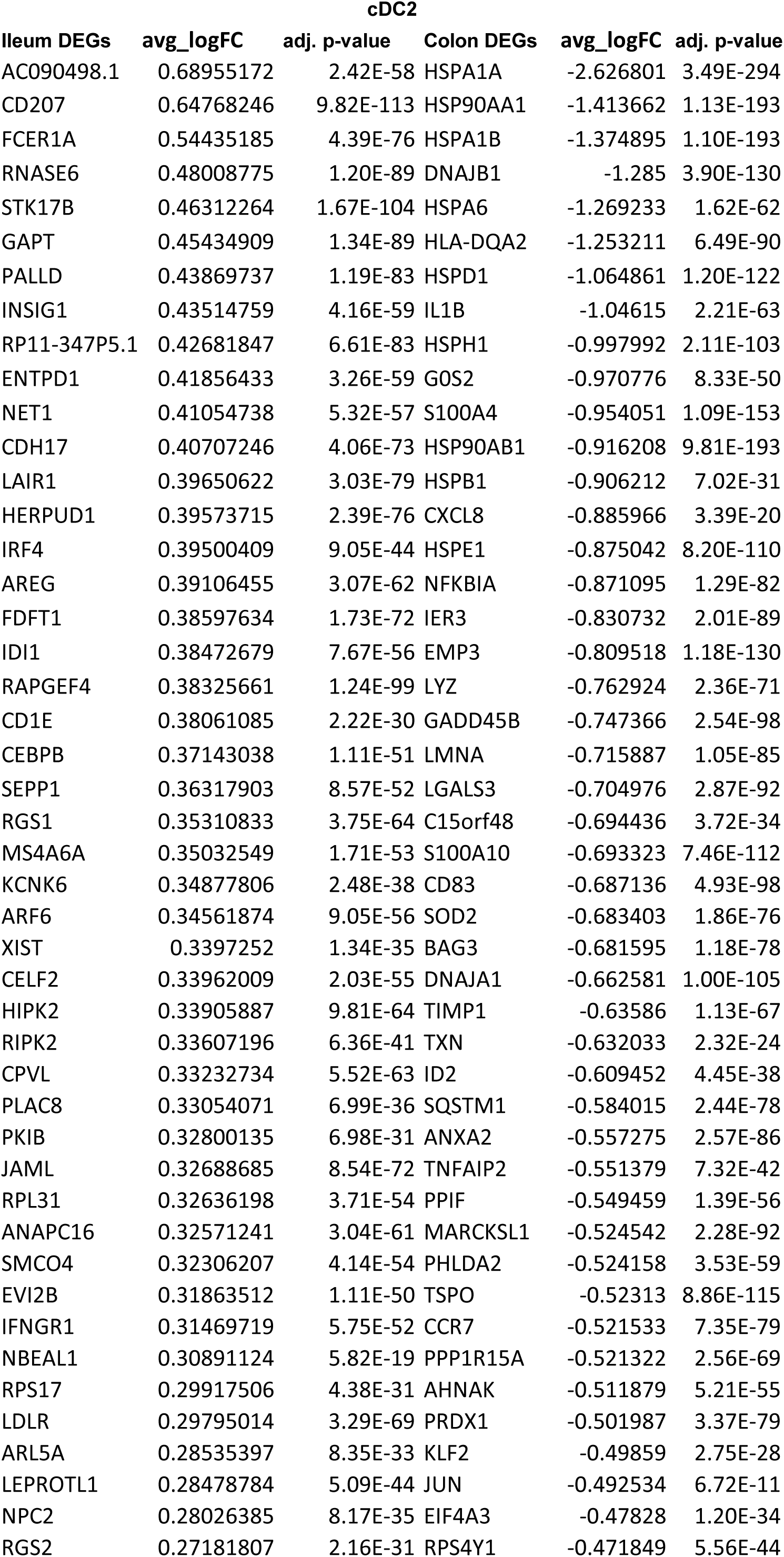
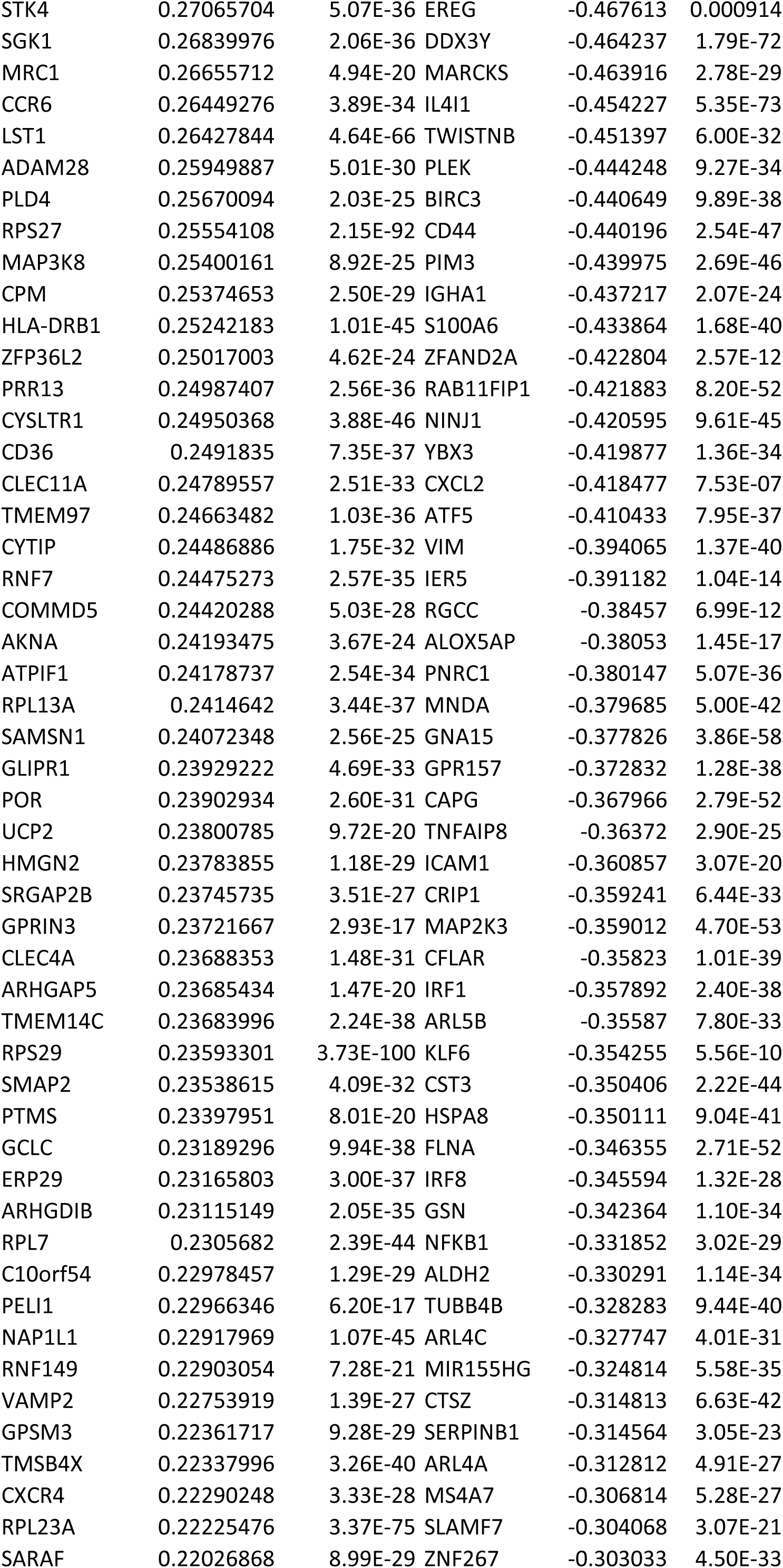
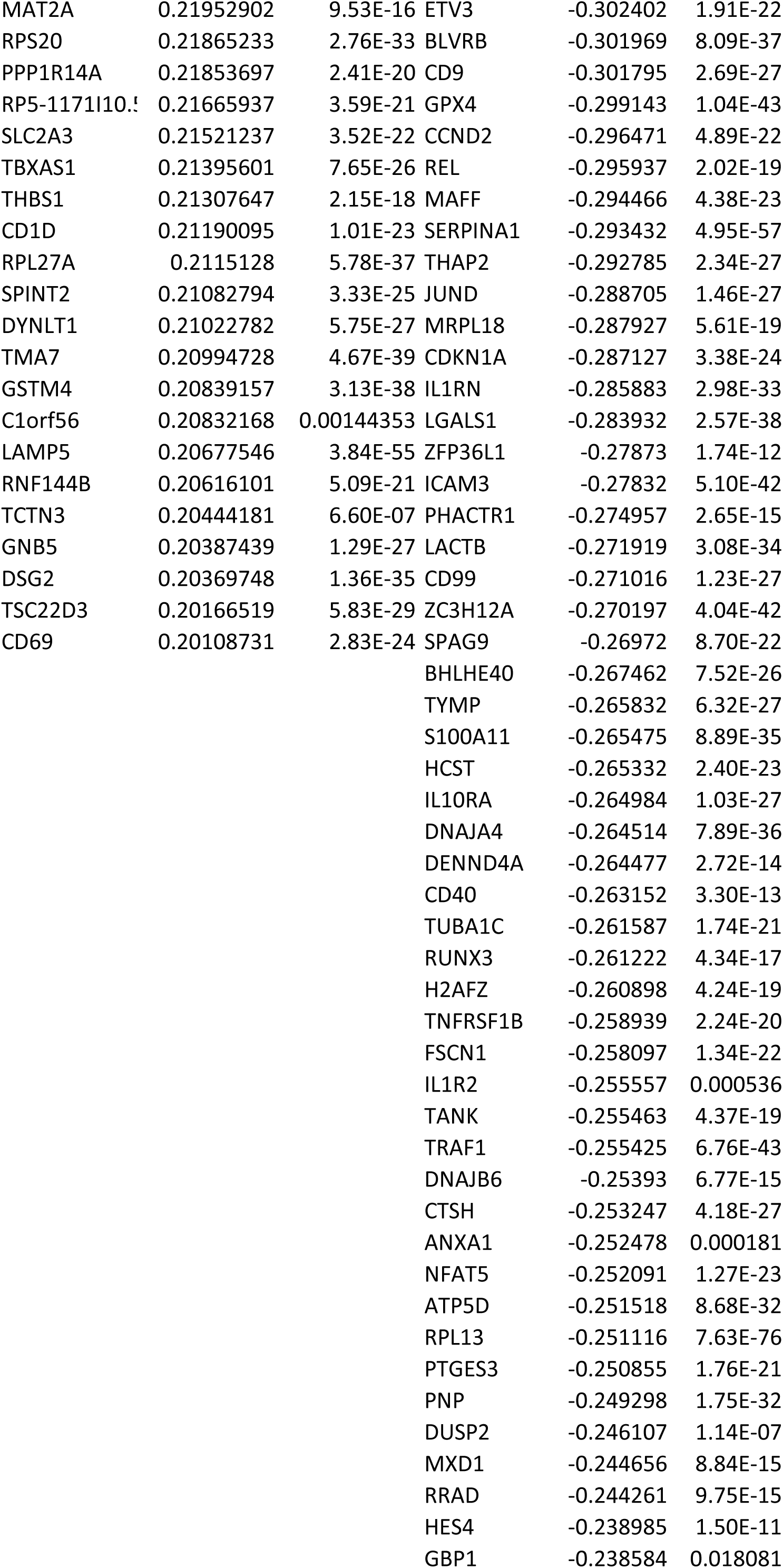
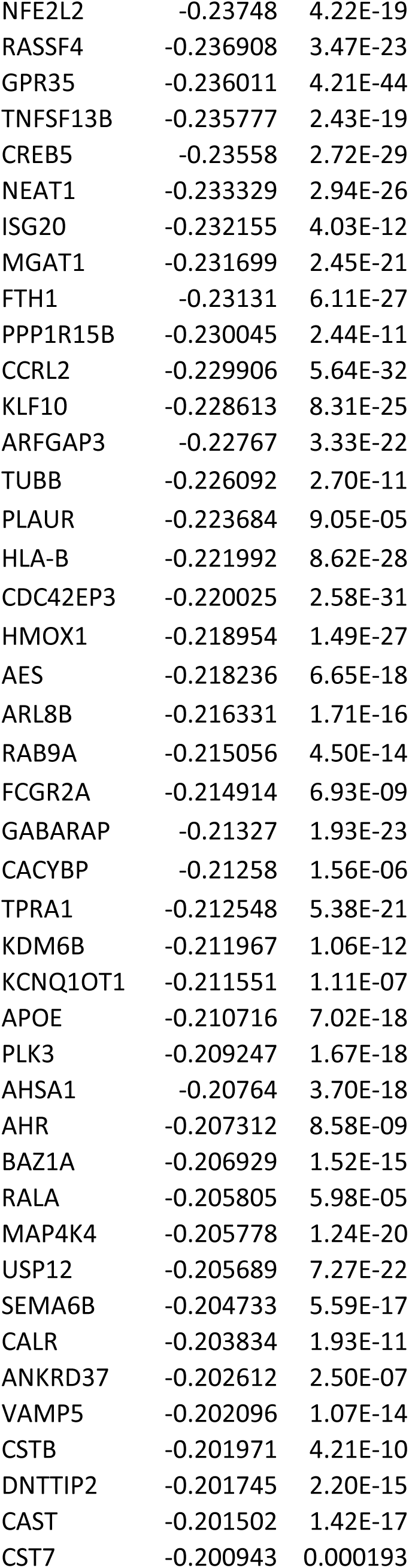
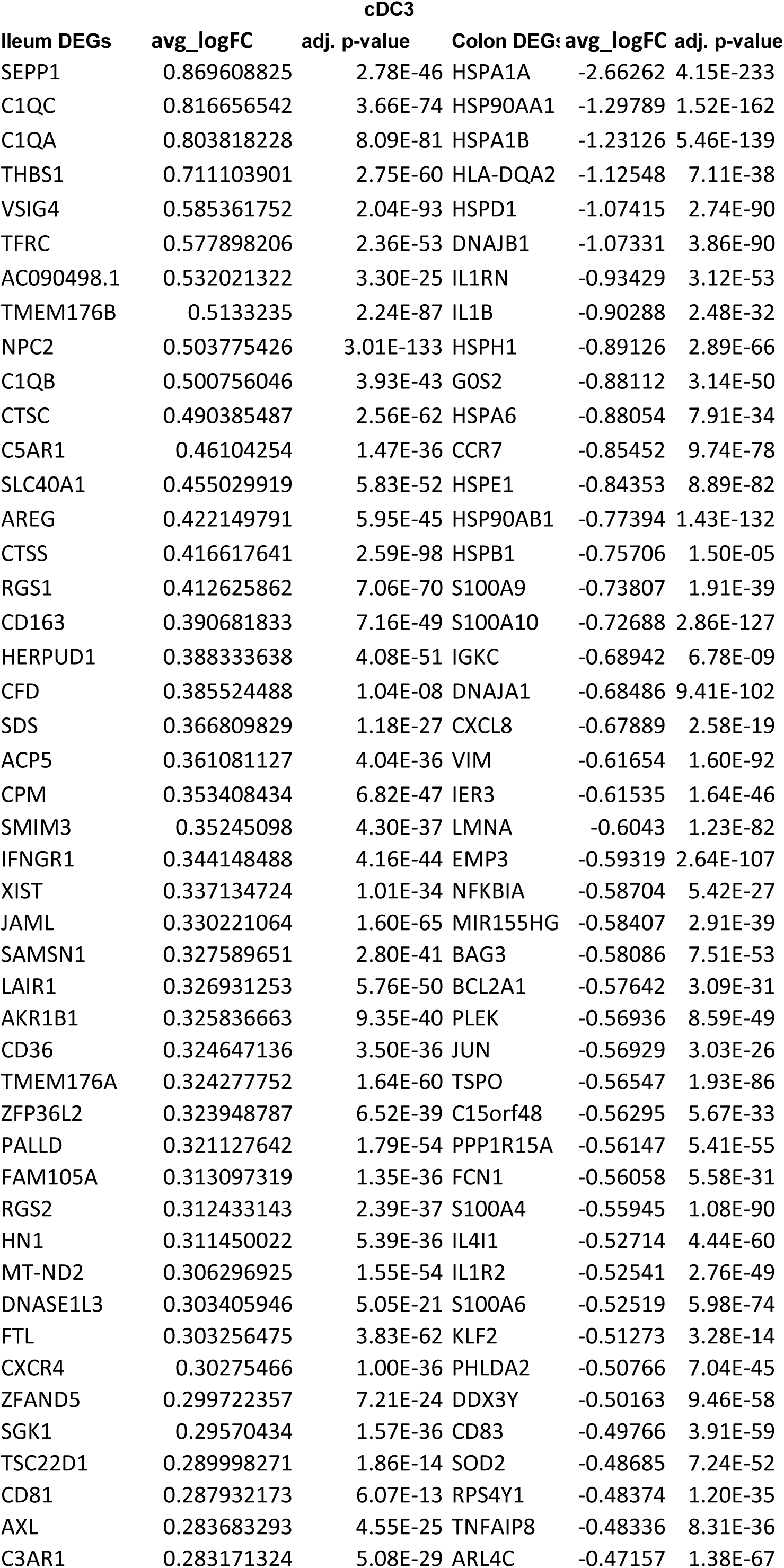
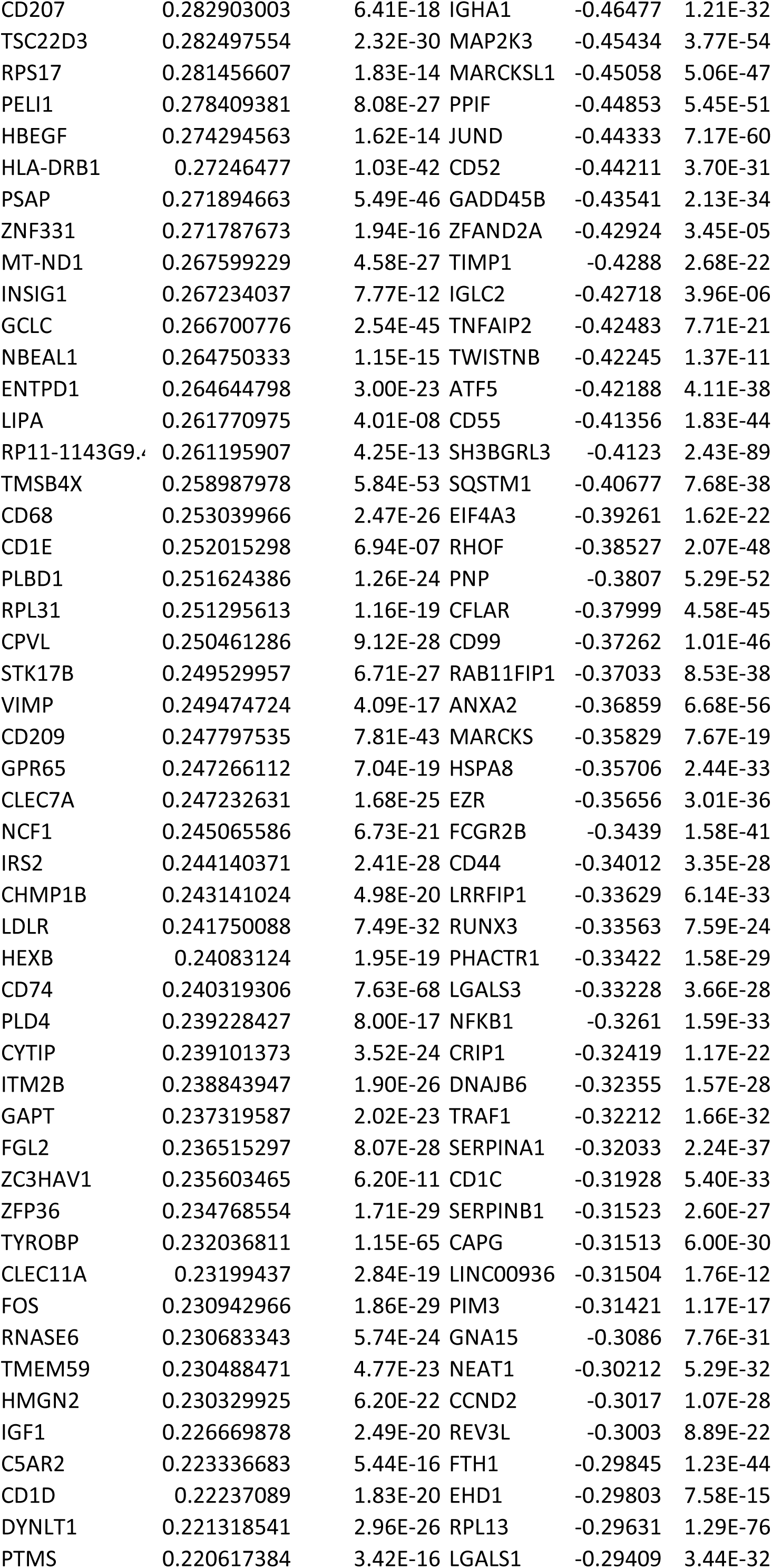
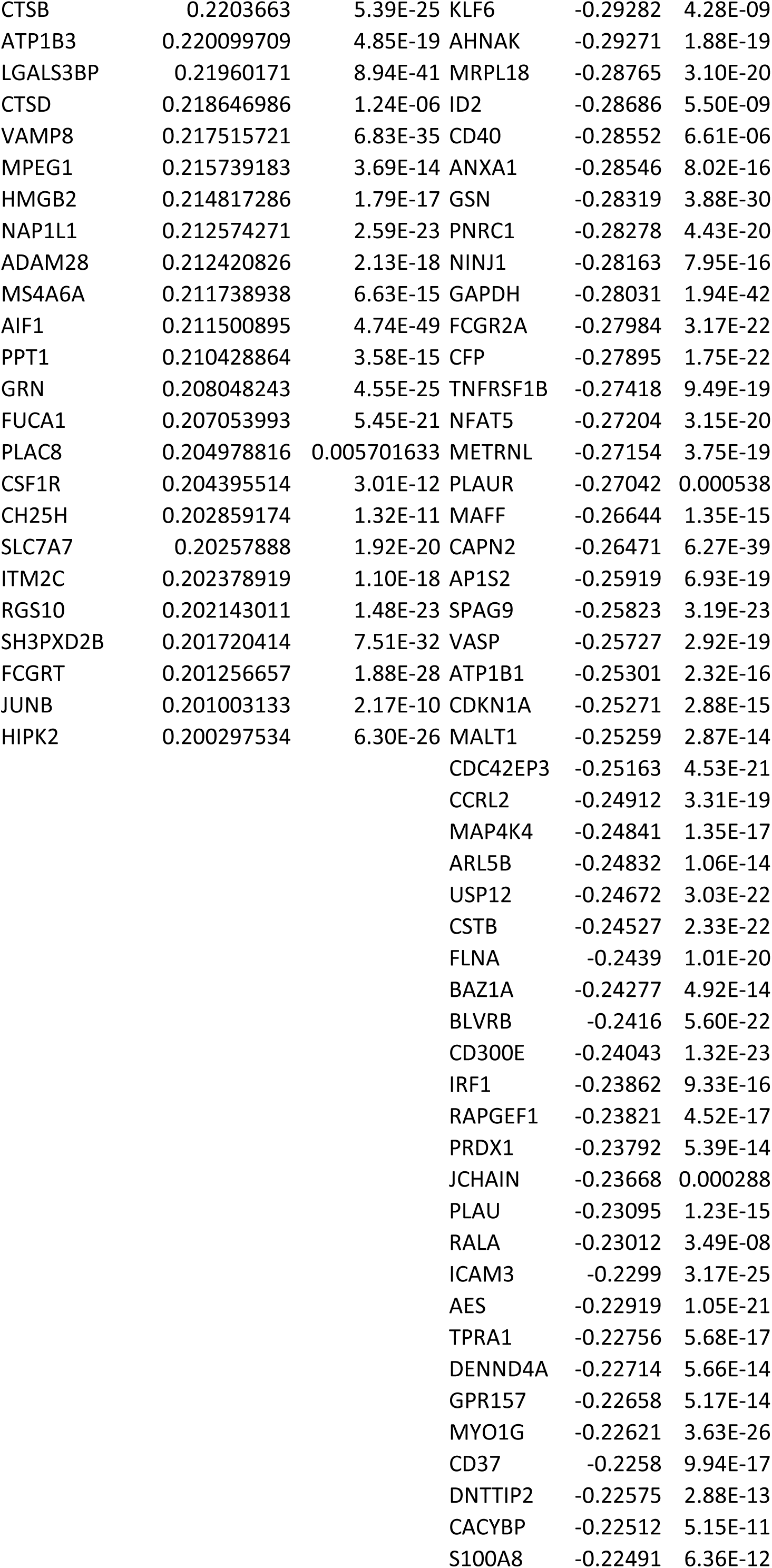
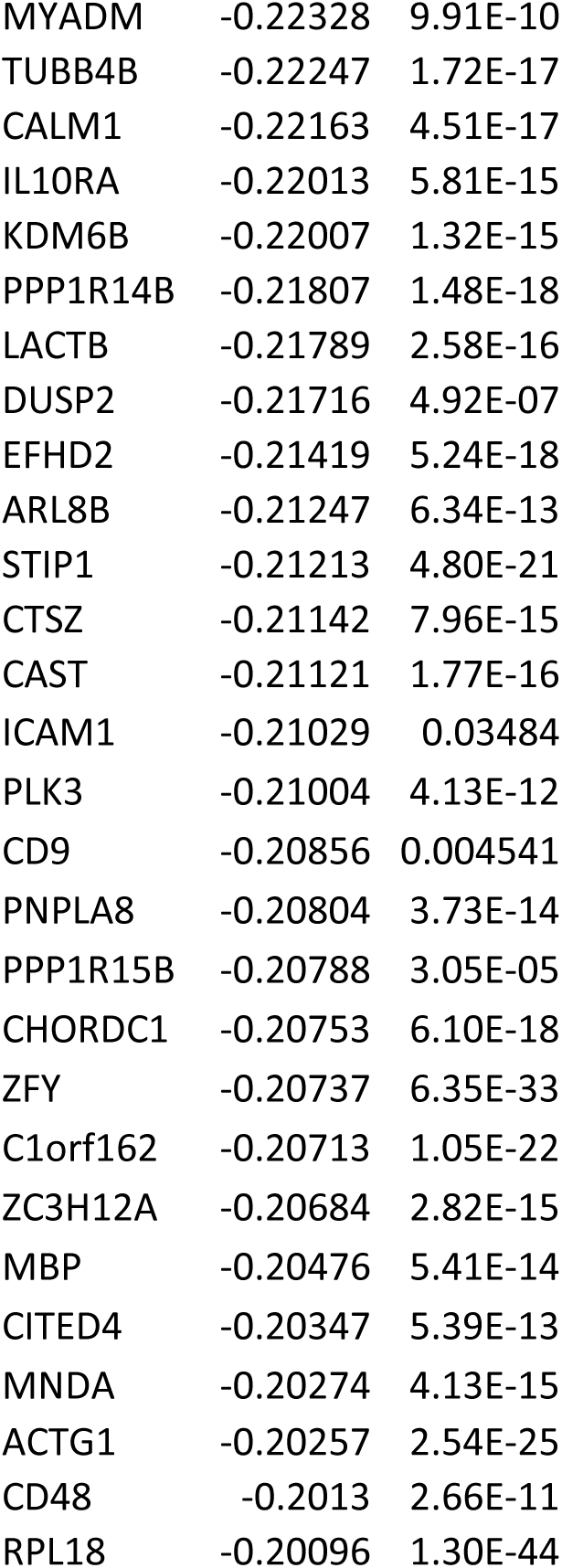
Complete list of DEG comparing ileal and colonic LP subsets for cDC1, cDC2 and cDC3.

**Table S4.** Complete list of DEG comparing mature macrophage clusters M7-M10, with data combined from ileal and colonic LP for each.

| cluster | gene | avg_logFC | p_val |
| --- | --- | --- | --- |
| M7 | FOSB | 1.009063 | 4.23E-276 |
| M7 | TNFAIP3 | 0.971054 | 5.07E-259 |
| M7 | MRC1 | 0.85316 | 1.41E-232 |
| M7 | KLF4 | 0.845135 | 3.19E-213 |
| M7 | NR4A2 | 0.852964 | 1.50E-207 |
| M7 | DUSP1 | 0.747596 | 2.21E-193 |
| M7 | FOS | 0.93288 | 2.97E-172 |
| M7 | CD83 | 0.620904 | 3.84E-170 |
| M7 | FOLR2 | 0.888235 | 9.32E-169 |
| M7 | KLF6 | 0.715811 | 2.92E-156 |
| M7 | CD163 | 0.612903 | 2.92E-148 |
| M7 | PNRC1 | 0.508571 | 2.85E-144 |
| M7 | ZFP36 | 0.522818 | 6.07E-132 |
| M7 | NEAT1 | 0.681366 | 9.53E-132 |
| M7 | SLC40A1 | 0.505537 | 1.53E-131 |
| M7 | ZFAND5 | 0.596218 | 2.78E-131 |
| M7 | EGR1 | 1.06785 | 1.02E-130 |
| M7 | F13A1 | 1.099542 | 7.28E-129 |
| M7 | TSC22D3 | 0.535036 | 5.21E-125 |
| M7 | CCL3 | 0.902018 | 2.06E-123 |
| M7 | JUNB | 0.517427 | 2.25E-122 |
| M7 | KLF2 | 0.80861 | 2.63E-121 |
| M7 | MAF | 0.540628 | 3.15E-118 |
| M7 | NFKBIZ | 0.581204 | 7.87E-112 |
| M7 | CCNL1 | 0.526068 | 1.80E-111 |
| M7 | HES1 | 0.698375 | 4.54E-111 |
| M7 | JUND | 0.510955 | 1.06E-110 |
| M7 | JUN | 0.816444 | 1.31E-110 |
| M7 | ICAM1 | 0.588734 | 2.77E-110 |
| M7 | MAFF | 0.590731 | 6.69E-108 |
| M7 | KDM6B | 0.566587 | 2.33E-103 |
| M7 | IER3 | 0.712718 | 2.96E-103 |
| M7 | STAB1 | 0.622278 | 1.38E-99 |
| M7 | IER2 | 0.674185 | 5.07E-99 |
| M7 | LYVE1 | 0.692658 | 1.53E-97 |
| M7 | ATF3 | 0.667888 | 2.40E-97 |
| M7 | CCL4 | 0.856588 | 4.37E-96 |
| M7 | HSPA5 | 0.518535 | 1.78E-94 |
| M7 | CXCL2 | 0.780531 | 3.09E-94 |
| M7 | LTC4S | 0.589625 | 7.54E-94 |
| M7 | ELL2 | 0.553941 | 1.25E-84 |
| M7 | CXCL8 | 0.590623 | 3.14E-81 |
| M7 | PDK4 | 0.657982 | 2.34E-79 |
| M7 | BTG2 | 0.579485 | 1.26E-78 |
| M7 | CCL4L2 | 0.755012 | 4.10E-76 |
| M7 | CXCR4 | 0.538895 | 2.11E-75 |
| M7 | IL1B | 0.599742 | 2.99E-72 |
| M7 | PTGS2 | 0.634458 | 6.29E-72 |
| M7 | MYADM | 0.508809 | 4.69E-69 |
| M7 | TRA2B | 0.546561 | 1.65E-65 |
| M7 | NR4A1 | 0.506879 | 2.39E-64 |
| M7 | NLRP3 | 0.538157 | 2.68E-59 |
| M7 | DUSP6 | 0.518986 | 4.77E-57 |
| M7 | CCL3L3 | 0.813307 | 4.25E-52 |
| M7 | HSPA1B | 0.533624 | 1.97E-50 |
| M7 | CXCL3 | 0.578672 | 8.07E-49 |
| M7 | PRDM1 | 0.501032 | 1.10E-46 |
| M7 | RNASE1 | 0.611347 | 6.06E-40 |
| M7 | THBS1 | 0.553799 | 7.57E-31 |
| M7 | IGLC3 | 0.523912 | 0.000629 |
| M8 | ACP5 | 0.790933 | 1.36E-174 |
| M8 | LGALS3 | 1.005682 | 1.08E-170 |
| M8 | CD9 | 0.692784 | 4.52E-132 |
| M8 | SDS | 0.704316 | 2.94E-130 |
| M8 | LYZ | 0.593559 | 4.88E-125 |
| M8 | FABP5 | 0.638491 | 9.66E-85 |
| M8 | GPNMB | 0.580736 | 1.62E-82 |
| M8 | MMP12 | 0.51479 | 1.69E-78 |
| M8 | TIMP1 | 0.500267 | 3.27E-74 |
| M8 | APOC1 | 0.508046 | 1.47E-33 |
| M8 | MT2A | 0.566905 | 2.70E-25 |
| M8 | MT1G | 0.523668 | 1.14E-23 |
| M8 | MT1X | 0.608234 | 7.40E-20 |
| M9 | IL22RA2 | 1.170857 | 9.76E-226 |
| M9 | NRG1 | 0.70072 | 3.41E-167 |
| M9 | PLA2G2D | 0.841636 | 3.28E-120 |
| M9 | IL2RG | 0.839857 | 5.79E-91 |
| M9 | MMP12 | 2.122317 | 6.28E-88 |
| M9 | TNFRSF9 | 0.567362 | 1.63E-77 |
| M9 | IL7R | 0.801193 | 1.60E-67 |
| M9 | MMP9 | 1.932434 | 1.50E-62 |
| M9 | ENPP2 | 1.191284 | 1.12E-60 |
| M9 | LGALS2 | 1.323239 | 1.59E-58 |
| M9 | IL4I1 | 0.830911 | 4.47E-55 |
| M9 | PTGDS | 2.020959 | 2.84E-54 |
| M9 | HLA-DQB2 | 0.734882 | 7.48E-50 |
| M9 | ATOX1 | 1.152716 | 1.54E-49 |
| M9 | CST7 | 0.74162 | 4.94E-47 |
| M9 | TMEM176 | 0.851909 | 2.78E-46 |
| M9 | CTSH | 0.895847 | 5.36E-44 |
| M9 | TMEM176 | 0.780614 | 7.90E-44 |
| M9 | LYZ | 1.106362 | 2.33E-43 |
| M9 | IL1RN | 0.893798 | 2.83E-43 |
| M9 | GSTP1 | 0.694707 | 2.41E-42 |
| M9 | S100B | 1.054142 | 8.93E-42 |
| M9 | SERPINA1 | 0.655293 | 5.70E-41 |
| M9 | ADAMDEC | 0.783831 | 7.92E-39 |
| M9 | ARHGDIB | 0.662509 | 1.07E-38 |
| M9 | DNASE1L3 | 0.915718 | 4.43E-37 |
| M9 | BASP1 | 0.77034 | 1.21E-36 |
| M9 | ENO1 | 0.634591 | 1.61E-35 |
| M9 | CAPG | 1.020336 | 2.83E-35 |
| M9 | GSN | 0.745927 | 5.91E-34 |
| M9 | PLA2G7 | 0.651554 | 8.98E-33 |
| M9 | PRELID1 | 0.69906 | 1.45E-30 |
| M9 | PPA1 | 0.790676 | 1.92E-30 |
| M9 | ALOX5AP | 1.278542 | 4.85E-29 |
| M9 | ITGB2 | 0.53707 | 1.37E-25 |
| M9 | CSTA | 0.585328 | 2.20E-25 |
| M9 | SERPINF1 | 0.659014 | 2.55E-25 |
| M9 | SYNGR2 | 0.573878 | 5.68E-25 |
| M9 | PRDX1 | 0.509078 | 7.01E-23 |
| M9 | RALA | 0.506197 | 2.22E-22 |
| M9 | EFHD2 | 0.544071 | 5.66E-22 |
| M9 | C15orf48 | 0.66877 | 1.98E-20 |
| M9 | TYMP | 0.552502 | 2.85E-20 |
| M9 | COTL1 | 0.512371 | 9.51E-20 |
| M9 | MYDGF | 0.528733 | 3.76E-19 |
| M9 | PFN1 | 0.555179 | 2.85E-18 |
| M9 | CXCL9 | 1.167317 | 7.34E-14 |
| M9 | FUCA1 | 0.566332 | 2.17E-12 |
| M9 | IL18 | 0.542143 | 7.13E-12 |
| M9 | GOS2 | 0.528025 | 3.84E-08 |
| M10 | FTL | 0.717363 | 2.45E-92 |
| M10 | OAZ1 | 0.509002 | 1.12E-66 |
| M10 | GAPDH | 0.679266 | 2.96E-63 |
| M10 | RPL7A | 0.543585 | 1.31E-62 |
| M10 | YBX1 | 0.583305 | 6.37E-53 |
| M10 | S100A11 | 0.619082 | 3.37E-45 |
| M10 | GNAS | 0.570088 | 4.66E-41 |
| M10 | GSTP1 | 0.557185 | 5.09E-41 |
| M10 | BRI3 | 0.569526 | 2.18E-33 |
| M10 | APOE | 1.210938 | 2.01E-27 |
| M10 | FUCA1 | 0.980098 | 2.51E-26 |
| M10 | LGALS3 | 0.695002 | 2.45E-21 |
| M10 | PRDX1 | 0.598175 | 1.85E-19 |
| M10 | NUPR1 | 0.720171 | 3.18E-19 |
| M10 | CSTB | 0.734237 | 3.53E-19 |
| M10 | CTSD | 0.665014 | 1.90E-16 |
| M10 | APOC1 | 0.843922 | 2.43E-14 |
| M10 | ATP6V0D2 | 0.595397 | 8.64E-10 |
| M10 | PLA2G7 | 0.675315 | 7.28E-09 |
| M10 | VEGFB | 0.571188 | 3.68E-07 |
| M10 | MT1G | 1.007437 | 7.43E-07 |
| M10 | CRYAB | 0.776308 | 8.32E-07 |
| M10 | RAB42 | 0.538144 | 7.49E-06 |
| M10 | MT1X | 0.744601 | 2.01E-05 |
| M10 | MT1E | 0.557885 | 0.00012 |
| M10 | MT2A | 0.615267 | 0.000725 |

**Table S5.**
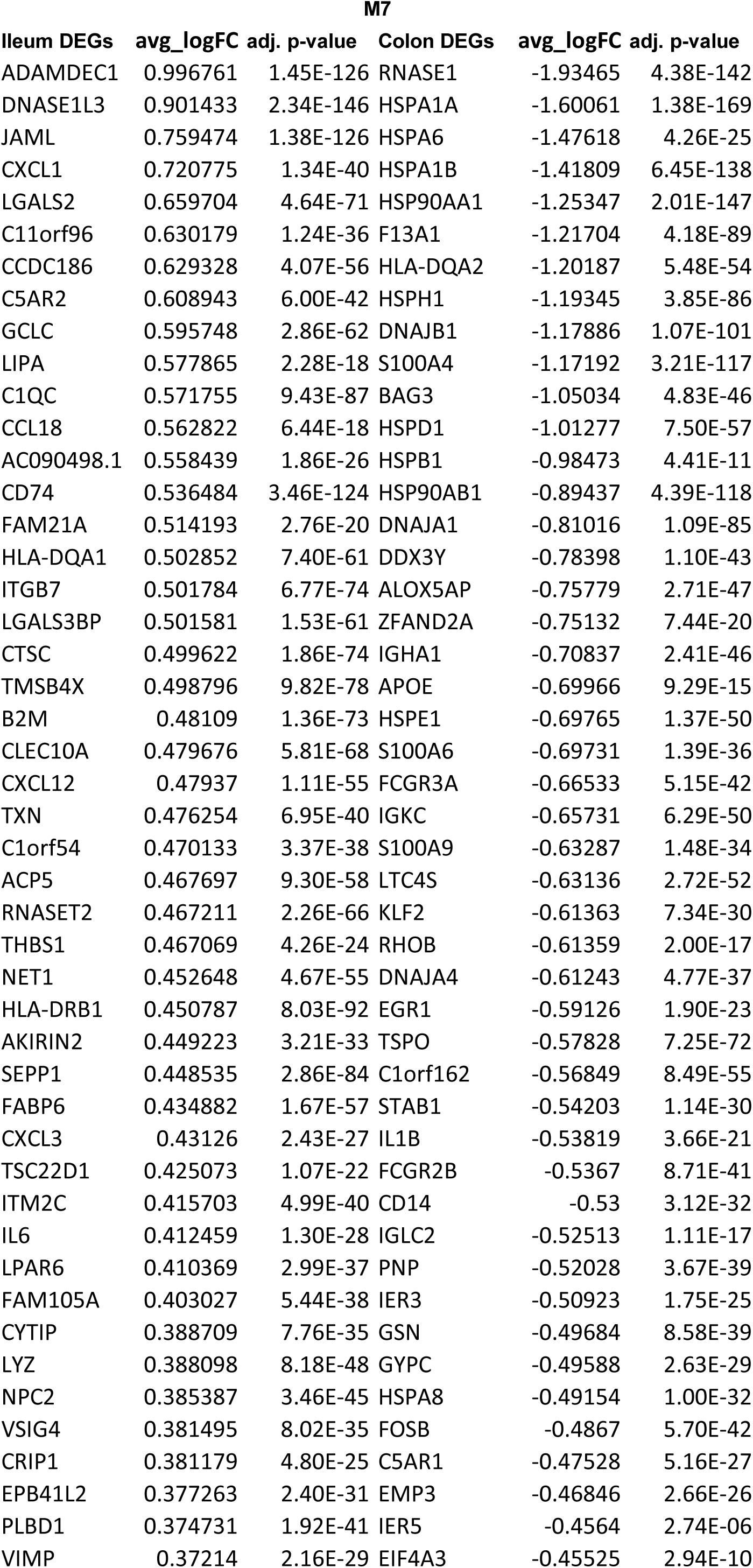
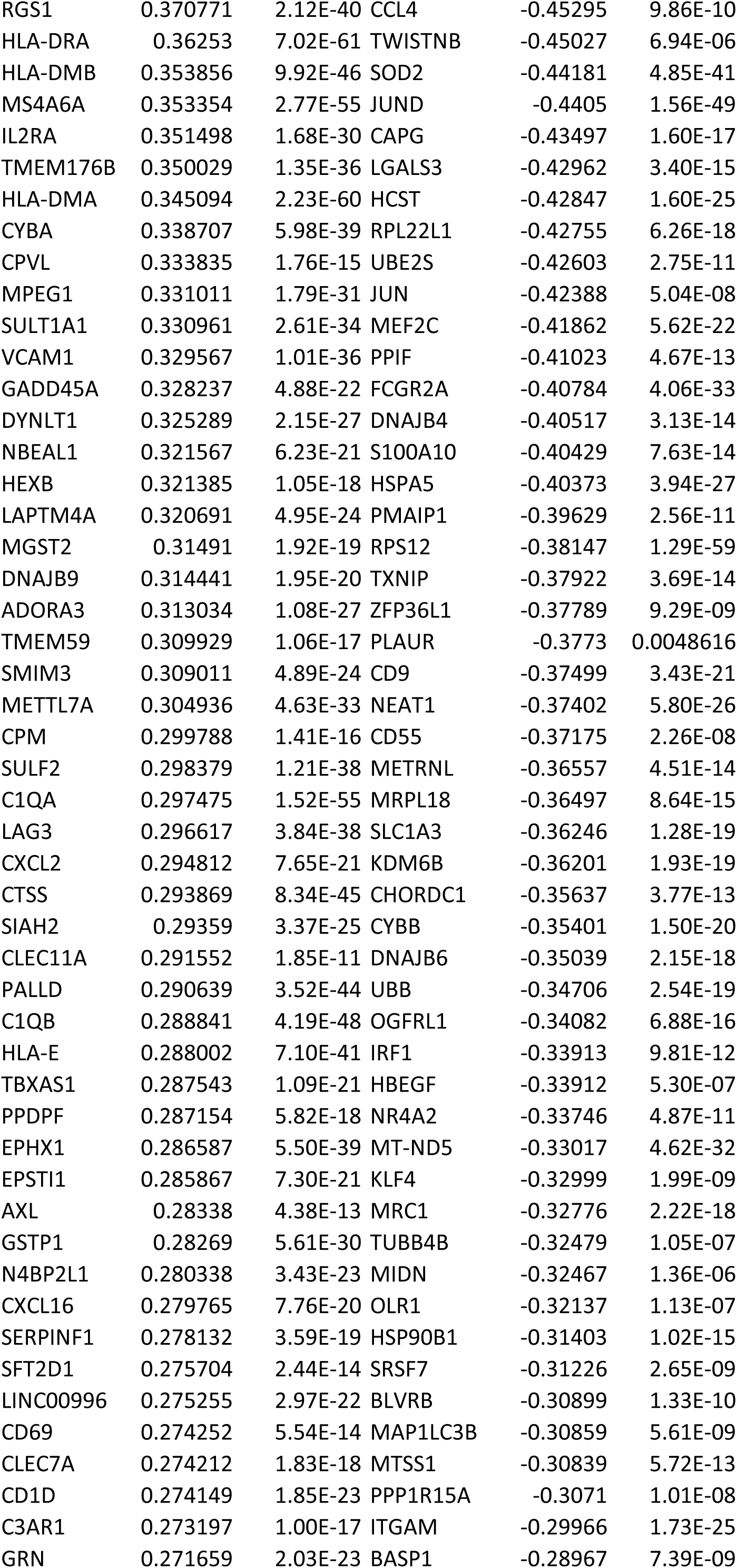
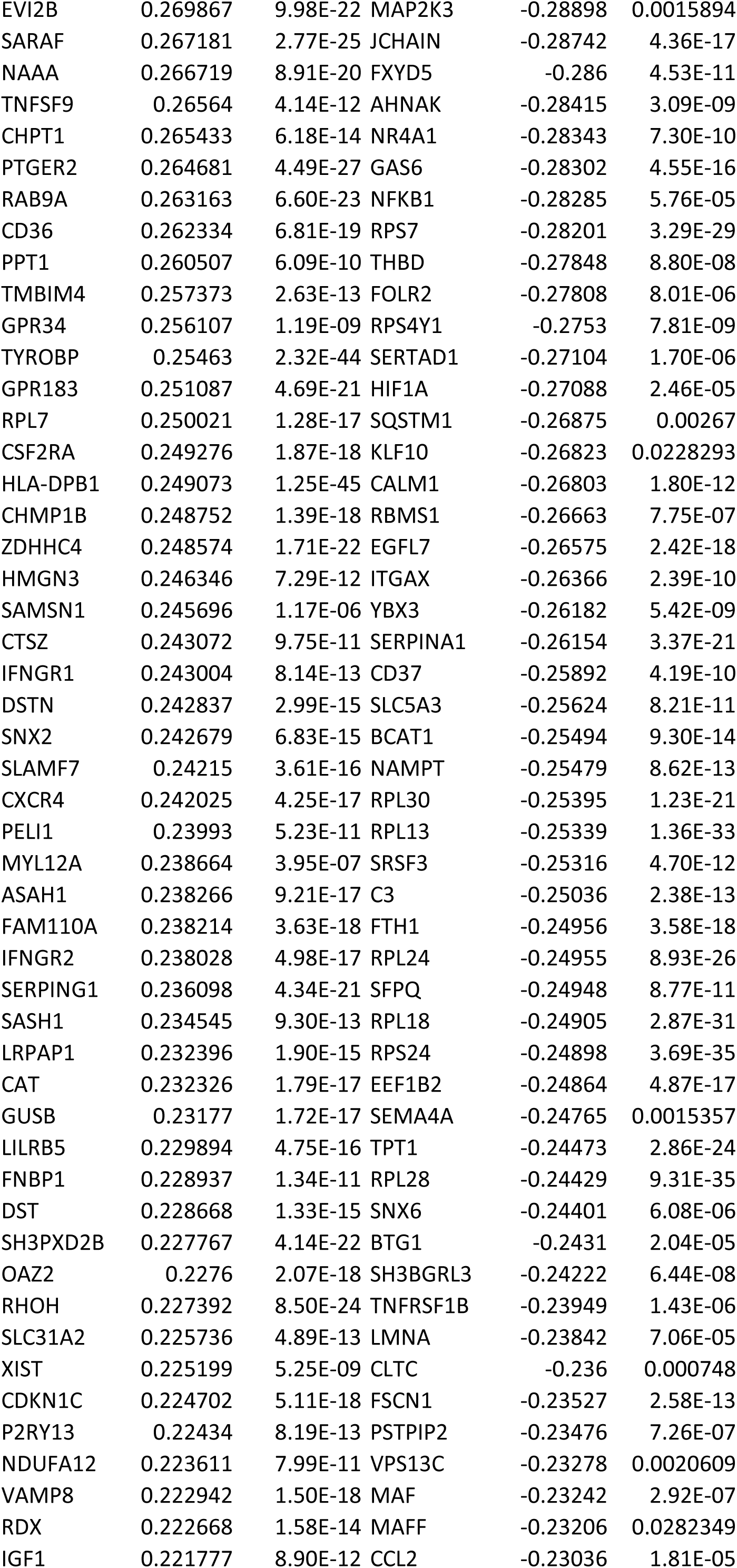
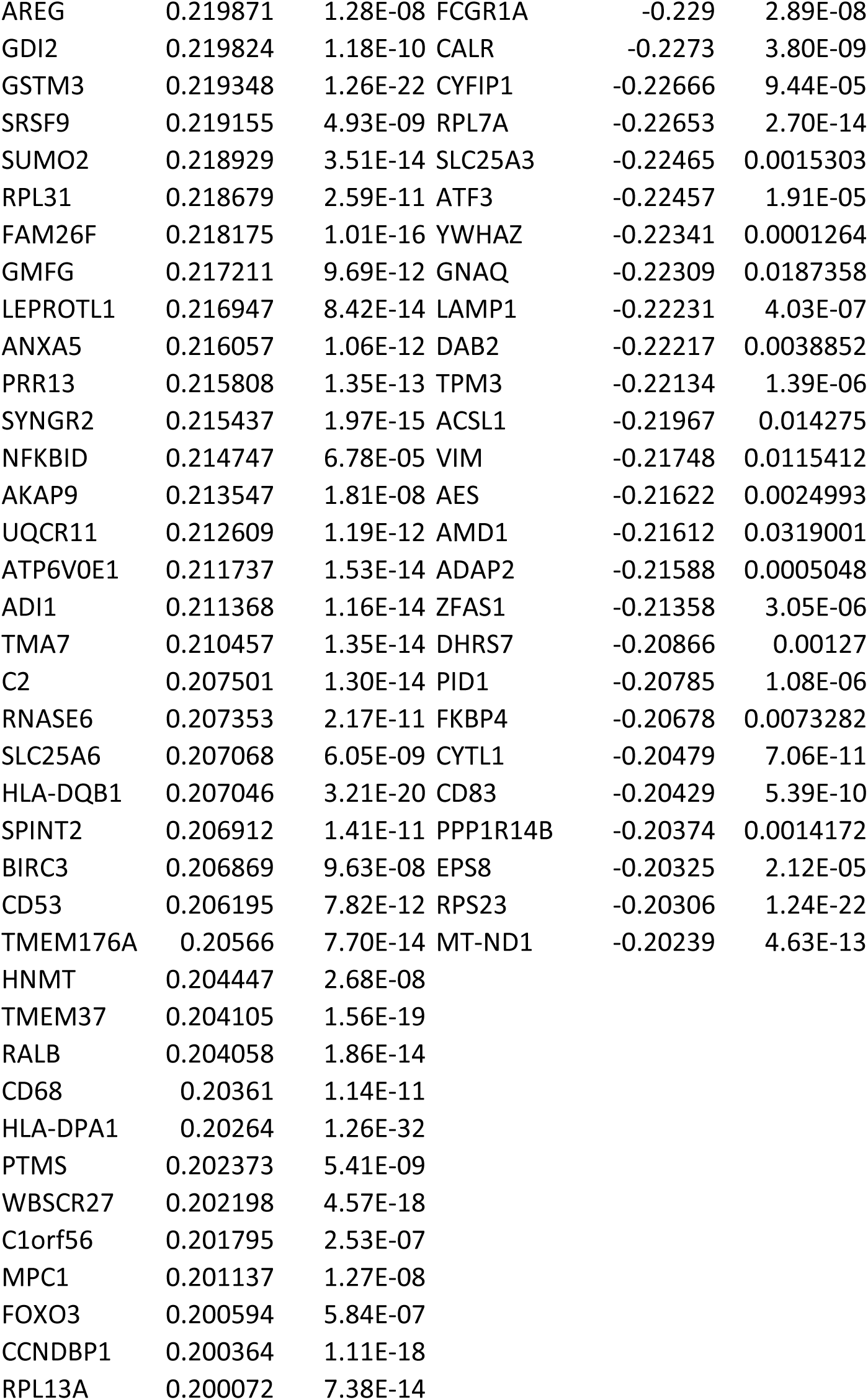
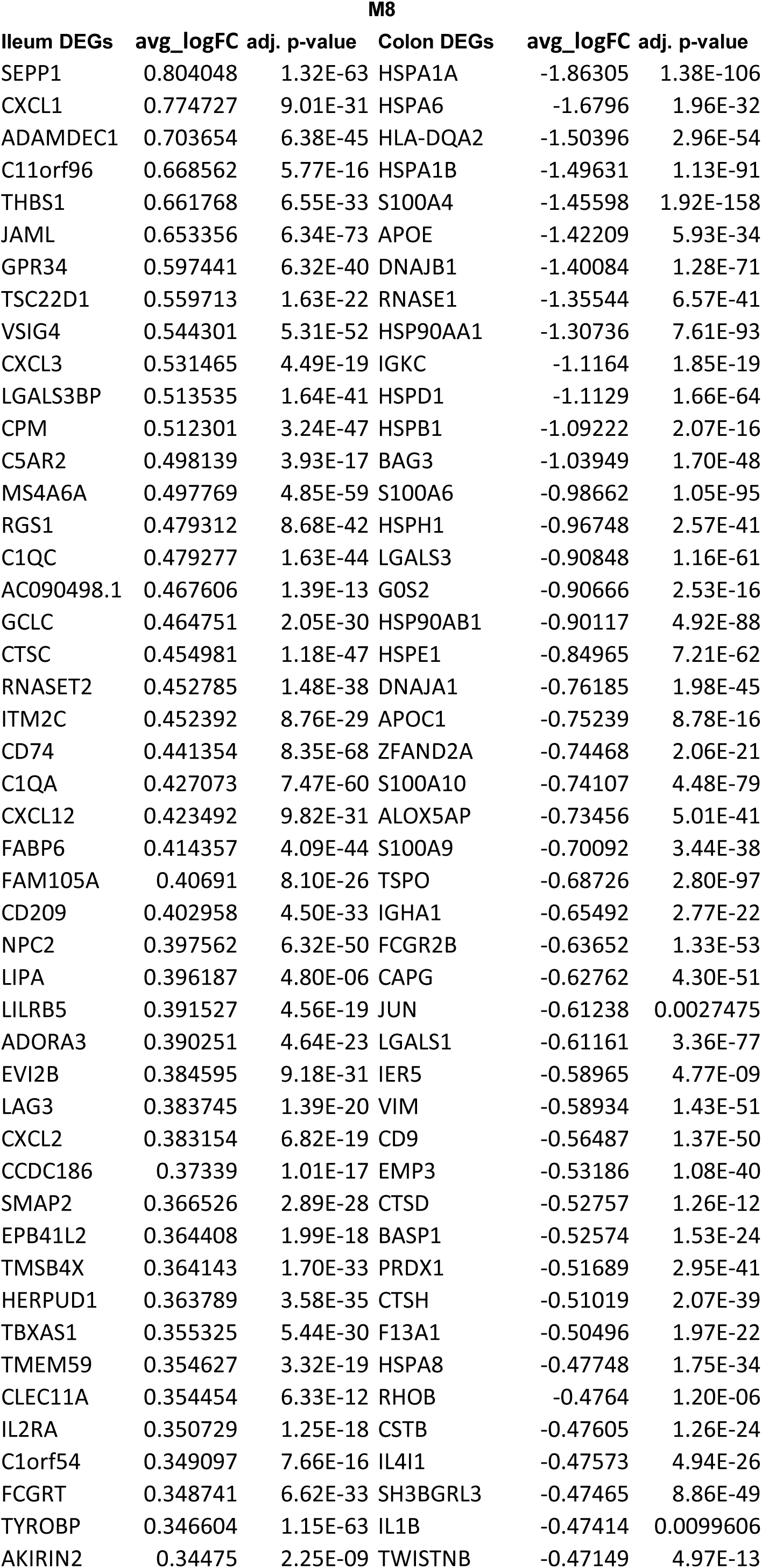
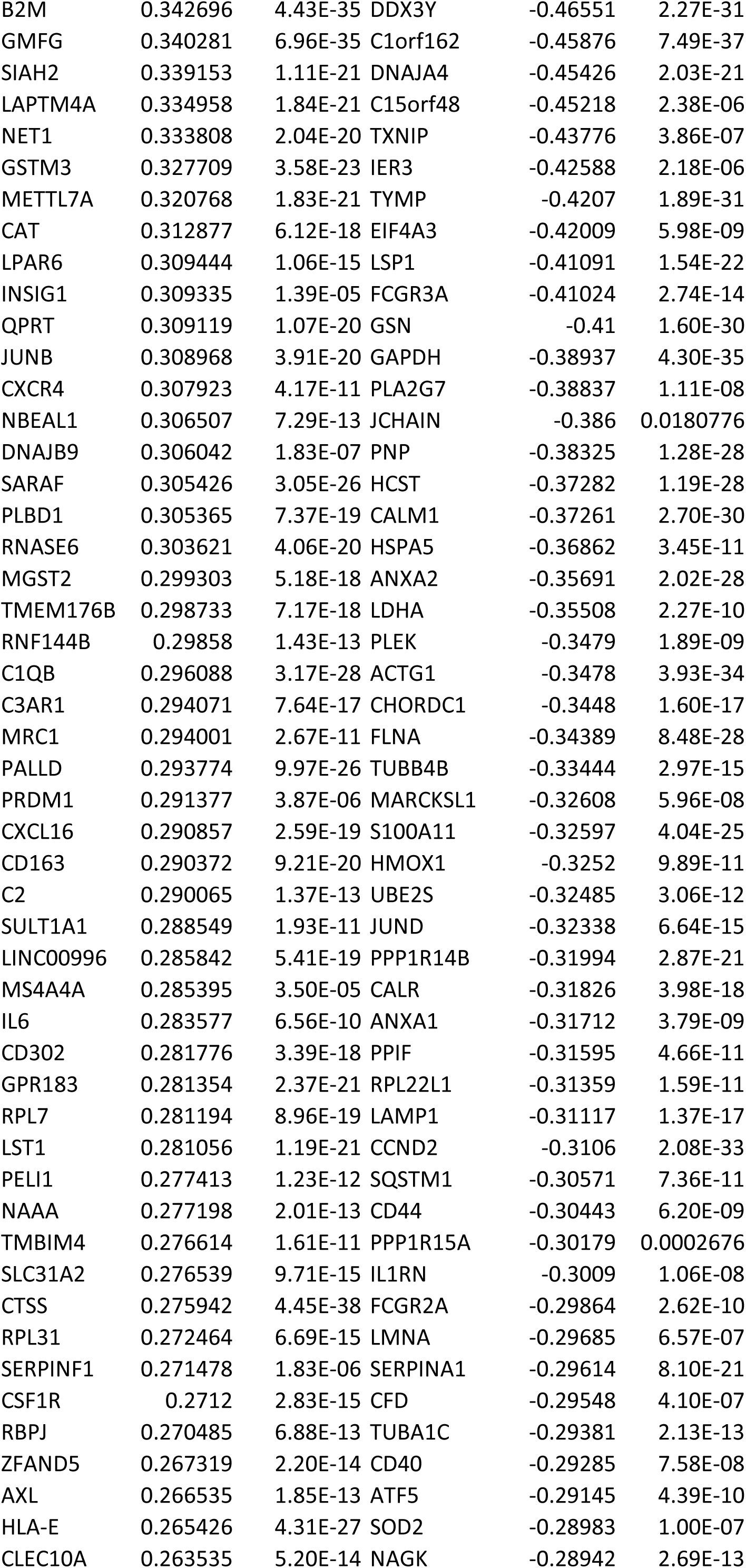
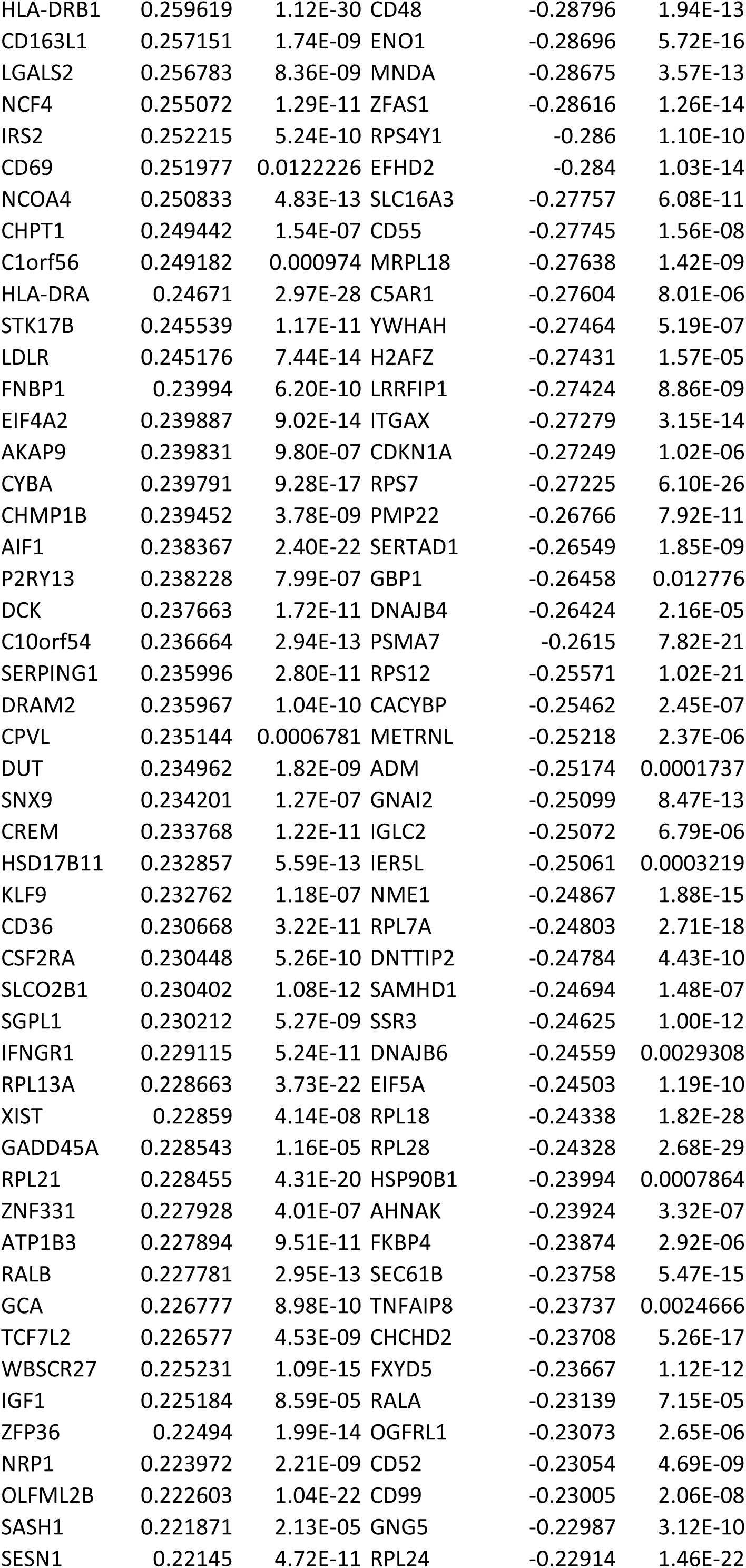
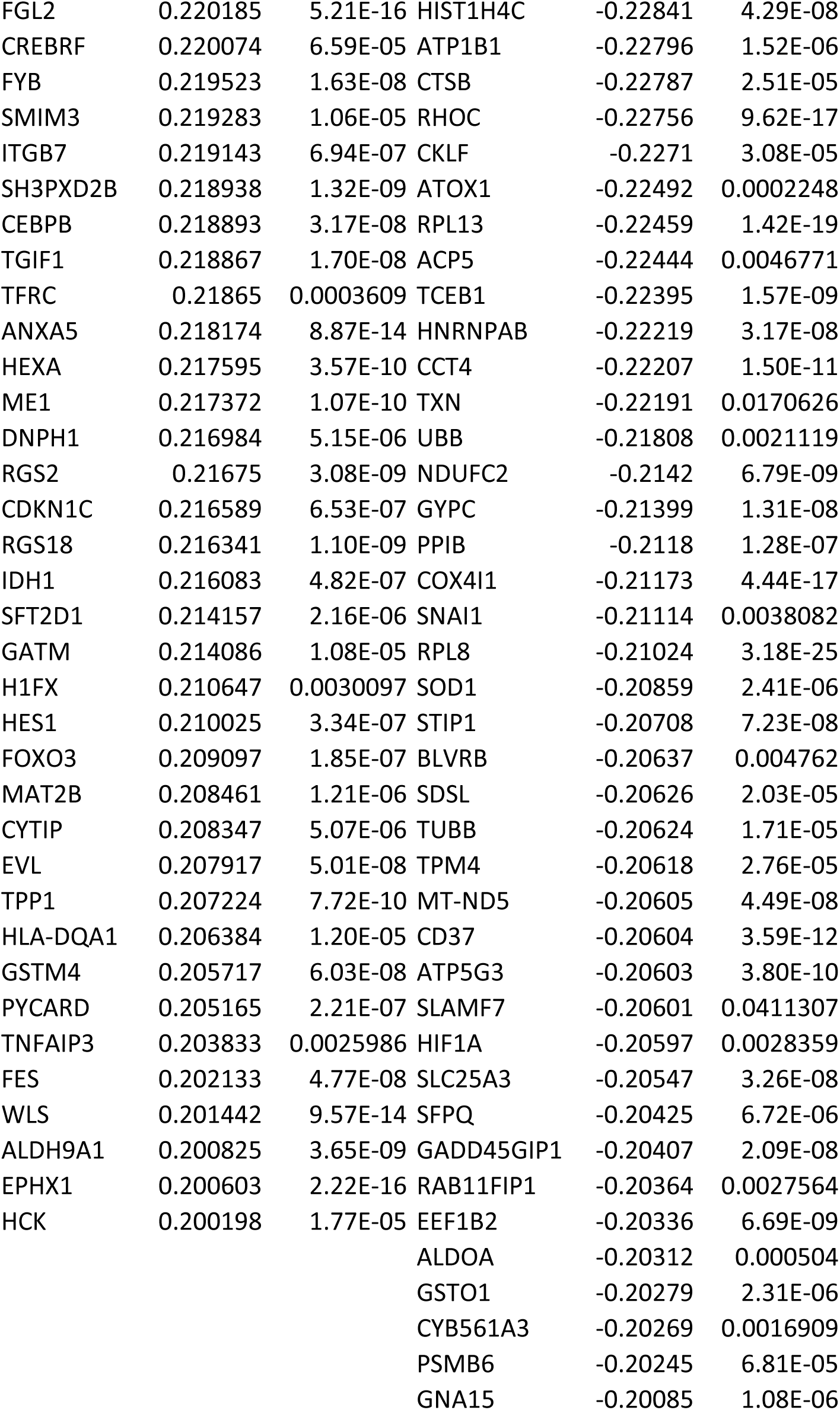
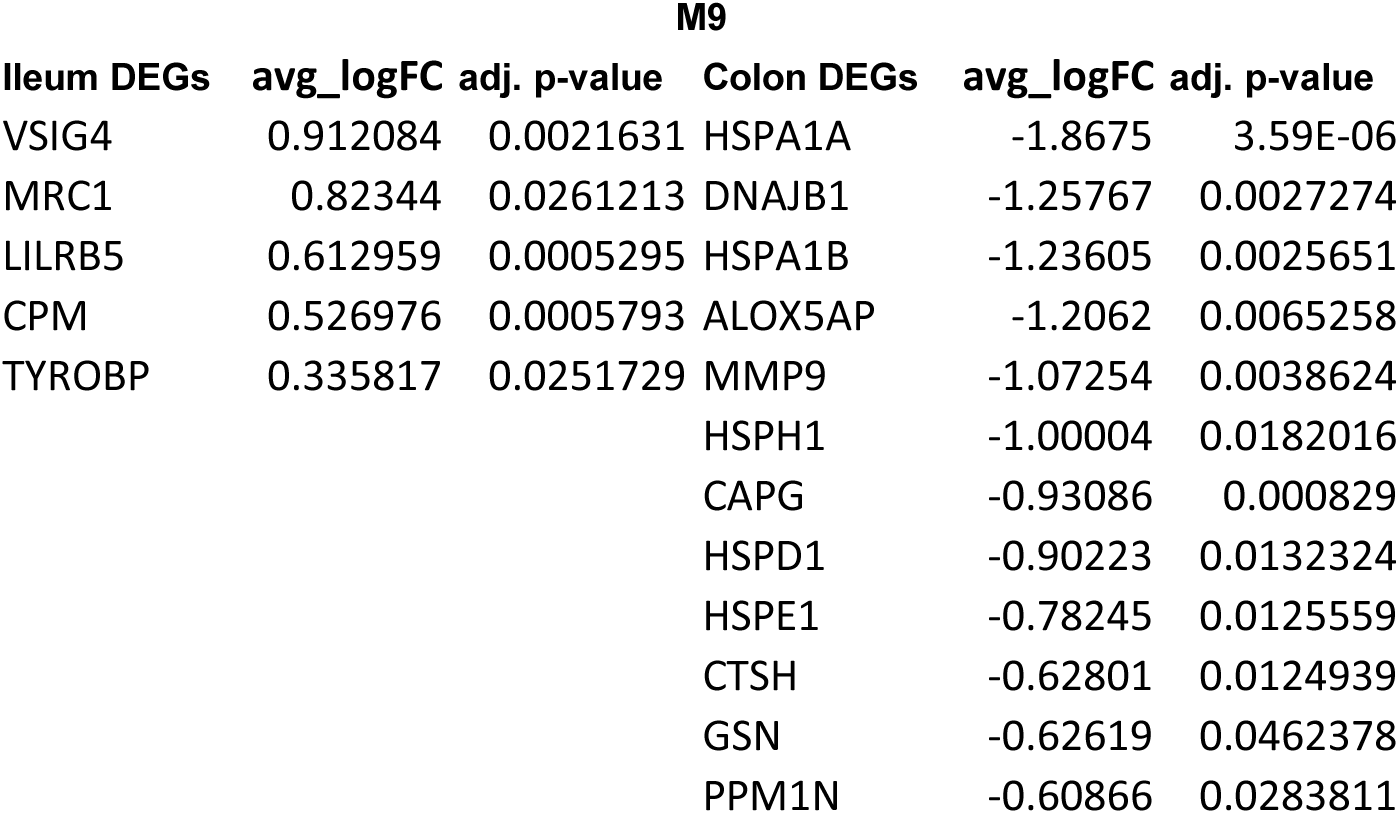
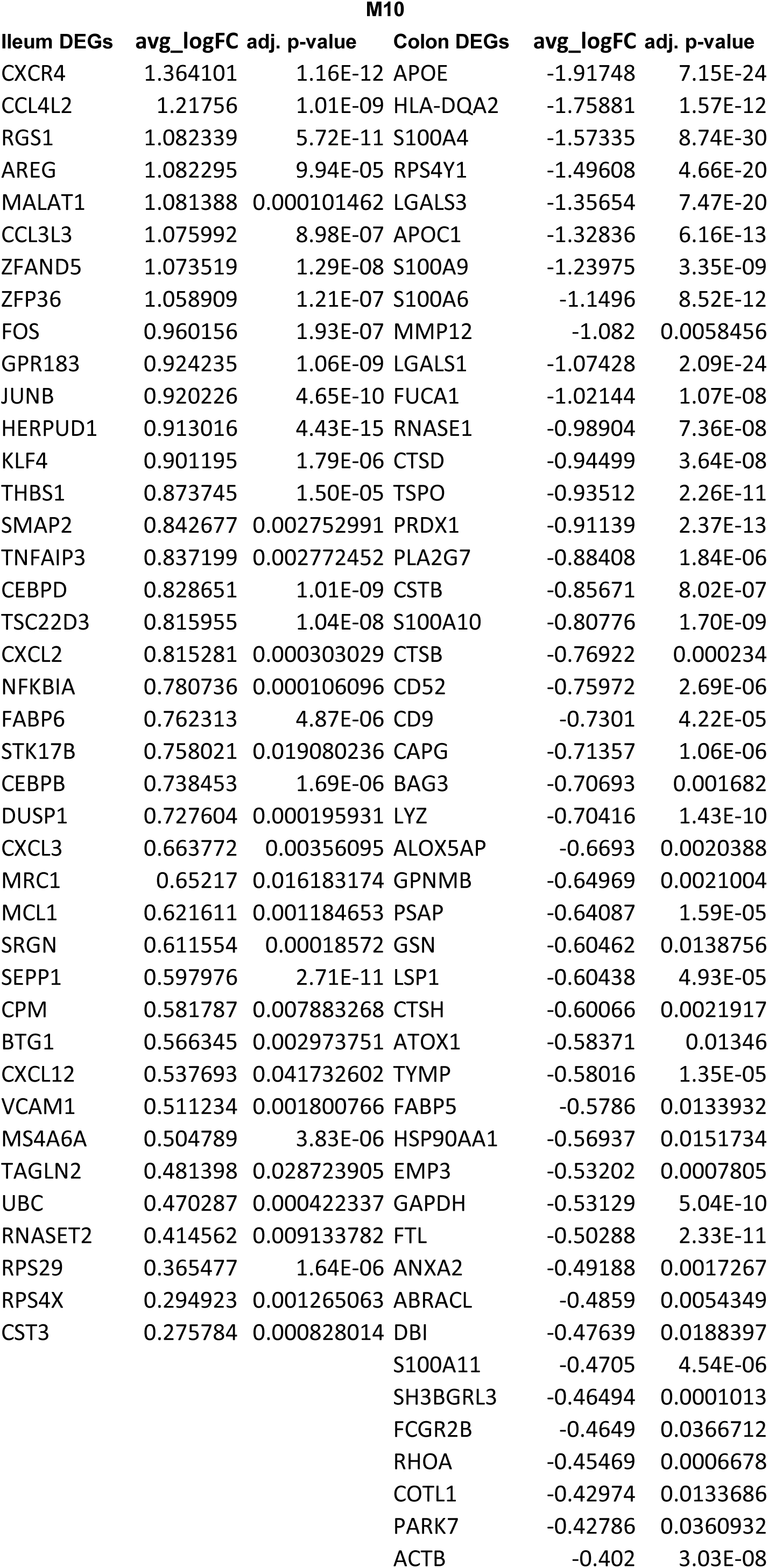
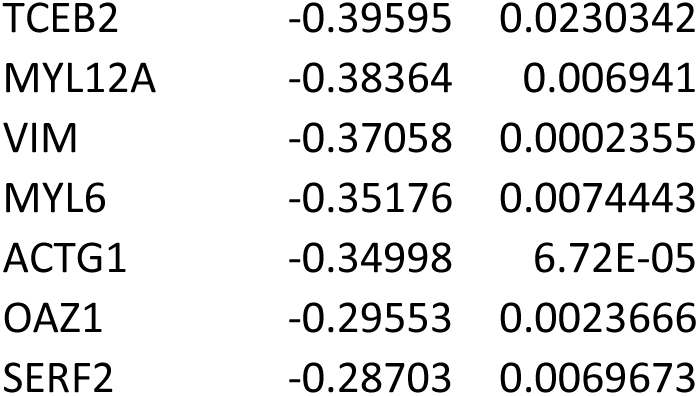
Complete list of DEG comparing ileal and colonic LP mature macrophages for clusters M7-M10.

**Table S6.** Characteristics of anonymized biopsy patient samples used for flow cytometry analysis.

| Patient | Diagnosis | Site | Severity | Vedolizumab |
| --- | --- | --- | --- | --- |
| 1 | CD | Colon | Active- mild | YES |
| 1 | CD | Colon | Quiescent | YES |
| 1 | CD | Ileum | Quiescent | YES |
| 2 | CD | Colon | Active- mod |  |
| 2 | CD | Colon | Quiescent |  |
| 2 | CD | Ileum | Active- mild |  |
| 3 | CD | Colon | Active- mild |  |
| 3 | CD | Colon | Quiescent |  |
| 4 | CD | Colon | Active- mild |  |
| 4 | CD | Colon | Active- mild |  |
| 4 | CD | Colon | Active- mod |  |
| 5 | IBDU (CD) | Colon | Active- mild |  |
| 5 | IBDU (CD) | Colon | Quiescent |  |
| 6 | CD | Colon | Active-mod |  |
| 7 | IBDU (CD) | Colon | Active- mild |  |
| 7 | IBDU (CD) | Colon | Quiescent |  |
| 8 | UC | Colon | Active- mod |  |
| 8 | UC | Colon | Quiescent |  |
| 9 | UC | Colon | Active- mod | YES |
| 9 | UC | Colon | Quiescent | YES |
| 9 | UC | Ileum | Quiescent | YES |
| 10 | CD | Colon | Quiescent |  |
| 10 | CD | Ileum | Active- mild |  |
| 11 | IBDU (UC) | Colon | Quiescent |  |
| 11 | IBDU (UC) | Ileum | Quiescent |  |
| 12 | UC | Colon | Quiescent |  |
| 13 | UC | Colon | Active- mod |  |
| 13 | UC | Colon | Quiescent |  |
| 14 | Healthy | Colon | Healthy |  |
| 14 | Healthy | Ileum | Healthy |  |
| 15 | Healthy | Colon | Healthy |  |
| 16 | Healthy | Colon | Healthy |  |
| 16 | Healthy | Ileum | Healthy |  |
| 17 | Healthy | Colon | Healthy |  |
| 17 | Healthy | Ileum | Healthy |  |
| 18 | UC | Colon | Active- mild |  |
| 18 | UC | Ileum | Active-mild |  |
| 19 | CD | Colon | Quiescent |  |
| 20 | CD | Colon | Active- mild |  |
| 20 | CD | Ileum | Active- mod |  |
| 21 | UC pan | Colon | Active- mod | YES |
| 21 | UC pan | Ileum | Quiescent | YES |
| 22 | UC pan | Colon | Quiescent |  |
| 22 | UC pan | Ileum | Quiescent |  |
| 23 | Healthy | Colon | Healthy |  |
| 23 | Healthy | Ileum | Healthy |  |
| 24 | Healthy | Colon | Healthy |  |
| 25 | CD | Ileum | Quiescent |  |
| 25 | CD | Ileum | Quiescent |  |
| 26 | UC | Colon | Quiescent | YES |
| 26 | UC | Colon | Quiescent | YES |
| 26 | UC | Rectum | Active- mild | YES |

**Table S7.**
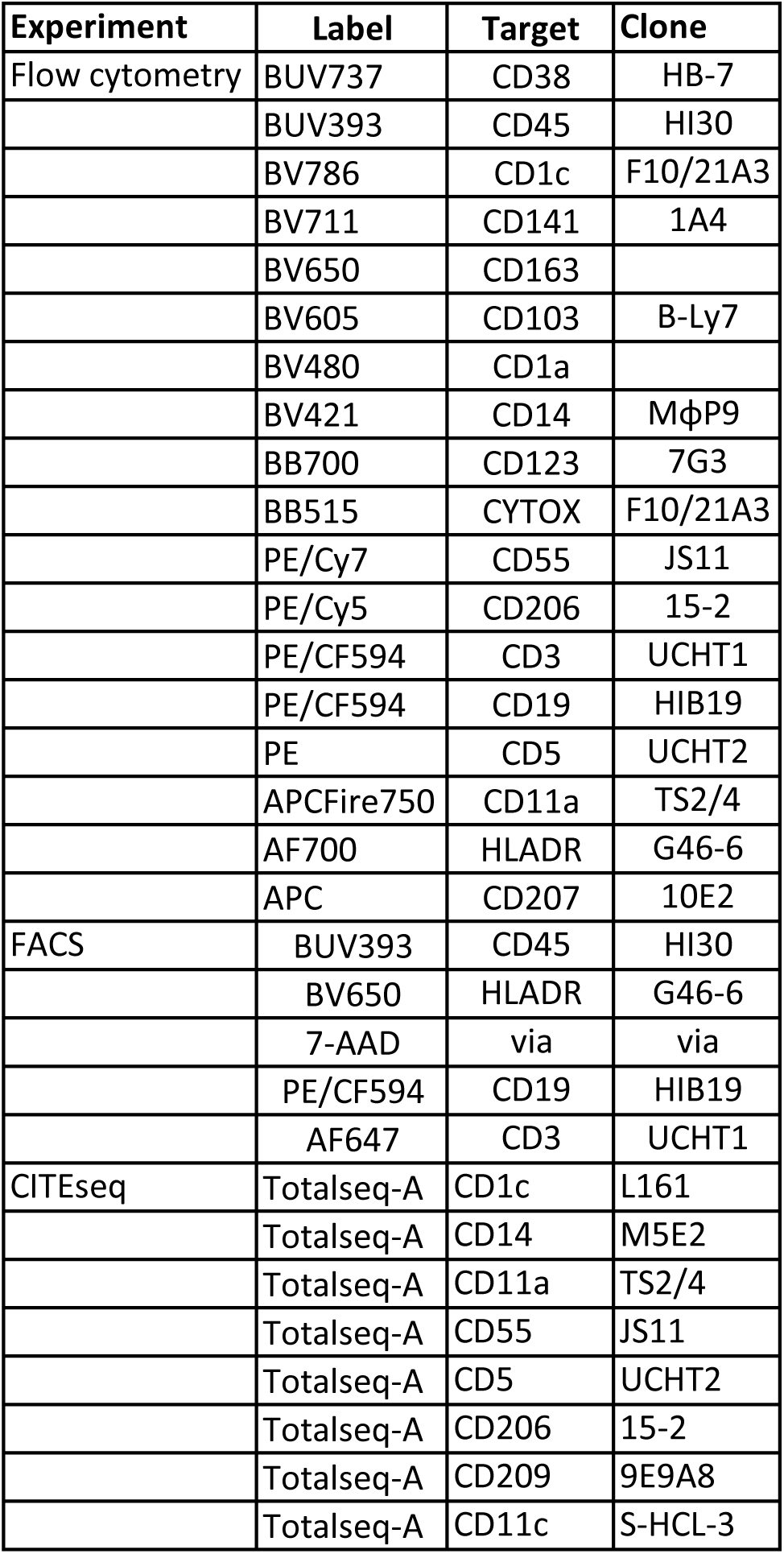
List of antibodies used for flow cytometry, FACS, and CITEseq analysis of mononuclear phagocyte subsets.

| **Experiment** | **Label** | **Target** | **Clone** |
| --- | --- | --- | --- |
| Flow cytometry | BUV737 | CD38 | HB-7 |
|  | BUV393 | CD45 | HI30 |
|  | BV786 | CD1c | F10/21A3 |
|  | BV711 | CD141 | 1A4 |
|  | BV650 | CD163 |  |
|  | BV605 | CD103 | B-Ly7 |
|  | BV480 | CD1a |  |
|  | BV421 | CD14 | MφP9 |
|  | BB700 | CD123 | 7G3 |
|  | BB515 | CYTOX | F10/21A3 |
|  | PE/Cy7 | CD55 | JS11 |
|  | PE/Cy5 | CD206 | 15-2 |
|  | PE/CF594 | CD3 | UCHT1 |
|  | PE/CF594 | CD19 | HIB19 |
|  | PE | CD5 | UCHT2 |
|  | APCFire750 | CD11a | TS2/4 |
|  | AF700 | HLADR | G46-6 |
|  | APC | CD207 | 10E2 |
| FACS | BUV393 | CD45 | HI30 |
|  | BV650 | HLADR | G46-6 |
|  | 7-AAD | via | via |
|  | PE/CF594 | CD19 | HIB19 |
|  | AF647 | CD3 | UCHT1 |
| CITEseq | Totalseq-A | CD1c | L161 |
|  | Totalseq-A | CD14 | M5E2 |
|  | Totalseq-A | CD11a | TS2/4 |
|  | Totalseq-A | CD55 | JS11 |
|  | Totalseq-A | CD5 | UCHT2 |
|  | Totalseq-A | CD206 | 15-2 |
|  | Totalseq-A | CD209 | 9E9A8 |
|  | Totalseq-A | CD11c | S-HCL-3 |

### Supplementary Materials

**Figure S1.**
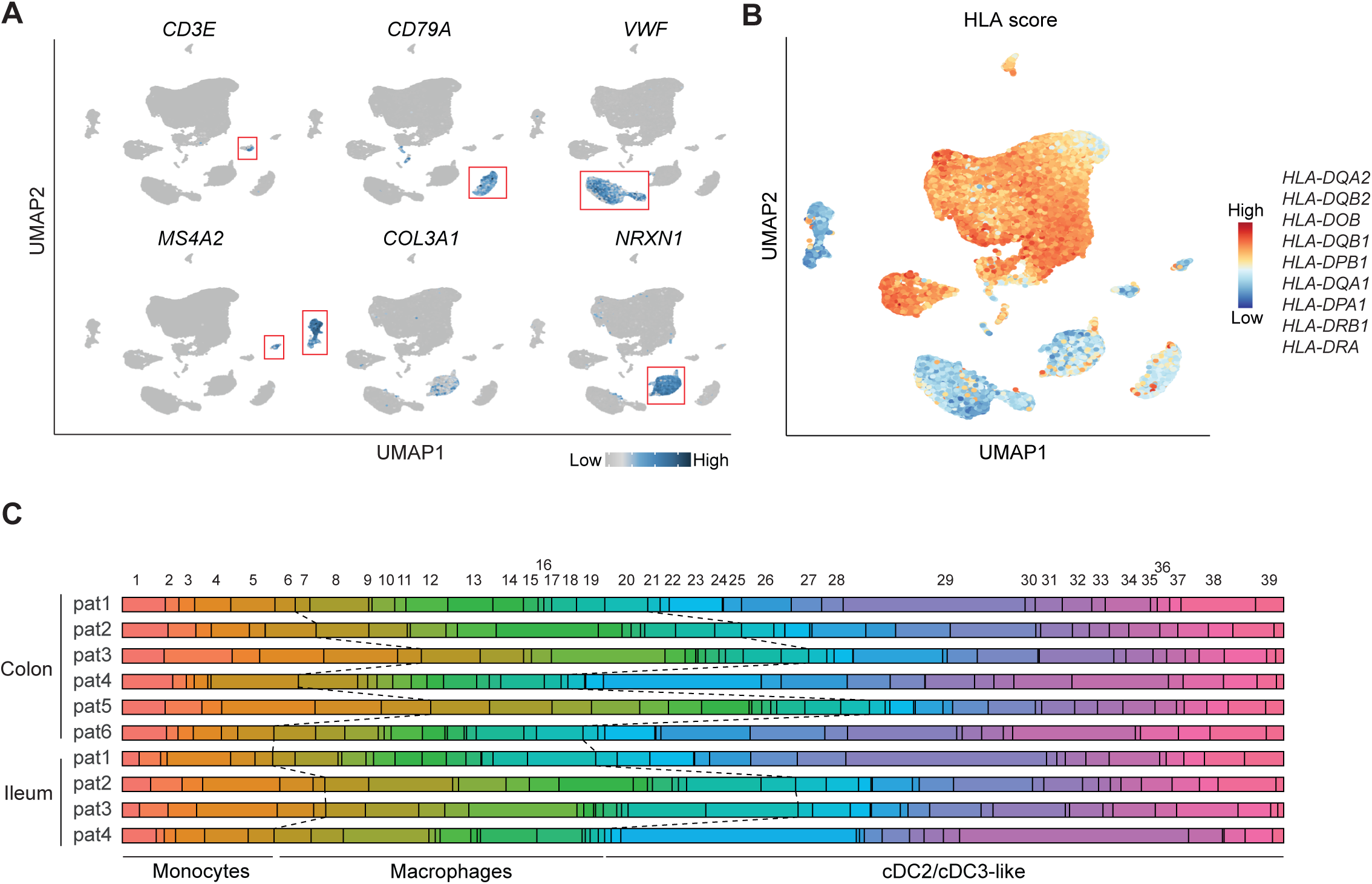
Identification of intestinal LP cell types. (**A** and **B**) UMAP depicting scRNA-seq data of enriched ileal and colonic LP HLA-DR^+^ cells (42,506 cells) isolated using the pipeline depicted in Fig. 1A. (**A**) Examples of signature gene expression used to identify contaminating T cells, B cells, endothelia, mast cells, stromal cells, and glial cells and (**B**) HLA score based on indicated HLA genes. (**C**) Relative abundance of high resolution MNP clusters in ileal and colonic LP samples. Related to Figure 1.

**Figure S2.**
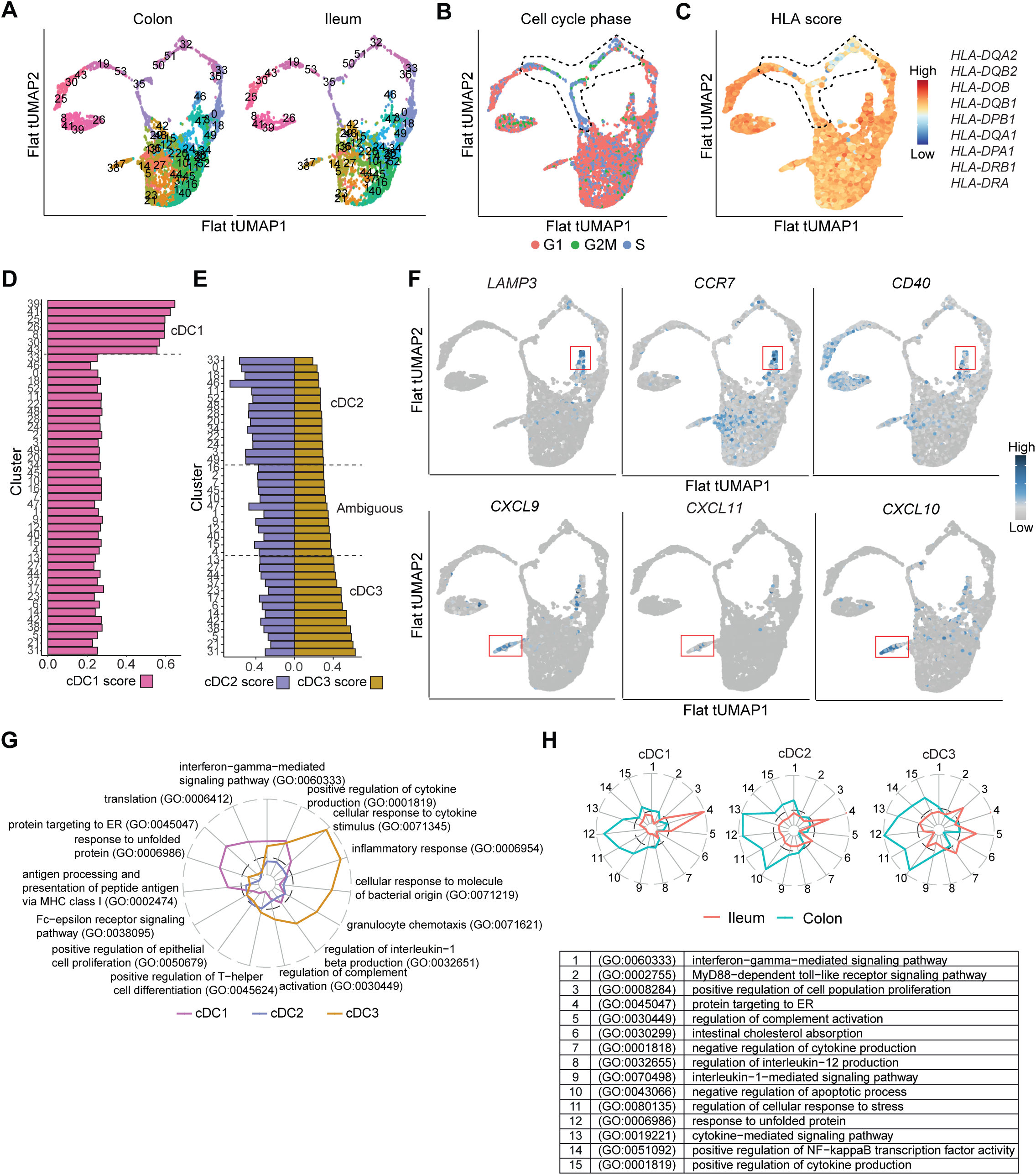
Transcriptional characterization of intestinal LP cDC subsets. **(A)** Flattened 3D tSPACE UMAP (tUMAP) of bioinformatically isolated and re-clustered cDC with high resolution Louvain clustering (52 clusters) and split into ileum and colon LP. (B) Cell cycle profile (**C**) HLA score based on indicated HLA genes. (**B** and **C**) Dashed line represents clusters enriched in cells in G2M/S phase and with low HLA score. (**D**) cDC1 score based on cDC1 signature genes (*CLEC9A, CADM1, XCR1, BATF3,* and *IRF8*) by pseudo-bulk cDC clusters (44 clusters) after removal of the proliferating and HLA^low^ clusters in **B** and **C**. Dashed line indicating cDC1 identity, threshold cDC1 score > 0.4 for cDC1 identity. (**E**) Ranked expression of cDC2 and cDC3 scores by remaining cDC clusters (37 clusters) using signature genes identified by Bourdeley et al^30^. Dashed lines indicating cDC2, cDC3 and ambiguous identities. Clusters were classified as cDC2 when cDC2 score > 0.4 & cDC3 score < 0.3 and as cDC3 when cDC3 score > 0.4. (**F**) tUMAP plots colored by expression of indicated genes (upper panel) associated with cDC maturation and migration migratory marker and (lower panel) interferon inducible genes. (**G** and **H**) Selected biologically relevant terms from GO analysis with EnrichR (Biological Processes 2021). Y-axis = sqrt(-log(adjusted P-value)). Dashed line indicates significance threshold of adjusted P-value = 0.05. (**G**) Based on DEGs from each cDC subset (supplementary table 2) (**H)** Based on DEGs Fig. 2E (supplementary table 4). Related to Figure 2.

**Figure S3.**
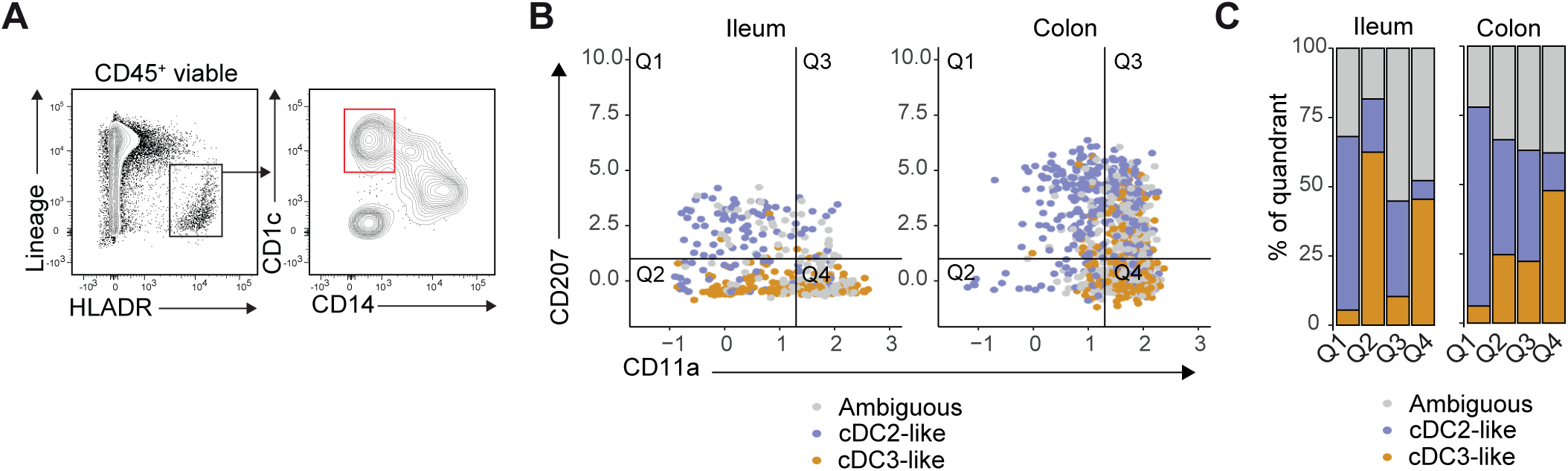
Identification of surface markers that help differentiate intestinal cDC2 and cDC3-like cells. (**A**) Pre-gating strategy to identify colonic-LP CD1c^+^CD14^-^ cDC2, ambiguous and cDC3-like cells. (**B)** CD207 and CD11a expression as assessed by DSB-normalized CITE-seq, Q, quadrant and (**C**) relative proportions of cDC2-like, cDC3-like and ambiguous cDCs within the four CD207 and CD11a CITE-seq quadrants (**B** and **C**) in indicated tissue from a single CRC resection patient. Related to Figure 3.

**Figure S4.**
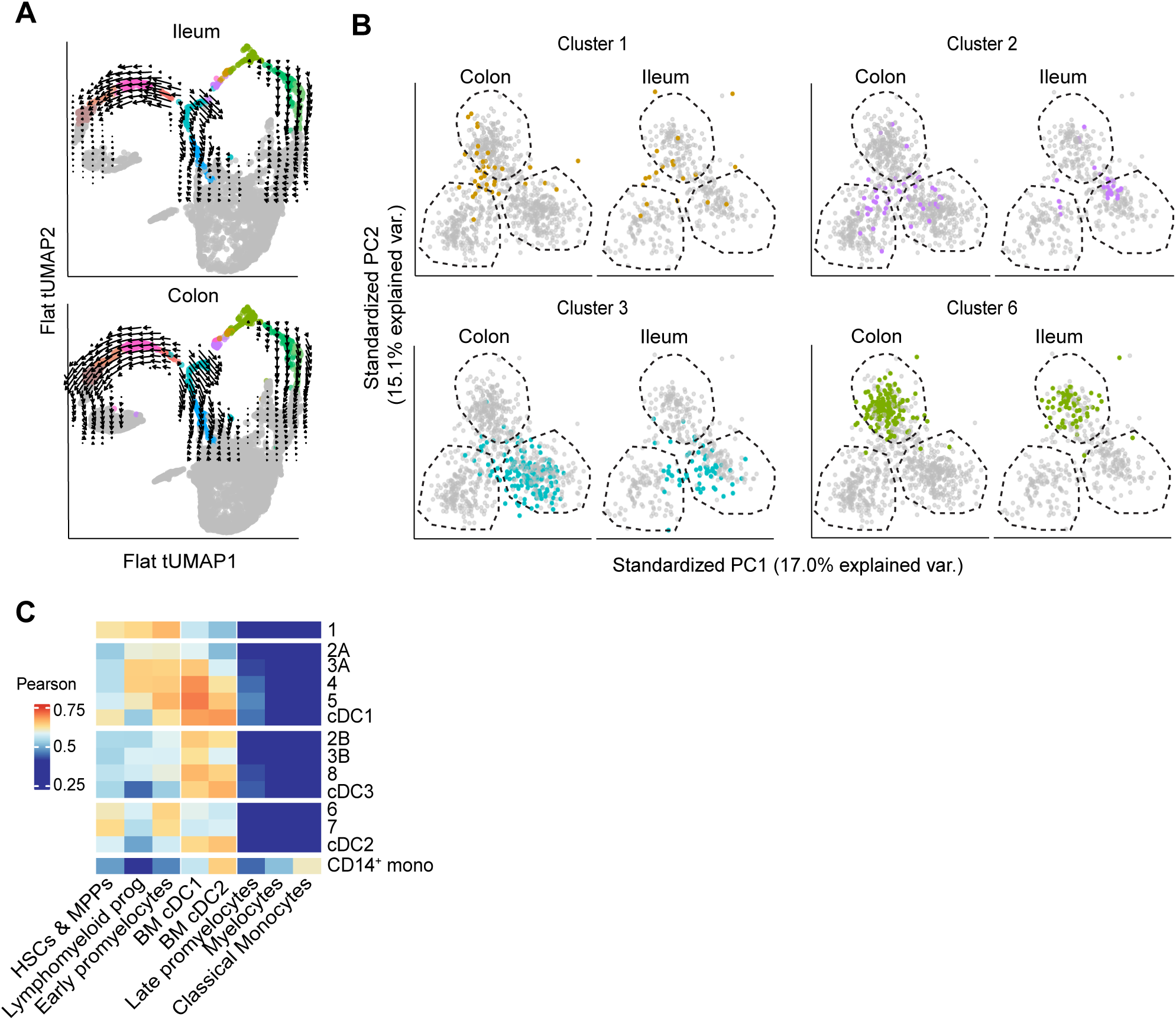
Identification and trajectories of ileum and colon cDC precursor clusters. (**A**) RNA velocities (arrows) of HLA-DR^low^ cDC clusters 3-5 and 7-8 split into ileum and colon derived cDC clusters and calculated with Velocyto package and embedded on Fig. 4A. (**B**) Location of indicated clusters not identifiable using DEG for mature cDC split into ileum and colon (Fig. 4D) on a PCA plot of clusters identified by shared DEG as either pre-cDC1, pre-cDC2 or pre-cDC3 (see Fig. 4F). (C) Heat map depicting Pearson correlation of each intestinal putative pre-cDC cluster and *CD14*^+^ monocytes with indicated progenitor populations from BM described in Triana et al. ^49^. Related to Figure 4.

**Figure S5.**
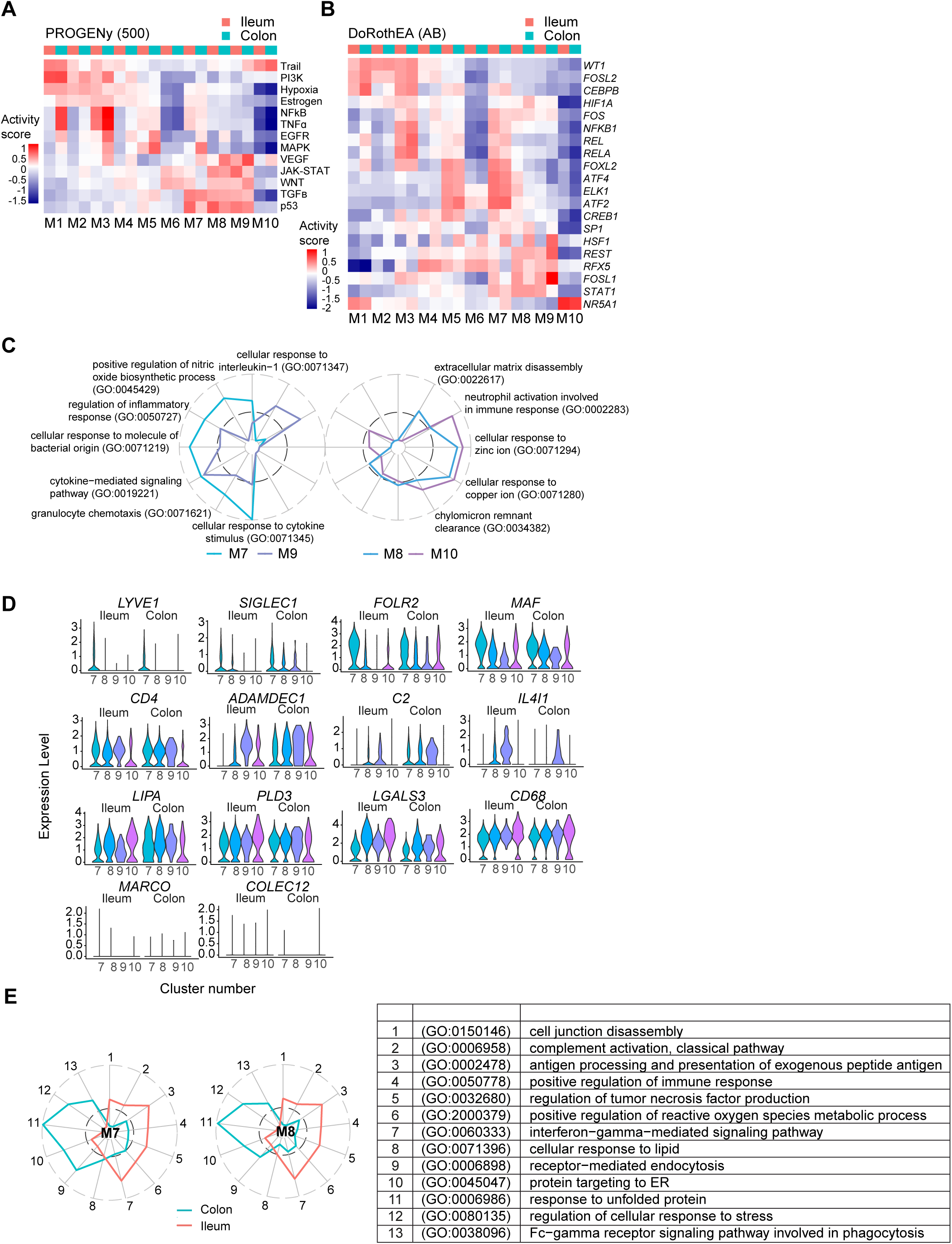
Bioinformatic analysis of intestinal LP macrophage populations. (**A)** PROGENy and (**B**) DoRothEA analysis of indicated clusters from the ileum and colon LP. (**C**) Selected biologically relevant terms from GO analysis with EnrichR (Biological Processes 2021). Y-axis = sqrt(-log(adjusted P-value)). Dashed line indicates significance threshold of adjusted P-value = 0.05. Data are based on DEGs from each mature macrophage subset (supplementary Table 6). (**D**) Violin plots of indicated genes for colonic and ileal LP mature macrophage clusters. (**E**) Significance levels of selected biologically relevant terms from GO analysis with EnrichR (Biological Processes 2021). Y-axis = sqrt(-log(adjusted P-value)). Dark grey dashed line indicates significance threshold of adjusted P-value = 0.05. Data are based on DEGs in Fig. 5H and supplementary Table 8). Related to Figure 5.

**Figure S6.**
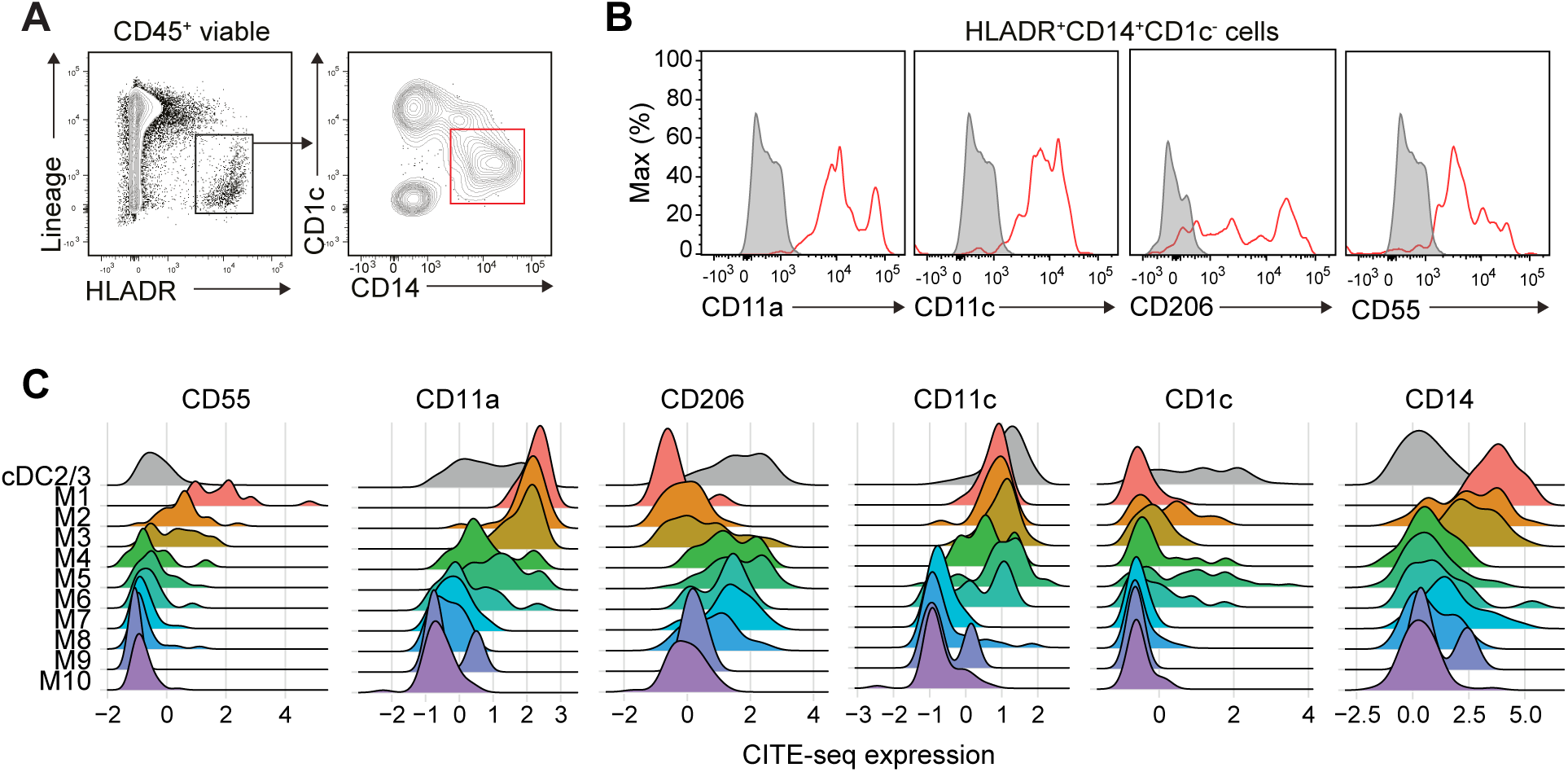
(**A**) Representative flow cytometry analysis showing pre-gating for CD14^+^ MNP. (**B**) Flow cytometry-based expression of indicated markers on CD14^+^ MNP using Legendscreen. Grey fill, FMO. Red line, specific antibody stain. (**C**) DSB-normalized CITE-seq expression of indicated surface markers on ileal LP macrophage clusters after exclusion of the minor proliferating M11 cluster based on n=1. Related to Figure 6.

